# Ultrasensitive small molecule fluorogenic probe for human heparanase

**DOI:** 10.1101/2020.03.26.008730

**Authors:** Jun Liu, Kelton A. Schleyer, Tyrel L. Bryan, Changjian Xie, Gustavo Seabra, Yongmei Xu, Arjun Kafle, Chao Cui, Ying Wang, Kunlun Yin, Benjamin Fetrow, Paul K. P. Henderson, Peter Z. Fatland, Jian Liu, Chenglong Li, Hua Guo, Lina Cui

**Affiliations:** Department of Medicinal Chemistry, College of Pharmacy, University of Florida, Gainesville, FL 32610, USA; Department of Chemistry and Chemical Biology, University of New Mexico, Albuquerque, NM 87131, USA; Division of Chemical Biology and Medicinal Chemistry, Eshelman School of Pharmacy, University of North Carolina, Chapel Hill, NC, USA

## Abstract

Heparanase is a critical enzyme involved in the remodeling of the extracellular matrix (ECM), and its elevated expression has been linked with diseases such as cancer and inflammation. The detection of heparanase enzymatic activity holds tremendous value in the study of the cellular microenvironment, and search of molecular therapeutics targeting heparanase, however, assays developed for this enzyme so far have suffered prohibitive drawbacks. Here we present an ultrasensitive fluorogenic small-molecule probe for heparanase enzymatic activity. The probe exhibits a 756-fold fluorescence turn-on response in the presence of human heparanase, allowing one-step detection of heparanase activity in real-time with a picomolar detection limit. The high sensitivity and robustness of the probe are exemplified in a high-throughput screening assay for heparanase inhibitors.

---

Heparanase, an endo-β-glucuronidase of the glycoside hydrolase 79 (GH79) family^1,2^, is responsible for the cleavage of heparan sulfate (HS) chains of heparan sulfate proteoglycans (HSPG)^3^. These protein-polysaccharide conjugated macromolecules, abundantly expressed in the extracellular matrix (ECM), play an essential structural role in maintaining the ECM integrity. Moreover, the HS side chains bind to an array of biological effector molecules, such as growth factors, chemokines, and cytokines, thereby serving as their reservoir that can liberate the desired signaling molecules when needed.

Therefore the HS-degrading activity of heparanase can lead to a variety of fundamental regulations to impact cellular behavior, such as remodeling and degradation of the ECM, generation of biologically active carbohydrate fragments, and liberation of biological mediators that are bound to HS side chains, thus controlling cellular signaling processes^4,5^. Under normal physiological conditions, heparanase regulates cellular homeostasis, and its high-level expression is mainly observed in the placenta and blood-borne cells such as platelets, neutrophils, mast cells, and lymphocytes^1^. Strikingly, the expression of heparanase is significantly elevated in most types of cancer tissues^6-9^, and increased heparanase activity is regularly linked with increased angiogenesis, metastasis, and shortened post-surgical survival^7,9,10^. The elevated enzymatic activity of heparanase has also been reported in various inflammatory diseases and autoimmune disorders that involve degradation of HS side chains of HSPGs and extensive ECM remodeling^11^.

Despite the existence of other isoforms, heparanase-1 is the only known enzyme for HS cleavage^1^. Heparanase enzymatic activity herein is referred to as the HS-degrading activity of heparanase-1. Furthermore, heparanase is initially expressed as a latent enzyme (proheparanase), which is processed in the lysosome and secreted to the ECM where it meets the substrate HS; therefore, its enzymatic activity cannot be represented by the mRNA level^12^. Methods that can directly measure the enzymatic activity of heparanase are highly desired to facilitate the basic biological study of heparanase in the context of ECM and cellular microenvironment. Such methods are also indispensable in the medical examination of various physiopathological conditions associated with heparanase activity, and in search of molecular therapeutics targeting heparanase or its regulatory molecules.

However, the progress on the development of heparanase probes is disproportionate to the biological importance of the enzyme. Several heparanase assays have emerged over the past two decades, but these assays have encountered disadvantages that limit their broad application^1,13^. Radioisotope-based assays are the most commonly used method for the detection of heparanase activity; these assays measure the radioactivity using radiolabeled heparan sulfate polysaccharides, and require a chromatographic separation of the reaction products, restricting their use to very small sample sizes^14,15^. A more recent homogeneous time-resolved fluorescence assay, currently considered the most convenient approach available to detect heparanase activity^16,17^, requires tedious multi-step reagent addition and the output signal and substrate degradation lacks linear correlation^18^. Another common drawback of these assays developed so far is the use of heterogeneous heparan sulfate polysaccharides, suffering from difficulties in the standardization of the assays due to the structural complexity and diversity of the polymer substrates^13,19,20^. To avoid the disadvantages resulting from heterogeneity, a colorimetric assay^21^ using a pentasaccharide fondaparinux, a commonly used anticoagulant, instead of heparan sulfate as the substrate^22^, but this assay has not gained broad application due to its low sensitivity and requirement of high substrate concentration, high temperature, and long incubation time.

Herein, we describe a fluorogenic small molecule with a defined chemical structure that offers 756-fold fluorescence turn-on response in the presence of human heparanase, allowing one-step detection of heparanase activity in real-time with a picomolar detection limit. Using a standard fluorescence microplate reader, we demonstrate the high sensitivity, simplicity, and robustness of the probe through its application in a high-throughput screening assay for heparanase inhibitors.

## Results

### Structurally defined fluorogenic heparanase probes

The challenge in the development of structurally defined heparanase probes with fluorescence readout largely stems from its elusive substrate recognition mode, despite the extensive investigations being made^23-25^. One known fact is that heparanase is an endoglycosidase that can cleave the internal glycosidic bond between a glucuronic acid (GlcUA) residue and an *N*-sulfoglucosamine (GlcN(NS)) residue bearing either a 3-*O*-sulfo or a 6-*O*-sulfo group^26^. Regarding the substrate size, the minimum heparanase recognition sequence discovered was a trisaccharide [GlcN(NS,6S)-GlcUA-GlcN(NS,6S)]^27^; indeed, molecules containing the disaccharide substrate [GlcN(NS)-GlcUA] have shown mixed results^16,28^. These known features pose a significant hurdle to the development of structurally defined heparanase probes with fluorescence readout. A dye-quencher system can be too bulky for the catalytic site of heparanase, in addition to the synthetic challenge; while a fluorogenic system requires the enzyme to have exoglycosidase properties to cleave the fluorophore off the reducing end (Figure 1). To obtain a fluorogenic probe, can we shift heparanase’s endoglycosidic mode of action to exoglycosidic?

**Fig 1.**
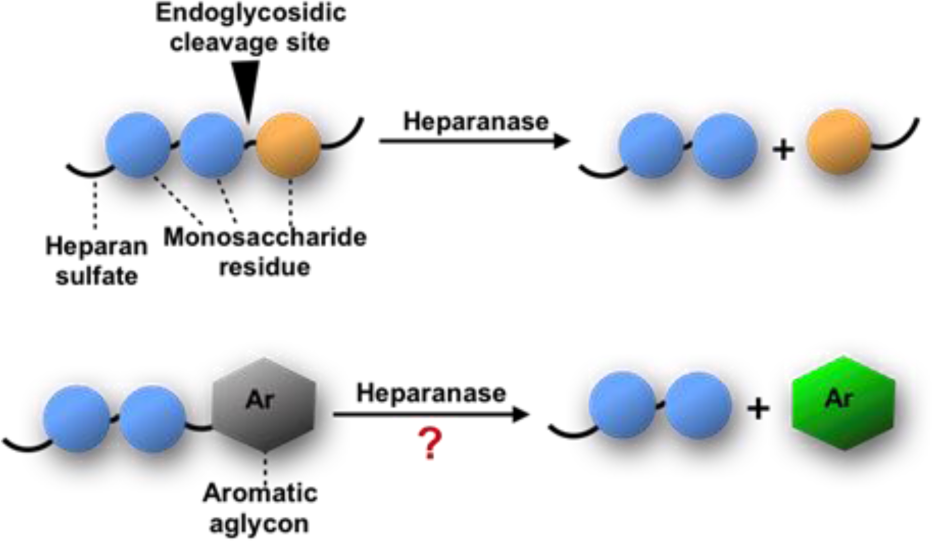
Heparanase cleavage mode.

### Probe response to heparanase can be modulated by electronic effect

To examine this hypothesis, we chemically synthesized compounds **1** to **3** and tested the response of compounds **1** to **3** towards heparanase (HPA)(Fig. 2a, 2b). These compounds are innately non-fluorescent and are stable towards solvolysis, but if the enzyme-catalyzed hydrolysis reaction occurs, the aglycon can be released and fluoresce due to restored intramolecular charge transfer (ICT)^29,30^. Our initial results indicated that probe **2**^28^ could not be activated by human heparanase, while compound **3** produced a slight fluorescence signal after a long incubation time. Compared with the structure of **2**, the marginally activatable compound **3** bears an electron-withdrawing difluoromethyl group, ortho to the glycosidic oxygen, presumably leading to the increased leaving group ability^31^ of the aglycon based on the enzymatic mechanism^32,33^. We envisioned that the activation energy barrier of the enzyme-catalyzed hydrolysis reaction could be further lowered by increased electron-withdrawing property of the aglycon group; therefore, we introduced another aglycon fluorophore 6,8-difluoro-7-hydroxy-4-methylcoumarin (**DiFMU**), carrying two electron-withdrawing fluorine atoms ortho to its phenolic hydroxyl group. Consistent with our hypothesis, incubation of the resulting probe **1** with human recombinant heparanase produced dramatic fluorescence enhancement higher than the background signal (Fig. 2b), and we termed compound **1** as the heparanase-activatable, difluorocoumarin-based probe (**HADP**). Additionally, the fluorine substituents of **DiFMU** increase the quantum yield, improve the photostability^34^, further enhancing the sensitivity of **HADP**. A recent docking study^35^ suggested that **4** should have greater affinity for heparanase owing to an additional sulfate group on the 6-*O* position of the glucosamine moiety. We successfully synthesized this compound, but no fluorescent response was observed (Fig. 2b).

**Fig. 2.**
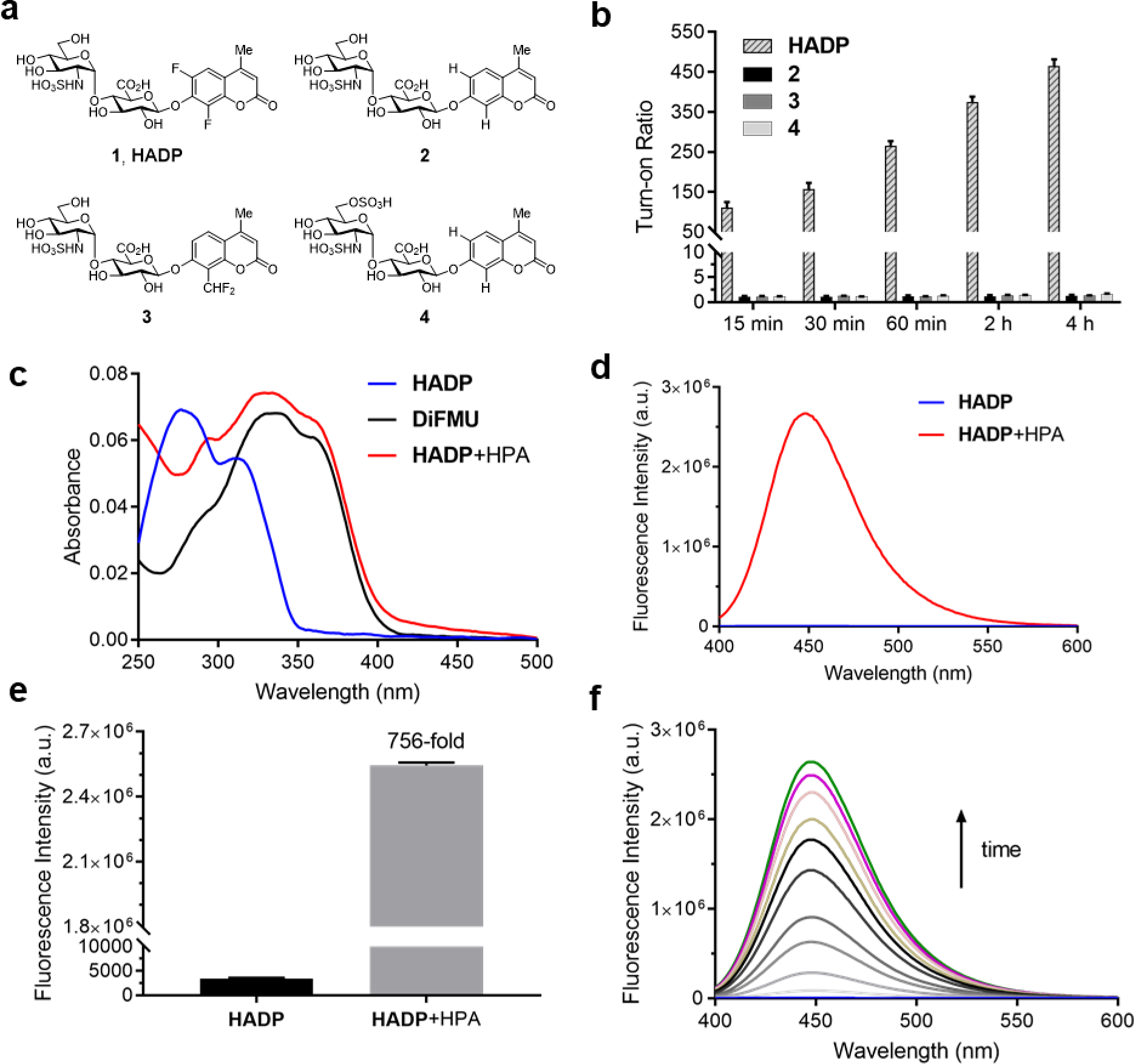
Enzymatic response of synthetic compounds. (**a**) chemical structures of **1** and its analogues **2**-**4**. (**b**) Fluorescence responses of probe **HADP** and control compounds **2**-**4** (each 5 µM) towards heparanase (1 µg) in 40 mM NaOAc buffer (pH 5.0) over time at 37 °C. (**c**) UV-vis absorption and (**d**) fluorescence (λex = 365 nm) spectra of probe **HADP** (5 µM) before (blue line) and after (red line) incubation with heparanase (2 µg) in 40 mM NaOAc buffer (pH 5.0) for 2 h at 37 °C. (**e**) fluorescent quantification of **d**; Turn-on ratio is defined as the ratio of fluorescence intensity after activation to that before activation. (**f**) Time dependence of fluorescence spectra of **HADP** (5 µM) with heparanase (1 µg) from 0 min to 3 h in 40 mM NaOAc buffer (pH 5.0). λex = 365 nm.

### Validation of activation of HADP by recombinant human heparanase

To confirm that the fluorescent response of **HADP** was due to liberation of the fluorophore by heparanase, the UV-vis absorbance and fluorescence emission profiles of **HADP** with or without human heparanase were examined. **HADP** alone displayed maximal absorption peaked at 277 nm and 313 nm (Fig. 2c); Upon incubation with heparanase, a large bathochromic shift was observed with an absorption bump ranging from 320 nm to 365 nm, consistent with the absorption of uncaged **DiFMU** (pH 5.0, NaOAc buffer). **HADP** should be nonfluorescent (Fig. 2d) because the hydroxyl group of **DiFMU** is caged with a disaccharide, resulting in suppressed intramolecular charge transfer (ICT). Upon addition of heparanase, enzyme-triggered cleavage of the glycosidic bond exposes the free hydroxyl group of **DiFMU**, which serves as a strong electron donor in the coumarin D-π-A system, thereby recovering the ICT process and restoring the fluorescence^30^. Consistent with this mechanism, **HADP** incubated with heparanase (2 µg) displayed a remarkable fluorescence enhancement of up to 756-fold (may vary due to different instrumentation setting) at 450 nm within 2 hours (Fig. 2e). Then the time dependence of fluorescence spectra of **HADP** (5 µM) with heparanase (1 µg) were investigated (Fig. 1f, Supplementary Fig. 1). The fluorescence increased dramatically in the initial stage and reached a plateau at 2.5 h. More importantly, the fluorescence intensity exhibited a good linearity to time in initial 15 minutes (Supplementary Fig. 1), facilitating the determination of enzyme kinetics.

To further verify this mechanism, we used HPLC equipped with a diode array detector to monitor the reactions (Supplementary Fig. 2a, 2b). After **HADP** (5 µM) was incubated with heparanase (1 µg) for 4 hours (red line), we observed a clean conversion of the probe to a new peak at 13.1 min, inconsistency with the retention time of **HADP** (black line); the product also shared the same absorption as that of **DiFMU** (Supplementary Fig. 2a inset), suggesting the complete activation of **HADP** by heparanase. Analysis of the new peak by high-resolution electrospray ionization (HR-ESI) mass spectrometry reported two signals at m/z 213.0359 and 235.0181 for the protonated and sodiated adducts of **DiFMU**, respectively (Supplementary Fig. 3). Collectively, heparanase enzymatic activity cleaves the glycosidic linkage of **HADP** to liberate the fluorescent **DiFMU**. No such conversion was observed in the analysis of compounds **2, 3** and **4** treated with heparanase (Supplementary Fig. 2b).

### Computational study of probe activation

We further studied the varied reactivity of compounds **1-3** with heparanase via density functional theory (DFT) and found that the length of the glycosidic bond of compound **1** (**HADP**) was slightly longer than that of compound **2** or **3**, and the energy barrier for compound **1** (**HADP**) (18.67 kcal/mol) in transition state is lower than that of compound **2** or **3** (**Fig. 3**. Supplementary Figs. 4-7). Docking study of the probes with the human heparanase (PDB id: 5E8M) revealed little differences in the binding mode – all three molecules bind to the enzyme in nearly identical mode, and the substituent groups on the coumarin residue in compounds **1** and **3** did not introduce additional binding energy (Figure 3, Supplementary Fig. 8-10, Supplementary Table 1, Supplementary movies 1-3). Collectively, the much higher reactivity of **1** (**HADP**) than **2** or **3** resulted from inherent ground state destabilization^36^ and transition state stabilization.

**Fig. 3.**
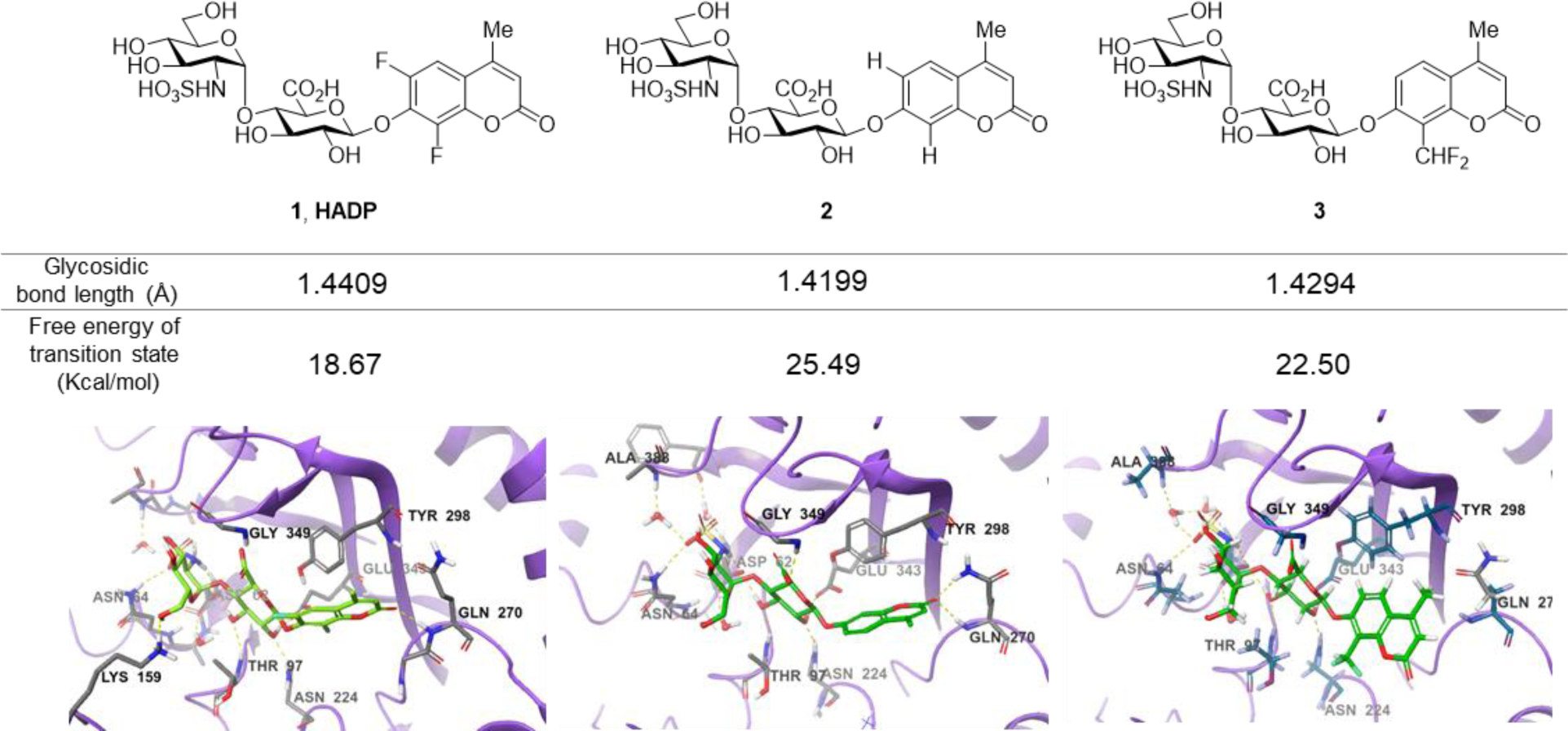
Calculated glycosidic bond length of compounds 1 (HADP), 2 and 3, free energy of transition state and binding modes in heparanase.

### HADP is highly selective for heparanase activity

Human heparanase has the highest enzymatic activity under acidic conditions (pH ∼ 5.4 – 6.8)^37^. We examined the effects of reaction buffer and pH on the activation of **HADP** by heparanase (Supplementary Fig. 11). The reported working buffers for heparanase include 50 mM 2-(*N*-morpholino)ethanesulfonic acid (MES) buffer (pH 6.0)^26^ and 40 mM sodium acetate (NaOAc) buffer (pH 5.0)^2,28^. To address the pH difference, we also tested 40 mM NaOAc (pH 6.0) buffer. At pH 6.0, greater probe activation was detected in the NaOAc buffer than what was observed in the MES buffer, and the reactivity increased further in NaOAc buffer when the pH was reduced to 5.0 (consistent with previous findings^14^).

Aware that pH also tremendously affects fluorescence intensity^38^, we constructed pH profiles of **HADP** and the fluorophore **DiFMU** to determine the sensitivity of the probe at the optimal pH for heparanase activity (Supplementary Fig. 12). There was no remarkable fluorescence change at 455 nm for **HADP** in the pH ranging from 1 to 11 upon excitation, but the free fluorophore **DiFMU** demonstrated significantly greater fluorescence intensity at pH values above its reported p*K*a of 4.7^34^. This indicates that **HADP** is inherently well-suited for heparanase activity assays, as the liberated fluorophore will generate strong signal at the ideal assay pH of 5.0. In contrast, the 4-methylumbelliferone fluorophore of compounds **2** and **4**, with a p*K*a of 7.8, would generate markedly low fluorescence enhancement at this pH and would require a buffering step to optimize the assay sensitivity – if it were activated by heparanase at all. The lower p*K*a of **DiFMU** than the pH of assay buffer contributes conveniently to achieve a simple one-step ‘mix-and-go’ assay without additional basification step.

To confirm that **HADP** is selective for heparanase, the probe was evaluated against a series of possible interfering biological molecules and enzymes at excessively high levels (Fig. 4a). Heparanase exhibited the largest “off-on” response, while no fluorescence increase was observed upon addition of any other molecules or enzymes even at superphysiological levels. For instance, **HADP** showed no response to 100 µM H_2_O_2_, 5 mM glutathione (GSH), or 5 mM cysteine (Cys), common biological oxidizing and reducing species in cells. Hyaluronidase and chondroitinase, which are polysaccharide lyases catalyzing the cleavage of glyosidic bonds between *N*-acetyl-D-glucosamine and D-glucuronic acid in hyaluronic acid, and between D-hexosaminyl and D-glucuronic acid in chondroitin, respectively, did not afford obvious fluorescence change. Likewise, β-glucuronidase did not trigger a fluorescence response of **HADP** possessing a glycosyl substituent on the O-4 position of the glucuronic acid residue. Overall, **HADP** demonstrates remarkable selectivity for heparanase over other glycosidases and common biomolecules.

**Fig 4.**
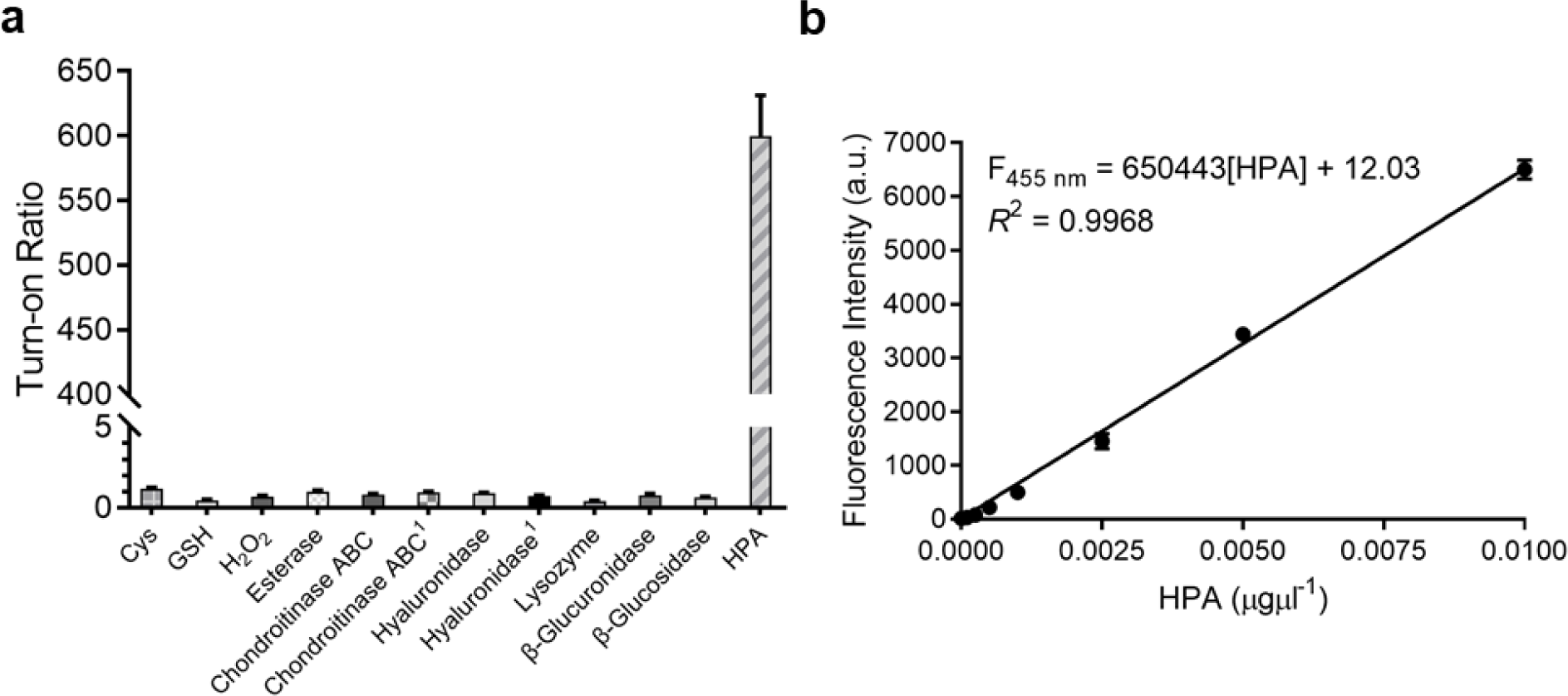
Sensitivity and selectivity of HADP. (**a**) Fluorescence responses of probe **HADP** to Cys (5 mM), GSH (5 mM), H_2_O_2_ (100 µM), Esterase, Chondroitinase ABC, Hyaluronidase, Lysozyme, β-Glucuronidase, β-Glucosidase (5 µg each) and HPA (2 µg) for 4 h. ^1^ 50 µg enzyme was used. (**b**) Linear plot of fluorescence intensity agaist various concentrations of heparanase at 28 min. λex/em = 365 nm/455 nm.

### Detection limit and kinetics of heparanase by HADP

To determine the detection limit of **HADP** against heparanase, concentration-dependent studies over time were performed with 5 µM **HADP** (Supplementary Fig. 13a). A good linear relationship of concentration-fluorescence intensity was obtained in the heparanase concentration range of 0-0.01 µgµL^−1^, with an equation of F_455 nm_ = 650443[HPA] + 12.03 (*R*^2^= 0.9968) (Fig. 4b). Based on the 3σ/s method, the limit of detection (LOD) of **HADP** was calculated to be 0.35 ngmL^-1^ (67 pM), indicating the ultra-sensitivity of **HADP** for heparanase detection.

To test if **HADP** was suitable for rapid and quantifiable heparanase detection, we evaluated the kinetics of the probe activation by heparanase. The kinetic parameters of the enzymatic activation of **HADP** by human heparanase were determined via time-dependent fluorescence intensity measurements in the presence of heparanase and **HADP** at different concentrations (Supplementary Fig. 13c). The Michaelis constant (*K*_M_), catalytic efficiency (*k*_cat_/*K*_M_) and turnover number (*k*_cat_) were calculated to be 8.3 µM, 0.29 µM ^−1^ min^−1^ and 2.4 min^−1^, respectively. Compared with the *K*_M_ of fondaparinux (46 µM)^21^, a pentasaccharide substrate for heparanase, **HADP** exhibited a significantly high affinity for heparanase.

### HADP-assisted high-throughput inhibitor screening

Heparanase has been considered a drug target for cancer and inflammation^39-44^. Nonetheless, only four saccharide-based inhibitors have been assessed in clinical trials^1,45^. The major challenge in screening inhibitors for heparanase is the lack of robust assays^13^. To evaluate the ability of **HADP** for inhibitor screening, we first tested whether it could accurately measure the potency of a known experimental heparanase inhibitor, suramin. Based on our assay, the calculated IC_50_ value of suramin was 1.0 µM (Fig. 5a), in good agreement with that determined by previously mentioned HTRF assay^46^.

**Fig. 5.**
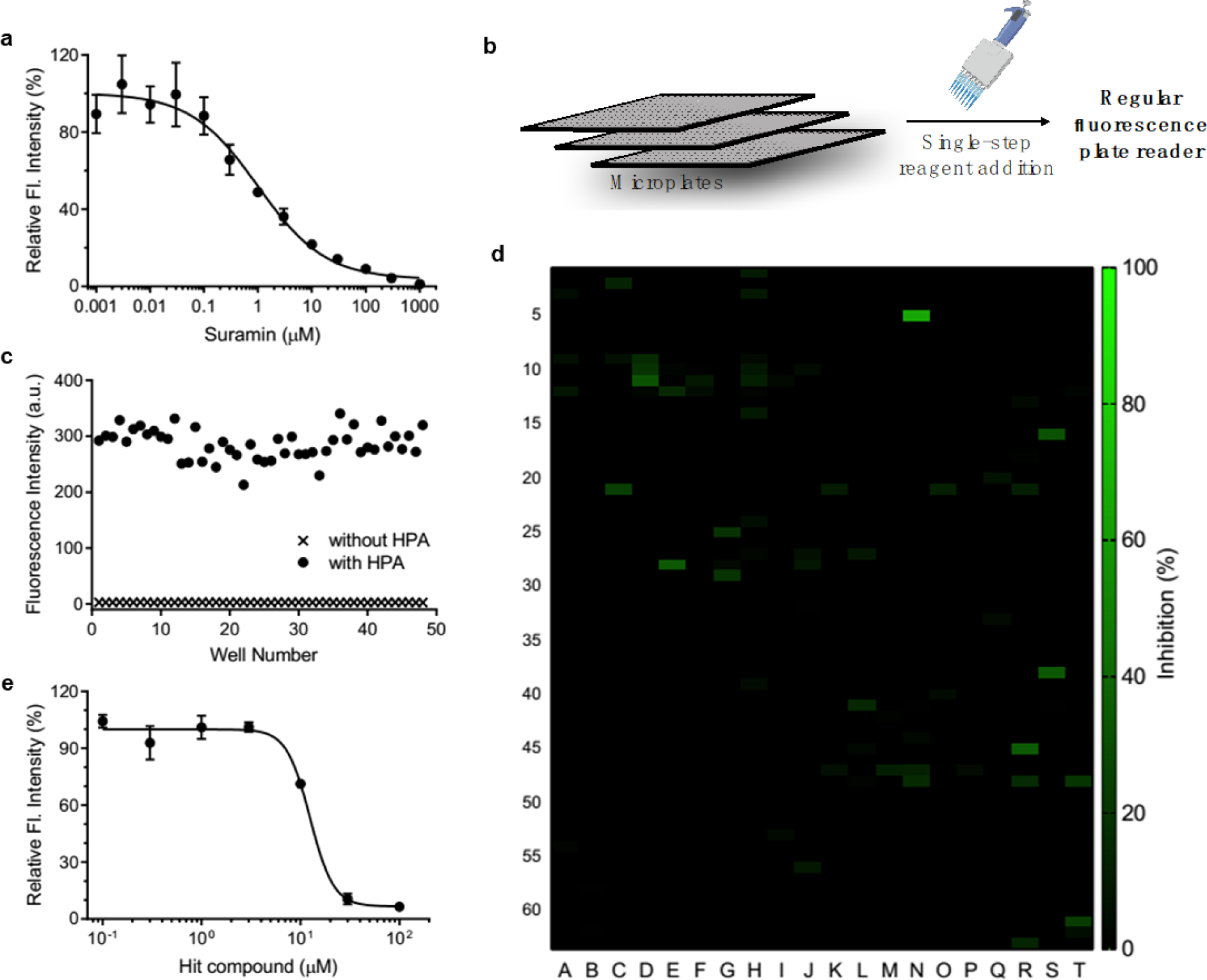
Inhibition evaluation and screening. (**a**) inhibitory activity of suramin. (**b**) schematic of mix-and-go assay for hpa inhibitor screening. (**c**) quality assessment of mix-and-go assay. (**d**) screening results of a commercial library. (**e**) validation of inhibitory activity of hit compound. λex/em = 365 nm/455 nm.

The accuracy of this assay was further evaluated via manual pipette operation to give a Z′-factor of approximately 0.7 (Fig 5c), indicating that the **HADP**-based assay can be an excellent platform for high-throughput drug screening^47^. Next, we accessed the performance of **HADP** in a semi-high-throughput format, screening through a library of 1280 compounds in a 384-well plate assay against heparanase (Fig 5d). Several hits were identified, and further validation via an inhibition assay identified a new drug scaffold with micromolar inhibitory activity toward heparanase (Fig 5e). These results suggest that **HADP** is a sensitive and robust probe for high-throughput screening in seeking potential heparanase inhibitors.

## Summary

In summary, via twisting the heparanase’s endoglycosidic mode of action through modulating the electronic properties of the algycon, we have developed the first structurally defined ultrasensitive fluorogenic probe **HADP** (**1**) for detecting heparanase activity; it offers up to 756-fold fluorescence turn-on response in the presence of human heparanase, allowing one-step detection of heparanase activity in real-time with a picomolar detection limit. We demonstrated the high sensitivity, simplicity, and robustness of **HADP** through high-throughput screening of novel heparanase inhibitors. The probe described will find broad applications in basic cell biology study, the development of diagnostics, and drug discovery.

## Methods

### Synthesis

For detailed synthesis of heparanase probe and its control compounds, see the Supplementary Information. NMR spectra were recorded on a Bruker instrument (500 MHz and 126 MHz, 600 MHz and 150 MHz, 800 MHz and 200 MHz respectively) and internally referenced to the residual solvent signals (^1^H: δ 7.26; ^13^C: δ 77.16 for CDCl_3_, ^1^H: δ 3.31; ^13^C: δ 49.0 for CD_3_OD respectively). NMR chemical shifts (δ) and the coupling constants (J) for ^1^H and ^13^C NMR are reported in parts per million (ppm) and in Hertz, respectively. The following conventions are used for multiplicities: s, singlet; d, doublet; t, triplet; m, multiplet; and dd, doublet of doublet. High resolution mass was recorded on Waters LCT Premier Mass Spectrometer.

### UV-Vis and fluorescence spectrum

To a solution of 20 µL of 50 µM probe **1** (**HADP**) and 160 µL 40 mM NaOAc (pH = 5.0) was added 20 µL 0.1 µg/µL heparanase to obtain 5 µM **1** (**HADP**) with 2 µg heparanase solution. After incubation at 37 °C for 2 h, the reaction solution was transferred to quartz cuvettes to measure absorbance or fluorescence. Absorbance spectrum scan range from 250 nm to 500 nm (1 nm increment). fluorescence spectrum setting: λex = 365 nm, Slit Wicdth 2.5 nm. Emission was record from 400 nm to 600 nm, Slit Width 5 nm. Absorption spectra were recordes on Shimadzu UV-2700 UV-VIS Spectrophotometer. Fluorescence spectra were recorded on Fluorolog TAU-3 Spectrofluorometerwith a xenon lamp (Jobin Yvon-Spex, Instruments S. A., Inc.)

### Fluorescence responses of probe HADP and control compounds 2-4 to heparanase

To a solution of 20 µL of 50 µM each compound and 160 µL 40 mM NaOAc (pH = 5.0) was added 20 µL 0.05 µg/µL heparanase to obtain 5 µM each compound with 1 µg heparanase solution in 96-well microplate. Each compound was performed in triplicates. Then the microplate was sealed with sealing film and incubated at 37 °C. Fluorescence intensity was recorded on Spectramax M5 Multimode plate reader (Molecular Devices, USA) at each timepoint. λex/em = 365 nm/455 nm.

### Fluorescence responses of probe HADP to biological species

5 µM probe **1** (**HADP**) was incubated with various biological species at the indicated amount/concentration in 96-well microplate. Cysteine (5 mM), Glutathione (5 mM), H_2_O_2_ (100 µM), Esterase, Chondroitinase ABC, Hyaluronidase, Lysozyme, β-Glucuronidase, β-Glucosidase (5 µg each) and heparanase (2 µg). For Chondroitinase ABC and Hyaluronidase, 50 µg was also tested. Each sample was performed in triplicates. Then the microplate was sealed with sealing film and incubated at 37 °C for 4 hours. Fluorescence intensity was recorded on Spectramax M5 Multimode plate reader (Molecular Devices, USA). λex/em = 365 nm/455 nm.

### Inhibition studies of heparanase using suramin

To 37.5 µL 40 mM NaOAc (pH 5.0) in 384-well microplate was added 2.5 µL 0.01 µg/µL heparanase. Then 5 µL suramin of various concentrations (0.01, 0.03, 0.1, 0.3, 1.0, 3.0, 10, 30, 100, 300, 1000, 3000, 10000 μM) was added. The plates were sealed and incubated at 37 °C for 1 h. Then probe 5 µL **1** (**HADP**) (50 µM) was added to the microplates. Then the microplates were sealed and incubated at 37 °C from 4 hours. Each compound was performed in triplicates. Fluorescence intensity was recorded on Spectramax M5 Multimode Platereader (Setting: λex = 365 nm, λem = 455 nm). The relative fluorescent intensity was plotted as a function of logarithm of inhibitor concentrations.

### Z’-factor determination

48 wells were used as positive controls (5 μM probe **HADP** with 0.025 µg hepranase, 50 µL total volume) and other 48 wells were used as negative controls (5 μM probe **HADP** only, 50 µL total volume). Then the microplates were sealed and incubated at 37 °C from 4 hours. Fluorescence intensity was recorded on Spectramax M5 Multimode plate reader (Molecular Devices, USA). Then Z’-factor was calculated using

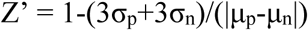

where σ_p_ and σ_n_ are the standard deviations of the positive and negative controls, respectively and μ_p_ and μ_n_ are the means of the positive and negative controls, respectively.

### Screening the commercial library

A library of 1280 compounds (10 mM, dissolved in DMSO) in 96-well format was purchased from Tocris. Transfer 2 µL of the master library into the daughter library containing 198 µL of water to obtain a daughter library (100 µM for each compound). To 35 µL 40 mM NaOAc (pH 5.0) in 384-well microplates were added 5 µL 0.005 µg/µL heparanase. Then the daughter library solution (5 µL) was added. The plates were sealed and incubated at 37 °C for 1 h. Then probe 5 µL **1** (**HADP**) (50 µM) was added to the microplates. Then the microplates were sealed and incubated at 37 °C from 4 hours. Each compound was performed in triplicates. Fluorescence intensity was recorded on Spectramax M5 Multimode Platereader (Setting: λex = 365 nm, λem = 455 nm).

### Measuring the IC_50_ of hit compound

To 40 µL 40 mM NaOAc (pH 5.0) in 384-well microplate was added 5 µL 0.005 µg/µL heparanase. Then 5 µL hit compound of various concentrations (0.1, 0.3, 1.0, 3.0, 10, 30, 100 μM) was added. The plates were sealed and incubated at 37 °C for 1 h. Then probe 5 µL **1** (**HADP**) (50 µM) was added to the microplates. Then the microplates were sealed and incubated at 37 °C from 4 hours. Each compound was performed in triplicates. Fluorescence intensity was recorded on Spectramax M5 Multimode Platereader (Setting: λex = 365 nm, λem = 455 nm). The relative fluorescent intensity was plotted as a function of logarithm of inhibitor concentrations.

### Data availability

The data that support the finding of this study are available from the corresponding authors upon reasonable request.

## Acknowledgements

This work is supported by research grants to Prof. L. Cui from the National Institute of General Medical Sciences of National Institutes of Health (Maximizing Investigators’ Research Award for Early Stage Investigators, R35GM124963), the Department of Defense (Congressionally Directed Medical Research Programs Career Development Award, W81XWH-17-1-0529), University of New Mexico (UNM Startup Award), and the University of Florida (UF Startup Fund). P.K.P.H. was supported by UNM-NIH Initiative for Maximizing Student Development (IMSD) Scholarship. P.Z.F. was supported by Dr. Thomas Whaley Endowed Memorial Scholarship from the Department of Chemistry and Chemical Biology, UNM. Part of the high-resolution mass spectroscopy analysis was performed at the Mass Spectrometry Research and Education Center at the University of Florida supported by grant NIH S10 OD021758-01A1. We thank Profs. Wei Wang (UNM) and Weihong Tan (UF) for the use of their fluorescence spectrometers.

## Author contributions

L.C. conceived the original idea. J.L. and L.C. designed the molecules. J.L. synthesized the compounds and performed the assays. T.L.B., Y.X., K.Y. and J.L. expressed the recombinant human heparanase. Y.W. participated in the preliminary activity test. K.A.S. and A.K. assisted in the optimization for reaction conditions. C.C. assisted in the compound screening and performed fluorescence microscopy. C.X. and H.G conducted the theoretical calculations and analysis. G.S. and C.L. performed docking studies. B.F., P.K.P.H., and P. F. assisted in the chemical synthesis of compounds. J.L. and L.C. analyzed the results and wrote the manuscript. All authors commented on the manuscript.

## Additional information

**Supplementary information** is available.

**Correspondence and requests for materials** should be addressed to L.C.

