## supplemental information for "Ultrasensitive small molecule fluorogenic probe for human heparanase"

### General methods

Deuterated solvents were purchased from Sigma-Aldrich and Merck Millipore. Absorption spectra were recordes on Shimadzu UV-2700 UV-VIS Spectrophotometer. Fluorescence spectra were recorded on Fluorolog TAU-3 Spectrofluorometer with a xenon lamp (Jobin Yvon-Spex, Instruments S. A., Inc.).  Human heparanase was expressed and purified by reported protocol[1].

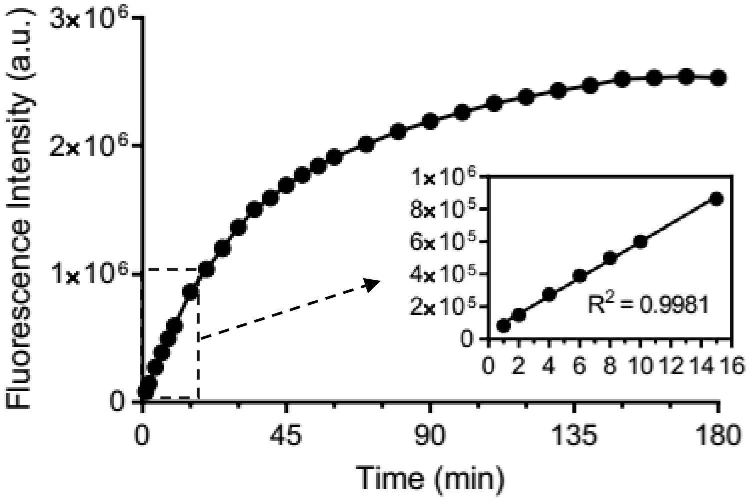

**Supplementary Figure 1.** Quantification of fluorescence intensity (Fig. 3d) at 455 nm of **HADP** (5 µM) with heparanase (1 µg) from 0 min to 3 h in 40 mM NaOAc buffer (pH 5.0). λex = 365 nm. Inset of supplementary Fig. 3b: relationship of fluorescent intensity over time (1-15 min).

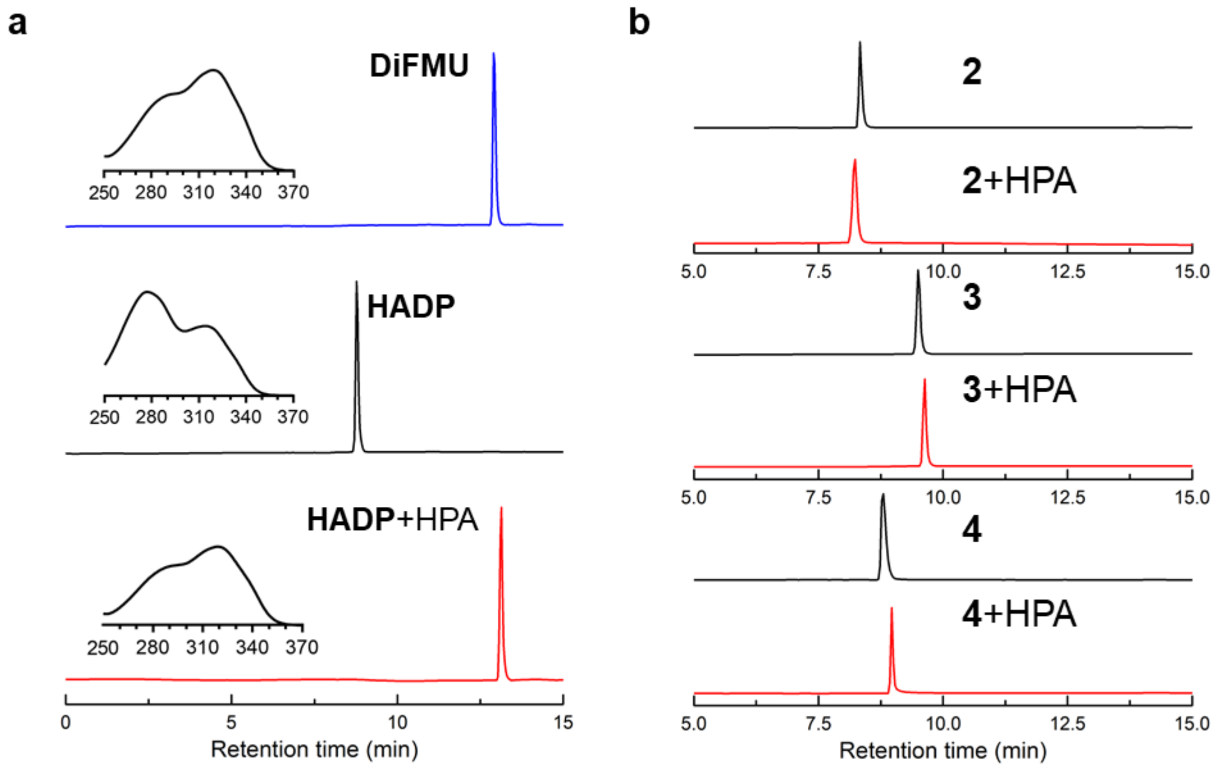


**Supplementary Figure 2.** HPLC traces of **a DiFMU** (blue line), **HADP** (black line) and **HADP** with heparanase (red line), Inset: absorption of each peak; **b** compounds 2-4 without (black line)/with (red line) heparanase.

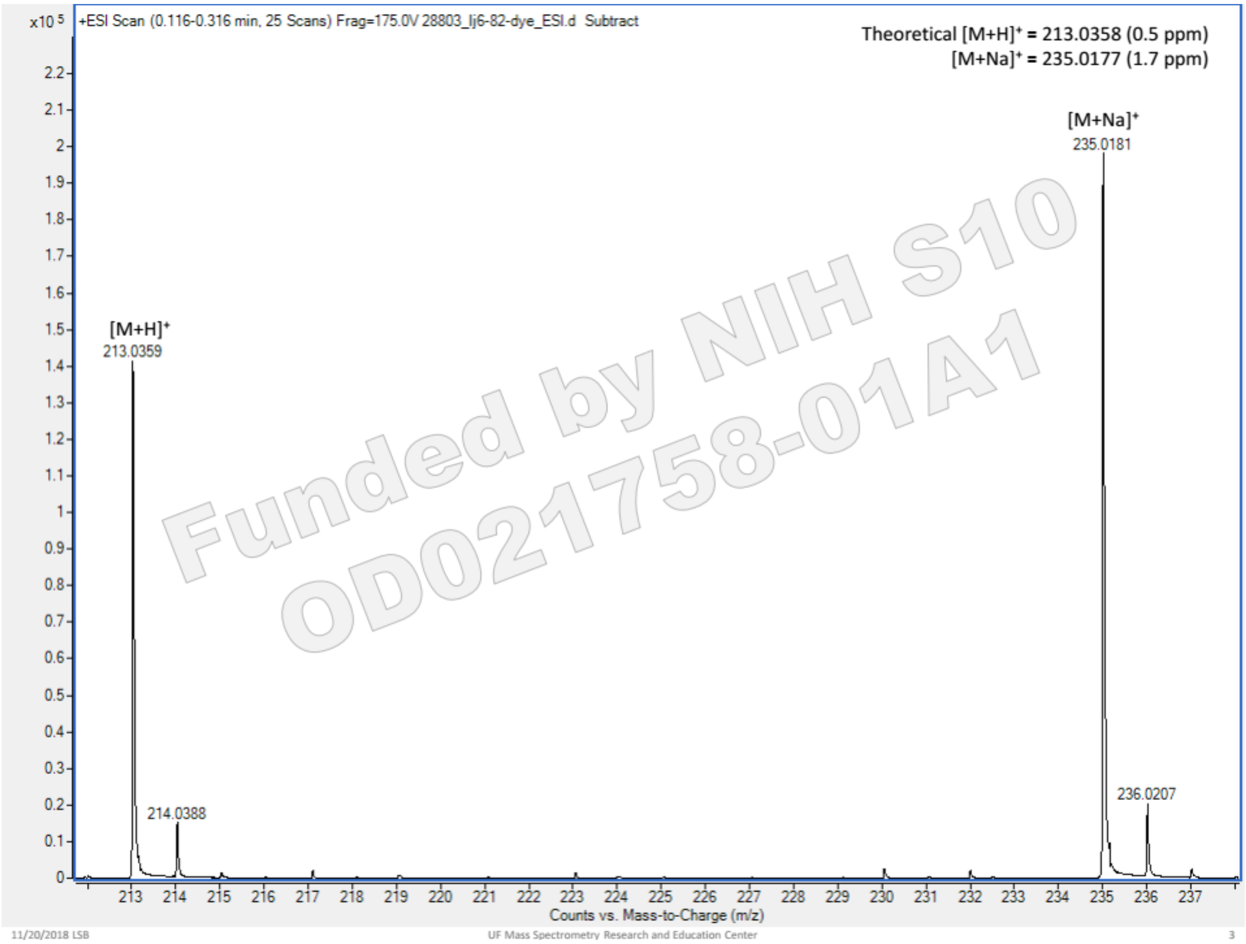


**Supplementary Figure 3**. Mass spectrum of the product after **HADP** was treated with heparanase.

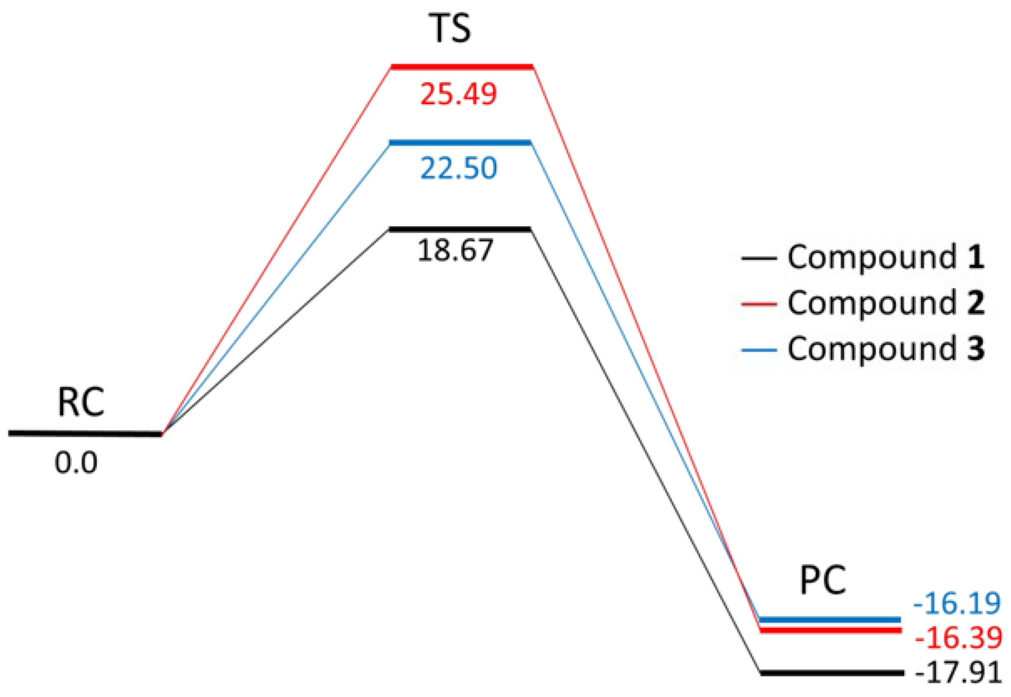


**Supplementary Figure 4** shows the reaction pathways for the three different compounds **1**, **2**, and **3**, which correspond to different substitution groups on the benzene ring. Compound 1 has two F atoms on the ortho-position of benzene, compound **2** has no substitution group on benzene ring and, compound **3** has a substitution group of $CHF_2$ on the ortho-position of benzene. The structures of the stationary points on the reaction pathways for the compounds **1**, **2**, and **3** are shown in Figs. S5-S7. Figure S8 shows the reaction pathways for all compounds. Compound **1** has a lowest barrier (18.67 kcal/mol) due to two strong electron-withdrawing substitution groups (two F atoms) on the ortho-position of benzene. Compound **3** has a higher barrier (22.50 kcal/mol) due to the electron-attracting group $CHF_2$ on the benzene. The highest barrier (25.49 kcal/mol) is found to be for the compound **2** without substitution group on the benzene. This is consistent with the behavior of the bond distances of breaking C-O bond and the charges on C and O atoms in RCs. The bond distance decreases in the order of compounds **1**, **3** and **2**, suggesting the glycosidic bond becomes stronger.

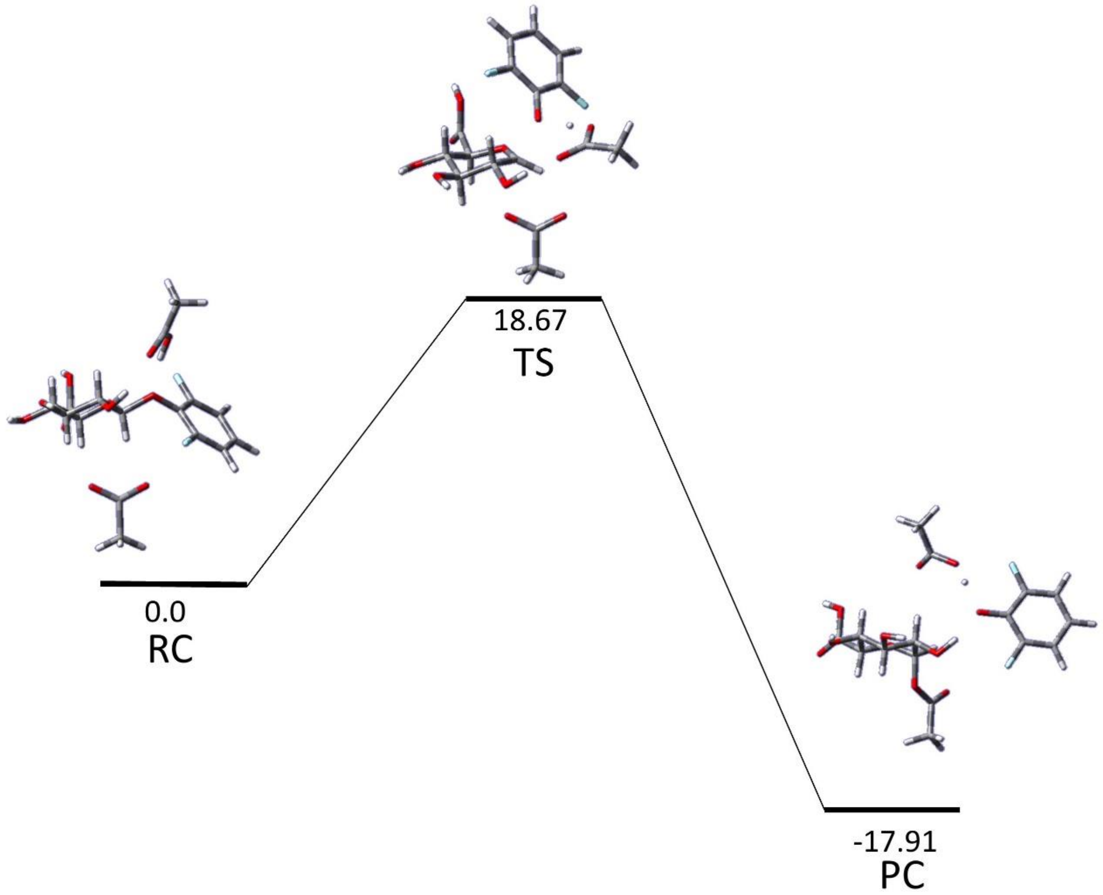


**Supplementary Figure 5**. Relative free energy (in Kcal/mol) profile of the reaction pathway for the compound 1. The structures of RC, TS, and PC are also shown.

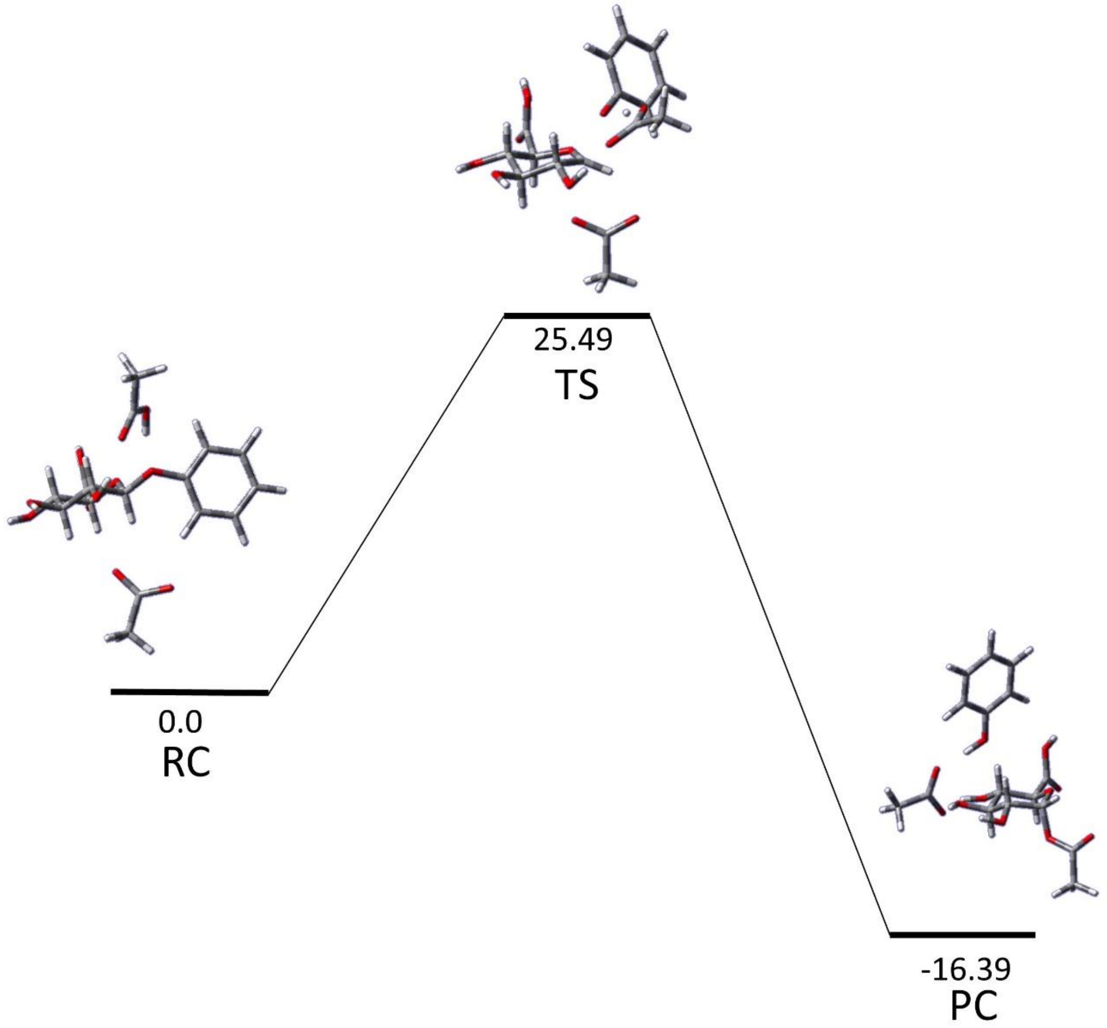


**Supplementary Figure 6.** Relative free energy (in Kcal/mol) profile of the reaction pathway for the compound 2. The structures of RC, TS, and PC are also shown.

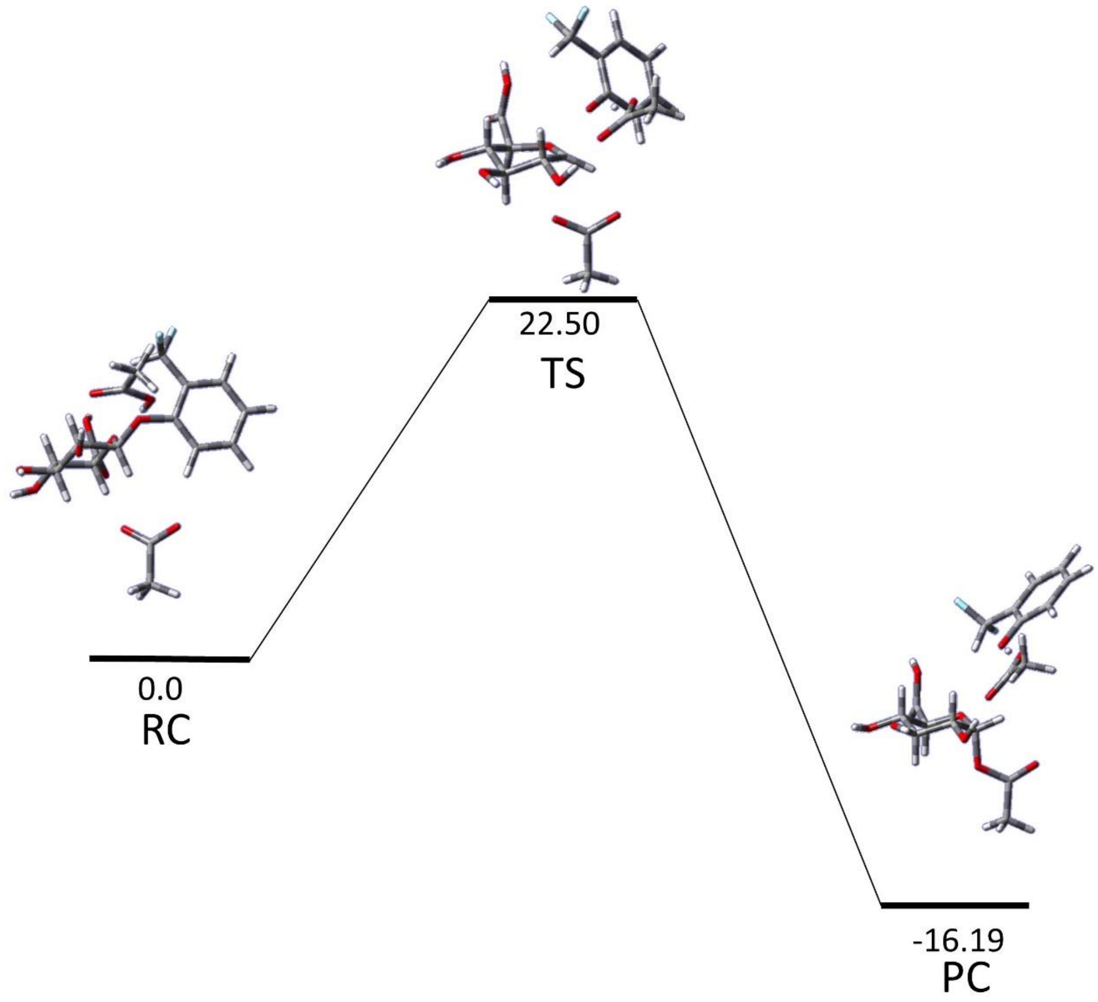


**Supplementary Figure 7**. Relative free energy (in Kcal/mol) profile of the reaction pathway for the compound **3**. The structures of RC, TS, and PC are also shown.

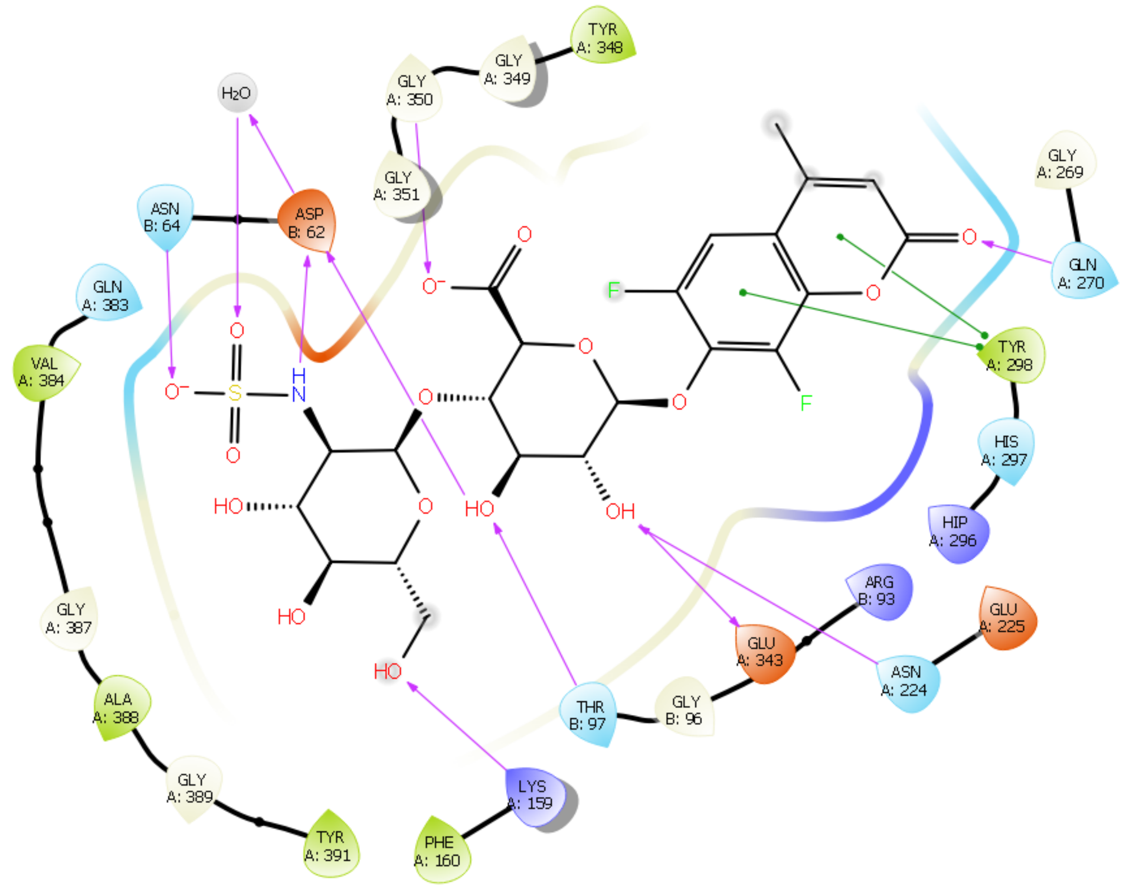


**Supplementary Figure 8**. Interaction diagram of heparanase and compound **1** (**HDAP**).

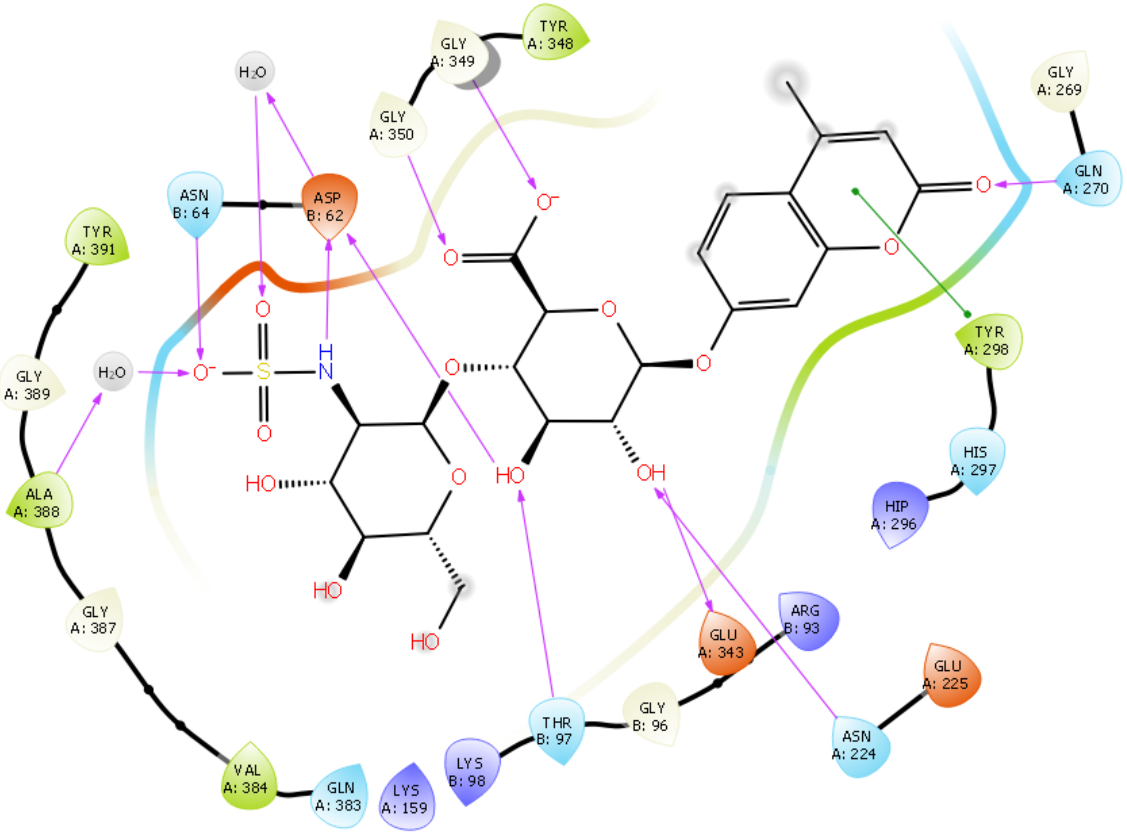


**Supplementary Figure 9**. Interaction diagram of heparanase and compound **2**.

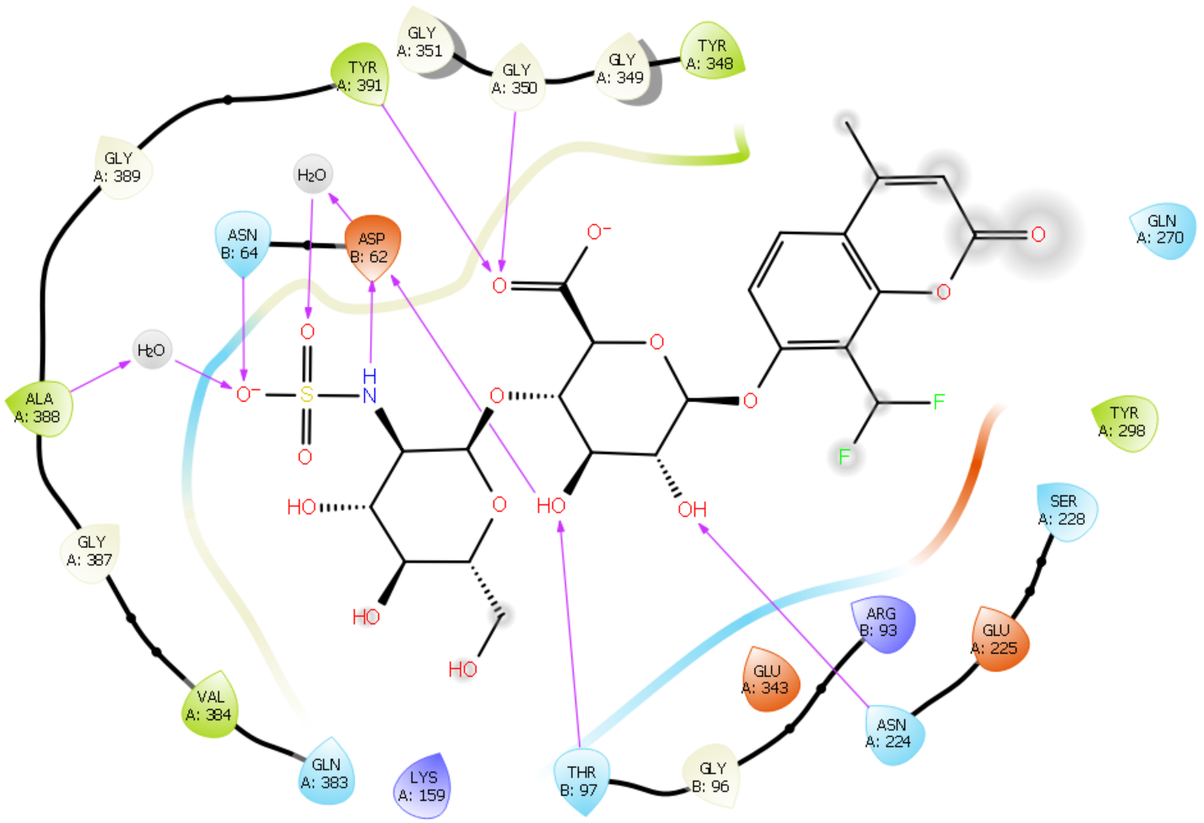


**Supplementary Figure 10**. Interaction diagram of heparanase and compound **3**.

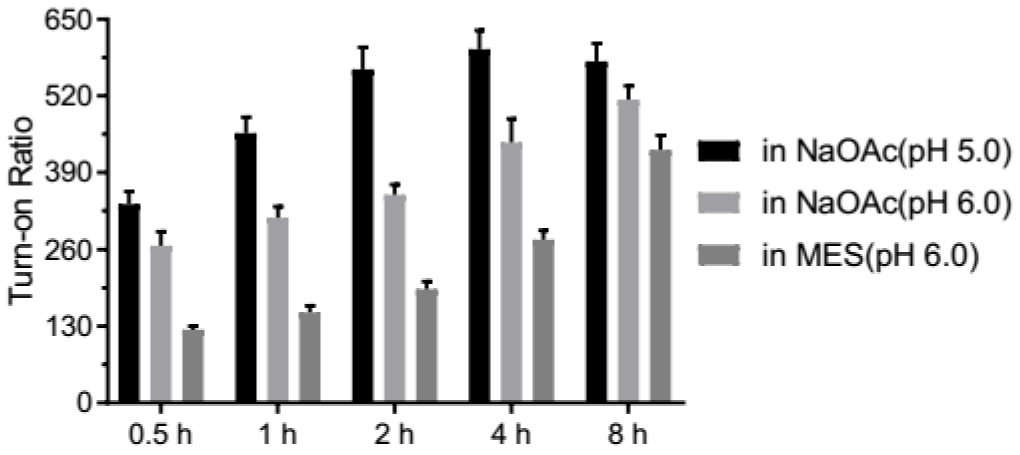


**Supplementary Figure 11.** Effect of buffer on the turn-on ratio of probe **HADP** (5 μM) with heparanase at 37 °C. λex/em = 365 nm/455 nm.

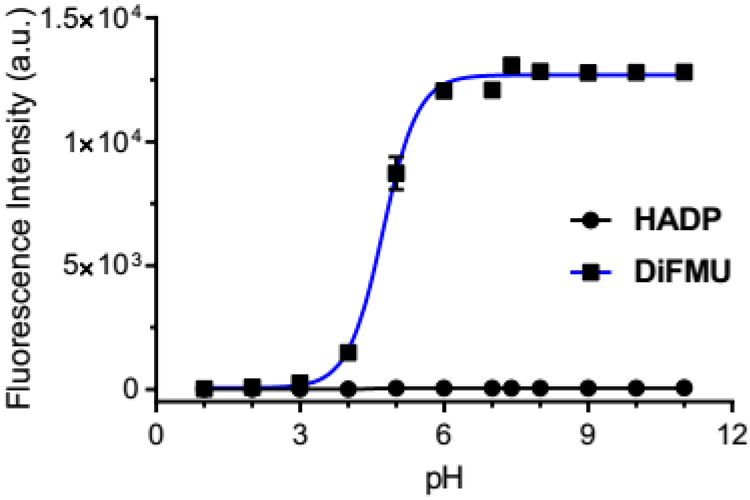


**Supplementary Figure 12.** pH-dependent fluorescence intensity of **HADP** and **DiFMU** (each 5 μM). λex/em = 365 nm/455 nm.

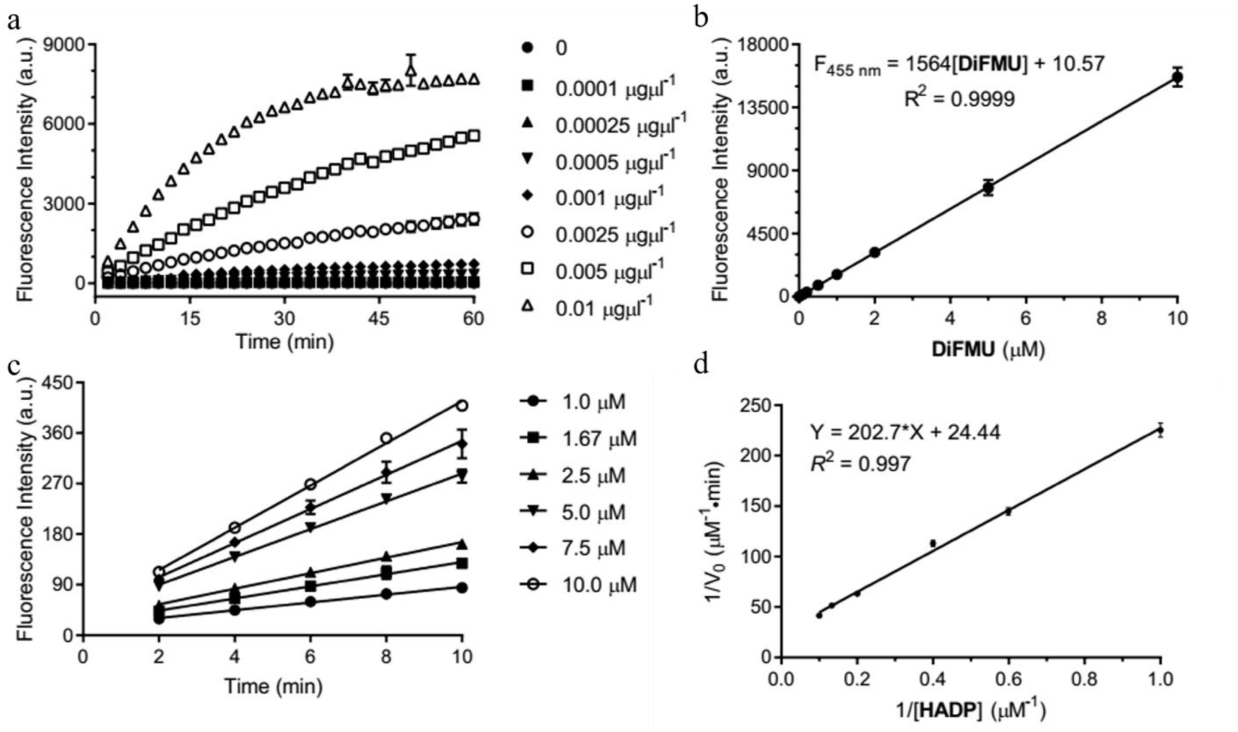


**Supplementary Figure 13.** (a) Time-dependent fluorescence intensity increment using **HADP** (5 μM) with various amounts of heparanase. (b) Standard fluorescence curve of **DiFMU** at different concentrations in 40 mM NaOAc buffer (pH 5.0). λex/em = 365 nm/455 nm. (c) Time-dependent fluorescence intensity increment using heparanase (0.001 $\mu g \mu l^{-1}$) with different concentrations of probe **HADP**. λex/em = 365 nm/455 nm. (d) Lineweaver-Burke plot of the probe activation by heparanase.

### Analysis of the binding mode of the fluorescent probes to Heparanase

There are 7 structures for human heparanase (hHPSE) in the Protein Data Bank (PDB):

| PDBID | Ligand | Resolution | Reference (year) |
|---|---|---|---|
| 5e8m | APO | 1.75 | 2015[1] |
| 5e97 | M04 S00a | 1.63 | |
| 5e98 | M04 S02a | 1.63 | |
| 5e9b | M09 S05a | 1.88 | |
| 5e9c | Dp4 | 1.73 | |
| 5L9Y | JJB355 | 1.88 | 2017[2] |
| 5L9Z (E343Q) | JJB355 | 1.57 | |

All the protein structures are very similar. Most residues are very well conserved, and there are few changes in side chain positions as the protein adapts to the ligand. Lys98, Lys159, Phe160 and Arg303 have small changes adjusting to the glycan tail, while Glu225, Arg272 and Gln270 occupy the space around the ligand head. There are 3 active-site water molecules conserved in all crystal structures. The first bridges the ligand to Ala388, the second to both Phe386 and Asp62, and a third bridging Lys98 to Asp62, locking the orientation of Asp62 in a favorable position.

The dockings were done using Schrödinger Maestro v.12.1.013 (Release 2019-3)[3]. We chose to use the PDB structure from entry 5e98 because the disaccharide in the molecules is identical to the last units in M04-S02a, with modification only in the substituent in the GlcUA unit. Hydrogens were added considering protonation states of the protein and ligands for a pH 5.5 ± 0.5. Alternative positions for protein residues were discarded, keeping only the first available. All ligands except for M04-S02a were removed, as well as the water molecules with the exception of the three conserved waters in the active site. In the end, the protein structure was energy-minimized in the presence of the M04-S02a ligand. The grid box was centered on the center of mass of the M04-S02a ligand, and core constraints were applied so that the RMSD between the ligand's and M04-S02a maximum common substructure be at most at 1.5 Å. The standard induced fit protocol was used, and in the first pass the side chains of Glu225, Gln270 and Arg272 are trimmed to facilitate docking of the ligand and up to 20 poses are generated. After docking, the trimmed residues are rebuilt and all residues within 5Å of the ligand are optimized. In the end, structures within 30 kcal/mol of the best result were redocked with XP precision, keeping the top 20 structures.

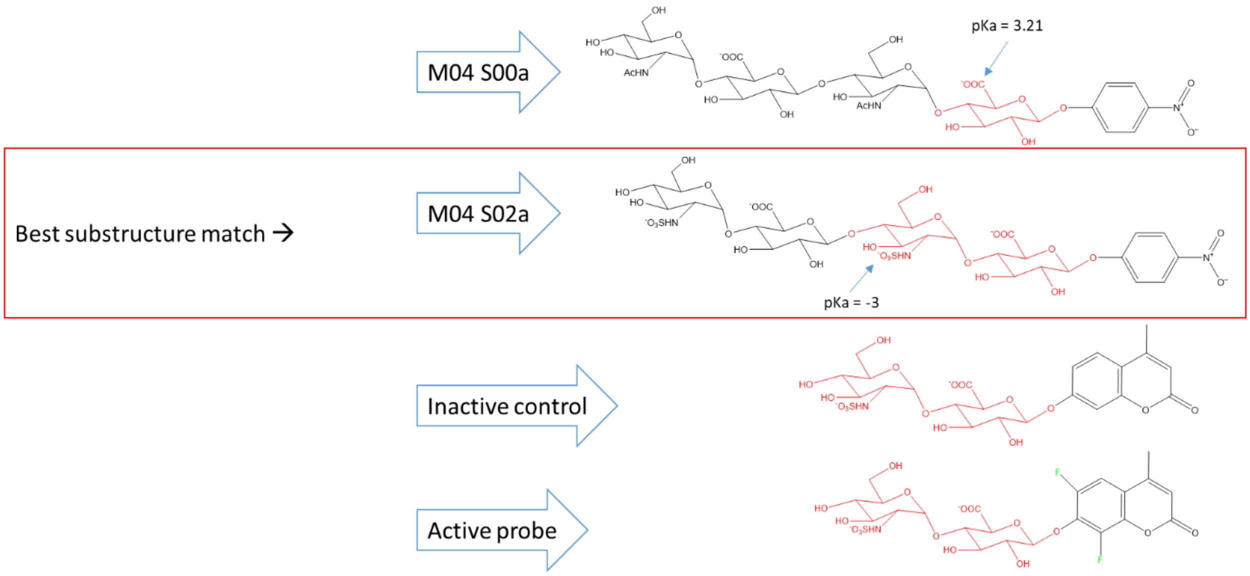


| Ligand | PDB entry | Residue sequence |
|---|---|---|
| M04 S00a | 5e97 | GlcNAc-GlcUA-GlcNAc-**GlcUA-**pNP |
| M04 S02a | 5e98 | GlcNS-GlcUA-**GlcNS-GlcUA-**pNP |
| M09 S05a | 5e9b | GlcUA-GlcNS-GlcUA-GlcNS-GlcUA-GlcNS-GlcUA-GlcNS(6S)-**GlcUA-**pNP |
| Dp4 | 5e9c | HexUA(2S)-GlcNS(6S)-IdoUA-GlcNS(6S) |
| JJB355 (Compound 5) | 5L9Y | 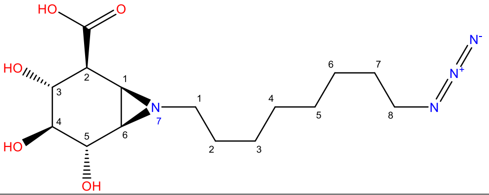 |
| Active probe | | **GlcNS-GlcUA-**6,8-difluoro-4-methyl-coumarin |
| Inactive control | | **GlcNS-GlcUA-**4-methyl-coumarin |

The glycan-tail from the ligands maintained the same interactions with the protein as in M04-S02a, as expected, with only minimal movement of side chains. The coumarin moieties fit into the cavity forming $\pi$-stacking interactions with Tyr348 and Tyr298. The coumarin can flip and, within the top 4 poses of the active probe (**Supplementary Movie 1**) the keto oxygen forms an H-bond to Gln270, and in the best two poses $\pi$-cation interactions are also possible between Lys231 and the coumarin ring. In the inactive probe **(Supplementary Movie 2)** those $\pi$-cation interactions are not noted, and the keto oxygen forms an H-bond to the Lys231 in 2 of the 3 best poses, while maintain the H-bond to Gln270 in the other poses. Although it was allowed to move, Glu225 did not participate in any interactions. In the case of the ligand **(Supplementary Movie 3),** the difluoro-methyl group causes steric clashes with Tyr298, and the coumarin cannot rotate as before, resulting in that all poses are very similar with the keto group pointing away from the active site residues.

Legend for interaction diagrams: *Ligand interaction diagrams for the best pose in each case. Negatively charged residues are depicted in red, positively charged residues in purple, polar residues in light blue and hydrophobic residues in green. Purple arrows indicate H-bonds, and green arrows indicate* π*-stacking interactions. The tip on the residue symbols indicate the orientation of the side chain.*

*Supplementary Table 1*. Results for the best 5 poses in each ligand. Glide XP Score is the energy of the last docking. Prime energy is a binding energy obtained after optimization of the binding site with MM-GBSA and implicit solvent. The final solutions are ranked according to tche Induced Fit Docking score, calculated as IFD Score = 1.0 × Glide XP Score + 0.05 × Prime Energy.

| **Molecule** | **Glide XP Score (kcal/mol)** | **Prime Energy (kcal/mol)** | **IFD Score (kcal/mol)** | **RMSD to Crystal (Å)** |
|---|---|---|---|---|
| M04 S02a | -13.83 | -18,645.00 | -946.08 | 0.35 |
| | -13.67 | -18,643.20 | -945.83 | 0.62 |
| | -13.68 | -18,639.35 | -945.64 | 0.66 |
| | -13.87 | -18,633.50 | -945.54 | 0.72 |
| | -13.17 | -18,630.06 | -944.67 | 0.41 |
| 1 (Active probe) | -15.56 | -18,669.30 | -949.02 | |
| | -15.16 | -18,664.67 | -948.39 | |
| | -14.80 | -18,657.78 | -947.69 | |
| | -13.38 | -18,659.14 | -946.34 | |
| | -12.98 | -18,665.52 | -946.25 | |
| 2 | -15.08 | -18,711.53 | -950.65 | |
| | -13.95 | -18,715.57 | -949.73 | |
| | -13.41 | -18,710.32 | -948.92 | |
| | -13.35 | -18,708.85 | -948.79 | |
| | -12.87 | -18,711.08 | -948.43 | |
| 3 | -14.84 | -18,711.08 | -950.40 | |
| | -14.75 | -18,710.80 | -950.29 | |
| | -14.58 | -18,709.70 | -950.07 | |
| | -14.54 | -18,708.45 | -949.97 | |
| | -14.41 | -18,710.76 | -949.94 | |

**UV-Vis and fluorescence spectrum.** To a solution of 20 µL of 50 µM probe **1** (**HADP**) and 160 µL 40 mM NaOAc (pH = 5.0) was added 20 µL 0.1 µg/µL heparanase to obtain 5 µM **1** (**HADP**) with 2 µg heparanase solution. After incubation at 37 ℃ for 2 h, the reaction solution was transferred to quartz cuvettes to measure absorbance or fluorescence. Absorbance spectrum scan range from 250 nm to 500 nm (1 nm increment). fluorescence spectrum setting: λex = 365 nm, Slit Wicdth 2.5 nm. Emission was record from 400 nm to 600 nm, Slit Width 5 nm. Absorption spectra were recordes on Shimadzu UV-2700 UV-VIS Spectrophotometer. Fluorescence spectra were recorded on Fluorolog TAU-3 Spectrofluorometerwith a xenon lamp (Jobin Yvon-Spex, Instruments S. A., Inc.)

**Time dependence of fluorescence spectra of HADP with heparanase.** Fluorescence micro cells was charged with 20 µL of 50 µM probe **1** (**HADP**) and 160 µL 40 mM NaOAc (pH = 5.0). Then 20 µL 0.05 µg/µL heparanase was added to the above solution. Fluorescence spectra were recorded from 0 min to 3 h. fluorescence spectrum setting: λex = 365 nm, Slit Width 2.5 nm. Emission was record from 400 nm to 600 nm, Slit Width 5 nm.

**Fluorescence responses of probe HADP and control compounds 2-4 to heparanase.** To a solution of 20 µL of 50 µM each compound and 160 µL 40 mM NaOAc (pH = 5.0) was added 20 µL 0.05 µg/µL heparanase to obtain 5 µM each compound with 1 µg heparanase solution in 96-well microplate. Each compound was performed in triplicates. Then the microplate was sealed with sealing film and incubated at 37 ℃. Fluorescence intensity was recorded on Spectramax M5 Multimode plate reader (Molecular Devices, USA) at each timepoint. λex/em = 365 nm/455 nm.

**Fluorescence responses of probe HADP to biological species.** 5 µM probe **1** (**HADP**) was incubated with various biological species at the indicated amount/concentration in 96-well microplate. Cysteine (5 mM), Glutathione (5 mM), $H_2O_2$ (100 µM), Esterase, Chondroitinase ABC, Hyaluronidase, Lysozyme, β-Glucuronidase, β-Glucosidase (5 µg each) and heparanase (2 µg). For Chondroitinase ABC and Hyaluronidase, 50 µg was also tested. Each sample was performed in triplicates. Then the microplate was sealed with sealing film and incubated at 37 ℃ for 4 hours. Fluorescence intensity was recorded on Spectramax M5 Multimode plate reader (Molecular Devices, USA). λex/em = 365 nm/455 nm.

**HPLC analysis of Compounds 1-3 with heparanase.** Heparanase (1 µg, 20 µL) was added to the compounds **1-3** solution (180 µL, 5 µM) in 40 mM NaOAc buffer (pH 5.0) and the mixture was incubated for 4 hours at 37 °C. The resulting mixtures were injected into HPLC for analysis. HPLC analysis was performed under the following conditions - mobile phase A: water with 0.1% TFA; B: acetonitrile with 0.1% TFA; 0-20 min: gradient elution, 25-95% B. The trace was monitored by DAD detector at 315 nm.

**Inhibition studies of heparanase using suramin.** To 37.5 µL 40 mM NaOAc (pH 5.0) in 384-well microplate was added 2.5 µL 0.01 µg/µL heparanase. Then 5 µL suramin of various concentrations (0.01, 0.03, 0.1, 0.3, 1.0, 3.0, 10, 30, 100, 300, 1000, 3000, 10000 µM) was added. The plates were sealed and incubated at 37 ºC for 1 h. Then probe 5 µL **1** (**HADP**) (50 µM) was added to the microplates. Then the microplates were sealed and incubated at 37 °C from 4 hours. Each compound was performed in triplicates. Fluorescence intensity was recorded on Spectramax M5 Multimode Platereader (Setting: λex = 365 nm, λem = 455 nm). The relative fluorescent intensity was plotted as a function of logarithm of inhibitor concentrations.

**Z'-factor determination.** 48 wells were used as positive controls (5 µM probe **HADP** with 0.025 µg hepranase, 50 µL total volume) and other 48 wells were used as negative controls (5 µM probe **HADP** only, 50 µL total volume). Then the microplates were sealed and incubated at 37 °C from 4 hours. Fluorescence intensity was recorded on Spectramax M5 Multimode plate reader (Molecular Devices, USA). Then Z'-factor was calculated using

$$Z' = 1-(3\sigma_p+3\sigma_n)/(|\mu_p-\mu_n|)$$

where $\sigma_p$ and $\sigma_n$ are the standard deviations of the positive and negative controls, respectively and $\mu_p$ and $\mu_n$ are the means of the positive and negative controls, respectively.

**Screening the commercial library.** A library of 1280 compounds (10 mM, dissolved in DMSO) in 96-well format was purchased from Tocris. Transfer 2 µL of the master library into the daughter library containing 198 µL of water to obtain a daughter library (100 µM for each compound). To 35 µL 40 mM NaOAc (pH 5.0) in 384-well microplates were added 5 µL 0.005 µg/µL heparanase. Then the daughter library solution (5 µL) was added. The plates were sealed and incubated at 37 ºC for 1 h. Then probe 5 µL **1** (**HADP**) (50 µM) was added to the microplates. Then the microplates were sealed and incubated at 37 °C from 4 hours. Each compound was performed in triplicates. Fluorescence intensity was recorded on Spectramax M5 Multimode Platereader (Setting: λex = 365 nm, λem = 455 nm).

**Measuring the $IC_{50}$ of hit compound.** To 40 µL 40 mM NaOAc (pH 5.0) in 384-well microplate was added 5 µL 0.005 µg/µL heparanase. Then 5 µL hit compound of various concentrations (0.1, 0.3, 1.0, 3.0, 10, 30, 100 μM) was added. The plates were sealed and incubated at 37 ºC for 1 h. Then probe 5 μL **1** (**HADP**) (50 μM) was added to the microplates. Then the microplates were sealed and incubated at 37 °C from 4 hours. Each compound was performed in triplicates. Fluorescence intensity was recorded on Spectramax M5 Multimode Platereader (Setting: λex = 365 nm, λem = 455 nm). The relative fluorescent intensity was plotted as a function of logarithm of inhibitor concentrations.

**Effect of buffer.** To a solution of 20 μL of 50 μM probe **HADP** (**1**) and 160 μL different buffers was added 20 μL 0.1 μg/μL heparanase to obtain 5 μM each compound with 2 μg heparanase solution in 96-well microplate. Each buffer was performed in triplicates. Then the microplate was sealed with sealing film and incubated at 37 °C. Fluorescence intensity was recorded on Spectramax M5 Multimode plate reader (Molecular Devices, USA) at each timepoint. λex/em = 365 nm/455 nm. Buffers: 40 mM NaOAc (pH 5.0); 40 mM NaOAc (pH 6.0); 50 mM MES with 10 mM NaCl (pH = 6.0).

**Time-dependent fluorescence intensity with various amounts of heparanase.** To a solution of 20 μL of 50 μM probe **HADP** (**1**) and 160 μL 40 mM NaOAc buffer (pH 5.0) buffers was added 20 μL heparanase with various concentrations (0-0.1 μg/μL) in 96-well microplate. Then final concentration of heparanase is from 0-0.01 μg/μL. Each concentration was performed in triplicates. Then the fluorescence intensity was recorded on Spectramax M5 Multimode plate reader (Molecular Devices, USA) at each timepoint. λex/em = 365 nm/455 nm.

**Time-dependent fluorescence intensity increment using heparanase (0.001 $\mu g \mu l^{-1}$) with different concentrations of probe HADP.** To a solution of 4, 6.7, 10, 20, 30, 40 μL of 50 μM probe **HADP** (**1**) and 176, 173.3, 170, 160, 150, 140 μL 40 mM NaOAc buffer (pH 5.0) buffers was added 20 μL heparanase (0.01 μg/μL) in 96-well microplate. Then final concentration of **HDAP** in each well is 1.0, 1.67, 2.5, 5.0, 7.5, 10 μM and heparanase is 0.001 μg/μL. Each concentration was performed in triplicates. Then the fluorescence intensity was recorded on Spectramax M5 Multimode plate reader (Molecular Devices, USA) at each timepoint. λex/em = 365 nm/455 nm.

**Detection Limit**. To calculate the detection limit, plot the fluorescence intensity against heparanase concentration and fit a line with linear regression. The detection limit was estimated with the following formula. LOD= 3 σ/s, where σ represents standard deviation of fluorescent intensity of blank sample, s is the calculated slope of linear regression equation.

**Transition energy calculations.** The geometry optimizations and vibrational frequency calculations of the stationary points, including the reactant complex (RC), the product complex (PC), and the transition state (TS), were performed at the Becke's three parameter hybrid method with the LYP correlation function (B3LYP)[4-6] level in gas phase with the 6-31+G(d) basis set for all the atoms. The vibrational analysis and intrinsic reaction coordinate (IRC) calculations were performed to confirm the transition states at the same computational level. The solvation energies were calculated using the polarized continuum model (PCM)[7-10] at temperature 37℃, the dielectric constant used in the calculation is ε=5.0 for the protein. All calculations were carried out using the GAUSSIAN 09.[11]

**Synthesis of the probes**

The donor[12] and compound S6[13] were prepared following the literature procedures.

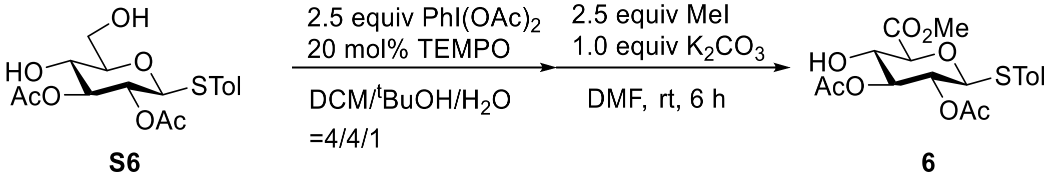


**S6** **6**

Methyl p-Tolyl (2,3-di-O-acetyl-1-thio-β-D-glucopyranosyluronate: To the solution of *p*-tolyl-2,3-di-*O*-acetyl-1-thio-β-D-glucopyranoside (S6, 3.7 g, 10 mmol) and TEMPO (312.5 mg, 2 mmol) was added iodobenzene diacetate (8.0 g, 25 mmol) in DCM/$^t$BuOH/$H_2O$ (4:4:1, 45 mL). The reaction was stirred at room temperature for 5 h to afford uronic acid. Then reaction was quenched with saturated $Na_2S_2O_3$ and extracted with ethyl acetate followed by concentration. The residue was dissolved in DMF (40 mL). then $K_2CO_3$ (1.38 g, 10 mmol) and methyl iodide (1.6 mL, 25 mmol) were added to above solution. The reaction mixture was stirred at room temperature for 5 h. After the completion of reaction, the mixture was added with cold water, extracted with ethyl acetate and washed with water for multiple times and then with brine, followed by drying over anhydrous $Na_2SO_4$. The organic fraction was then concentrated and purified by column chromatography (Hexanes: Ethyl Acetate = 5:1 to 2:1) to give the desired compound as a white solid (3.37 g, 85%).

$^{1}$H NMR (800 MHz, $CDCl_3$) δ 7.37 (d, $J$ = 8.0 Hz, 2H), 7.11 (d, $J$ = 8.0 Hz, 2H), 5.12 (dd, $J_{3,2}$ = $J_{3,4}$= 9.2 Hz, 1H, H-3), 4.88 (dd, $J$ = 9.6 Hz, 1H, H-2), 4.68 (d, $J_{1,2}$ = 9.6 Hz, 1H, H-1), 3.92 (d, $J_{5,4}$ = 9.6 Hz, 1H, H-5), 3.89 (ddd, $J_{4,5}$ = $J_{4,3}$ = 10.4 Hz, $J_{4,4\text{-OH}}$ = 4.0 Hz, 1H, H-4), 3.82 (s, 3H, -$OCH_3$), 3.29 (d, $J_{4\text{-}OH,4}$ = 4.0 Hz, 1H, 4-OH), 2.32 (s, 3H, -$CH_3$), 2.07 (s, 3H, $OCOCH_3$), 2.04 (s, 3H, $OCOCH_3$). $^{13}$C NMR (201 MHz, $CDCl_3$) δ 170.7 (O*C*OCH$_3$), 169.3 (O*C*OCH$_3$), 168.9 ($COOCH_3$), 138.8, 133.7, 129.9, 128.2, 87.0 (C-1), 78.2 (C-5), 75.9 (C-3), 70.2 (C-2,4), 52.8 (COO*C*H$_3$), 21.2 (-*C*H$_3$), 20.7 (2×OCO*C*H$_3$). HRMS (ESI) for $C_{18}H_{22}O_8SNa$ $[M+Na]^+$: Calcd: 421.0933, Found: 421.0939.

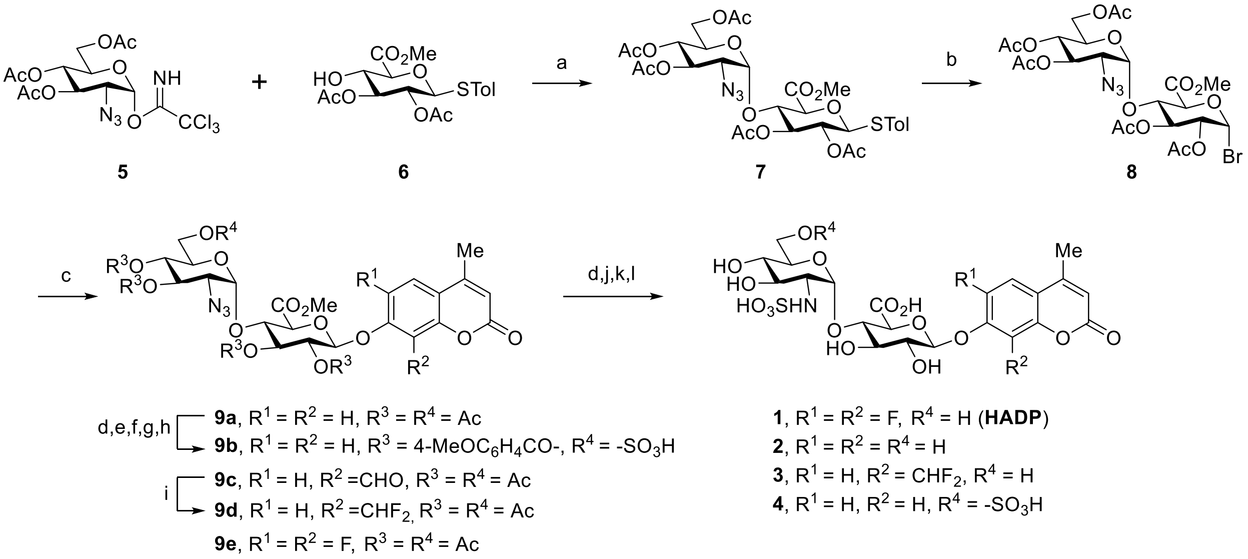


Reagents and Conditions: a) TMSOTf, 4 Å MS,Toluene/1,4-Dioxane=2/1, 0 °C; b) IBr, DCM, rt; c) 4-methylcoumarin derivatives, $Ag_2O$, MeCN; d) NaOMe, MeOH; e) TrCl, Py; f) 4-$MeOC_6H_4COCl$, DMAP, Py; g) TFA, DCM; h) $Et_3N{\cdot}SO_3$, DMF; i) DAST, DCM; j) $H_2$ balloon, Pd/C, MeOH, rt, 4 h; k) $H_2O$/THF=2/1, NaOH (pH=11), rt; l) Py· $SO_3$,rt.

With the acceptor and donor in hand, we embark on screening appropriate conditions for stereoselective construction of 1,2-*cis*-α-linkage taking advantage of the nonparticipation effect of azido group as a masked amine of donor. The glycosylation of **6** with trichloroacetimidate **5** was carried out employing trimethylsilyl triflate (TMSOTf) as promoter in the presence of 4 Å molecular sieves in dichloromethane (DCM) from -70 °C to room temperature. Only a little amount of desired disaccharide was observed. The unexpected intermolecular aglycone transfer[14-16] of **10** was the predominant product mainly due to the less reactive hydroxyl group on the 4$^{th}$ position. Switching the solvent to toluene increased the formation of desired disaccharide. Further screening shows that 48% of disaccharide was isolated after increasing reaction temperature to 0 °C. To our delight, significant improvement was achieved in the 2/1 mixture of toluene and 1,4-dioxane, yielding the disaccharide **7** in 79% yield with a complete 1,2-*cis*-α-selectivity, although it is impossible to thoroughly eliminate the formation of aglycone transfer product. The α-configuration of newly formed glyosidic bond was unambiguously determined by expected doublet for anomeric proton of the glucosamine moiety.

After successfully obtaining the disaccharide, we attempted to incorporate the fluorescent tag on the disaccharide moiety. Transformation of **7** into the corresponding brominated product and further coupling with the fluorescent tag afforded the expected product in satisfying yield (55%) using silver oxide ($Ag_2O$) as activating agent in acetonitrile. Global deacetylation of **9a** with sodium methoxide (NaOMe) in methanol, reduction of azido group to amine by hydrogenation, subsequent saponification reaction and final sulfation led to the desired compound **2**. Substrate **3** were obtained with the similar method.

It was postulated that **4** would likely be a better substrate owing to an additional sulfate group on 6-*O* position of the glucosamine moiety, providing extra affinity to heparanase. Recently, a docking study suggested that **4** should be a better substrate. Herein, we reported the successful synthesis of this compound. Starting with the intermediate **9a**, hydroxyl group on 6-postion on the glucosamine residue was selectively protected by bulky trityl group for the further sulfation. Upon completion of tritylation, 4-methoxybenzoyl group was employed to block the rest of hydroxyl groups. Then the trityl group was selectively removed under acid condition and sulfation was conducted on the exposed 6$^{th}$ position of glucosamine motif to introduce the desired group. It is noteworthy that 4-methoxybenzoyl group instead of simple acetyl group as a protecting group is to avoid acyl migration[17] from the 4$^{th}$ position to 6$^{th}$ position in the sulfation step. The following steps are identical with the synthesis of compounds **2** and **3** described above.

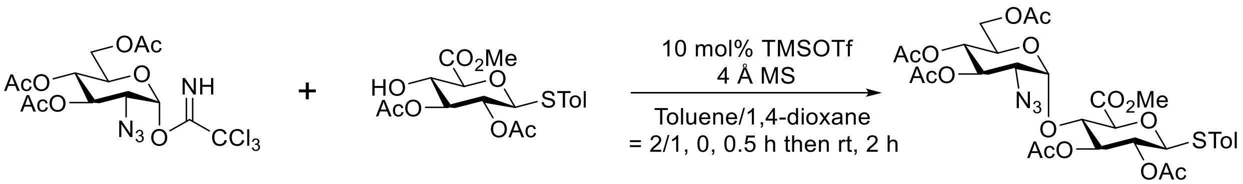

5 6 7

**Methyl *p*-Tolyl (3’,4’,6’-tri-*O*-acetyl-2’-azido-2’-deoxy-α-D-glucopyranosyl)-(1→4)-(2,3-di-O-acetyl-1-thio-β-D-glucopyranosyluronate:** 6-Tri-*O*-acetyl-2-azido-2-deoxy-α-D-glucopyranosyl trichloroacetimidate (951 mg, 2 mmol) and methyl p-tolyl (2,3-di-O-acetyl-1-thio-β-D-glucopyranosyluronate (398.4 mg, 1 mmol) were added into a dry round bottom flask and kept under high vacuum for 0.5 h. Then the flask was flushed with nitrogen and sealed with a rubber septum. Anhydrous toluene (4 mL) and 1,4-dioxane (2 mL) were added and stirred for 30 min. The reaction mixture was then maintained at 0 °C for addition of TMSOTf (18 μL, 0.1 mmol). After 0.5 h the reaction was allowed to stir at room temperature for next 2 h. The reaction was quenched with triethyl amine and filtered, concentrated. Then residue was purified by column chromatography (Hexane: Ethyl Acetate = 3:1 to 2:1) to give the desired compound as a white solid (560.8 mg, 79%). $^{1}$H NMR (800 MHz, $CDCl_3$) δ 7.33 (d, $J$ = 8.0 Hz, 2H), 7.11 (d, $J$ = 7.2 Hz, 2H), 5.30-5.27 (m, 2H, H-3, H-3’), 5.14 (d, $J_{1',2'}$ = 3.2 Hz, 1H, H-1’), 4.98 (dd, $J_{4',3'}$ = $J_{4',5'}$ = 10.4 Hz, 1H, H-4’), 4.80 (dd, $J_{2,1}$ = $J_{2,3}$ = 9.6 Hz, 1H, H-2), 4.69 (d, $J_{1,2}$ = 10.4 Hz, 1H, H-1), 4.23 (dd, $J_{6a',6b'}$ = 12.8 Hz, $J_{6a',5'}$ = 3.2 Hz, 1H, H-6a’), 4.12 (dd, $J_{4,3}$ = $J_{4,5}$ = 9.6 Hz, 1H, H-4), 4.06 (d, $J$ = 12 Hz, 1H, H-6b’), 3.99 (d, $J_{5,4}$ = 9.6 Hz, 1H, H-5), 3.79-3.77 (m, 4H, $COOCH_3$, H-5’), 3.34 (dd, $J_{2,3}$ = 10.4 Hz, $J_{2,1}$ = 3.2 Hz, 1H, H-2’), 2.33 (s, 3H, $-CH_3$), 2.06 (s, 3H, $COCH_3$), 2.05 (s, 3H, $COCH_3$), 2.03 (s, 3H, $COCH_3$), 2.01 (s, 3H, $COCH_3$), 1.99 (s, 3H, $COCH_3$). $^{13}$C NMR (201 MHz, $CDCl_3$) δ 170.5 (O*C*O$CH_3$), 169.7 (O*C*O$CH_3$), 169.5 (2×O*C*O$CH_3$), 169.4 (O*C*O$CH_3$), 167.7 (*C*OOCH3), 138.9, 133.9, 129.9, 127.7, 98.7 (C-1’), 86.7 (C-1), 78.0 (C-5), 76.0 (C-4), 75.3 (C-3), 70.6 (C-3’), 70.5 (C-2), 68.8 (C-5’), 68.6 (C-4’), 61.3 (C-6’), 61.3 (C-2’), 52.8 (-COO*C*$H_3$), 21.2 (-*C*$H_3$), 20.7 (2×CO*C*$H_3$), 20.6 (2×CO*C*$H_3$), 20.5 (-CO*C*$H_3$). HRMS (ESI) for $C_{30}H_{37}N_3O_{15}SNa$ $[M+Na]^+$: Calcd: 734.1843, Found: 734.1859.

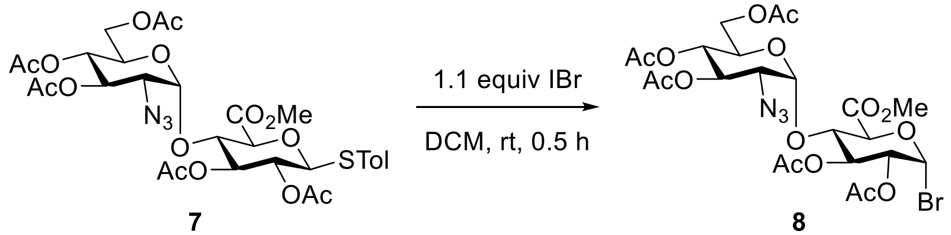

7 8

**Methyl (3’,4’,6’-Tri-*O*-acetyl-2’-azido-2’-deoxy-α-D-glucopyranosyl)-(1→4)-(1-bromo-1-deoxyl-2,3-di-*O*-acetyl-β-D-glucopyranosyluronate:** To a solution of **7** (569 mg, 0.8 mmol) in anhydrous dichloromethane (5 mL), freshly prepared 1M IBr solution (0.9 mL, 0.90 mmol, prepared in anhydrous dichloromethane) was added and stirred at room temperature for 0.5 h under dark condition. After the completion of the reaction it was quenched with saturated $Na_2S_2O_3$ and extracted with DCM followed by drying over anhydrous $Na_2SO_4$. Thus, the organic fraction was concentrated and subjected to column chromatography (Hexane: Ethyl Acetate = 2:1) for purification to give the desired compound as a white solid (394 mg, 74 %). $^1$H NMR (800 MHz, $CDCl_3$) δ $^1$H NMR (800 MHz, $CDCl_3$) δ 6.52 (d, $J_{1,2}$ = 4.0 Hz, 1H, H-1), 5.65 (dd, $J_{3,2}$ = $J_{3,4}$ = 9.6 Hz, 1H, H-3), 5.34 (dd, $J_{3',4'}$ = $J_{3'2'}$ = 9.6 Hz, 1H, H-3’), 5.16 (d, $J_{1',2'}$, = 4.0 Hz, 1H, H-1’), 5.01 (dd, $J_{4',5'}$ = $J_{4',3'}$ = 9.6 Hz, 1H, H-4’), 4.78 (dd, $J_{2,3}$ = 9.6 Hz, $J_{2,1}$ = 4.0 Hz, 1H, H-2), 4.53 (d, $J_{5,4}$ = 9.6 Hz, 1H, H-5), 4.24-4.21 (m, 2H, H-4, H-6a’), 4.08 (dd, $J_{6b',6a'}$ = 12.8 Hz, $J_{6b',5'}$ = 1.6 Hz, 1H, H-6b’), 3.87 (d, *J* = 9.6 Hz, 1H, H-5’), 3.81 (s, 3H, -$COOCH_3$), 3.45 (dd, $J_{2',3'}$ = 10.4, $J_{2'1'}$ = 4.0 Hz, 1H, H-2’), 2.08 (s, 9H, $COCH_3$), 2.06 (s, 3H, $COCH_3$), 2.02 (s, 3H, $COCH_3$). $^{13}$C NMR (201 MHz, $CDCl_3$) δ 170.6 (O*C*O$CH_3$), 169.9 (O*C*O$CH_3$), 169.7 (O*C*O$CH_3$), 169.6 (O*C*O$CH_3$), 169.0 (O*C*O$CH_3$), 167.6 (*C*OO$CH_3$), 98.8 (C-1’), 85.5 (C-1), 75.9 (C-4), 74.1 (C-5), 71.1 (C-2), 70.8 (C-3), 70.6 (C-3’), 69.1 (C-5’), 68.5 (C-4’), 61.5 (C-6’), 61.4 (C-2’), 53.2 (-COO*C*H3), 20.7 (2×COCH3), 20.6 (3×COCH3). HRMS (ESI) for $C_{23}H_{30}BrN_3O_{15}Na$ $[M+Na]^+$: Calcd: 690.0758, Found: 690.0771.

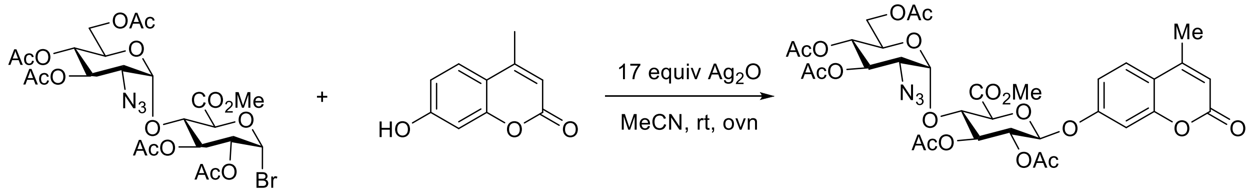

8 4-Methylumbelliferone 9a

**4’-methylumbelliferyl-3,4,6-Tri-*O*-acetyl-2-azido-2-deoxy-α-D-glucopyranosyl-(1→4)-methyl 2,3-di-*O*-acetyl-β-D-glucopyranosyluronate.** General Procedure for the glycosylation of disaccharide bromide with fluorescent tag. To the solution of the disaccharide bromide (**8**) and 4-methylcoumarin derivatives in anhydrous MeCN was added silver oxide (2 equiv or 17 equiv) at room temperature under dark condition. After the completion of the reaction it was filtered and concentrated. The residue was purified with column chromatography (Hexane: Ethyl Acetate = 1:1 to 1:2) to give the desired compound. **9a** 55% yild. $^{1}$H NMR (500 MHz, $CDCl_3$) δ 7.55 (d, *J* = 8.5 Hz, 1H), 6.96 – 6.92 (m, 2H), 6.22 (s, 1H), 5.41 (t, *J* = 8.5 Hz, 1H, H-3), 5.38 (t, *J* = 10.0 Hz, 1H, H-3’), 5.34 (d, *J* = 6.5 Hz, 1H, H-1), 5.27 (d, *J* = 3.5 Hz, 1H, H-1’), 5.20 (dd, *J* = 8.5, 6.5 Hz, 1H, H-2), 5.04 (t, *J* = 10.0 Hz, 1H, H-4’), 4.43 (t, *J* = 8.5 Hz, 1H, H-4), 4.30 (d, *J* = 9.0 Hz, 1H, H-5), 4.27 (dd, J = 9.0 Hz, 3.5 Hz, 1H, H-6’), 4.11 (dd, *J* = 12.5, 1.5 Hz, 1H, H-6’), 3.90 (ddd, *J* = 10.5, 2.0, 2.0 Hz, 1H, H-5’), 3.72 (s, 3H), 3.43 (dd, *J* = 11.0, 4.0 Hz, 1H, H-2’), 2.42 (s, 3H), 2.12 (s, 3H), 2.10 (s, 3H), 2.09 (s, 6H), 2.05 (s, 3H). $^{13}$C NMR (126 MHz, $CDCl_3$) δ 170.8, 169.9, 169.74, 169.73, 169.6, 167.5, 160.9, 158.9, 154.8, 152.3, 125.9, 115.7, 113.6, 113.3, 104.3, 98.9, 98.2, 75.1, 74.0, 72.9, 71.3, 70.1, 68.6, 68.0, 61.2, 60.9, 53.1, 20.8, 20.8, 20.7, 18.8. HRMS (ESI) for $C_{33}H_{38}N_3O_{18}^+$ $[M+H]^+$: 764.2145, Found 764.2090.

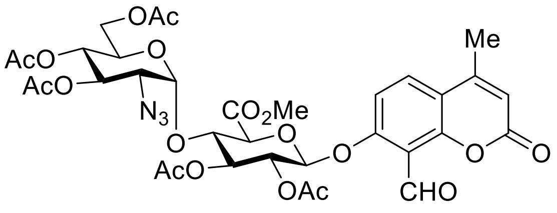


**9c**

**8’-formyl-4’-methyl-umbelliferyl-3,4,6-Tri-*O*-acetyl-2-azido-2-deoxy-α-D-glucopyranosyl-(1→4)-methyl 2,3-di-*O*-acetyl-β-D-glucopyranosyluronate.** Compound **9c** was synthesized in 76% yield according to the glycosylation procedure employed for the synthesis of compound **9a**. $^{1}$H NMR (800 MHz, $CDCl_3$) δ 10.50 (s, 1H, -CHO), 7.69 (d, *J* = 8.8 Hz, 1H), 7.06 (d, *J* = 8.8 Hz, 1H), 6.20 (s, 1H), 5.36 (d, $J_{1,2}$ = 5.6 Hz, 1H, H-1), 5.32 (dd, $J_{3',2'}$ = $J_{3',4'}$ = 10.0 Hz, 1H, H-3’), 5.28 (dd, $J_{3,2}$ = $J_{3,4}$ = 7.6 Hz, 1H, H-3), 5.19 (d, $J_{1',2'}$ = 4.0 Hz, 1H, H-1’), 5.17 (dd, $J_{2,1}$ = 6.0 Hz, $J_{2,3}$ = 7.6 Hz, 1H, H-2), 4.97 (dd, $J_{4',5'}$ = $J_{4',3'}$= 10.0 Hz, 1H, H-4’), 4.43 (dd, $J_{4,5}$ = $J_{4,3}$ = 8.0 Hz, 1H, H-4), 4.31 (d, $J_{5,4}$ = 8.0 Hz, 1H, H-5), 4.17 (dd, $J_{6a',6b'}$ = 12.0 Hz, $J_{6a',5}$ = 4.0 Hz, 1H, H-6a’), 4.05 (dd, $J_{6b',6a'}$ = 12.8 Hz, $J_{6a',5}$ = 1.6 Hz, 1H, H-6b’), 3.87 (d, $J_{5',4'}$ = 9.6 Hz, 1H, H-5’), 3.62 (s, 3H, $CO_2CH_3$), 3.36 (dd, $J_{2',3'}$ = 10.8 Hz, $J_{2',1'}$ = 3.6 Hz, 1H, H-2’), 2.37 (s, 3H, $CH_3$), 2.07 (s,

3H, $OCOCH_3$), 2.06 (s, 3H, $OCOCH_3$), 2.02 (s, 3H, $OCOCH_3$), 2.01 (s, 3H, $OCOCH_3$), 1.97 (s, 3H, $OCOCH_3$). $^{13}$C NMR (201 MHz, $CDCl_3$) δ 186.1 (*C*HO), 170.5 (O*C*OCH$_3$), 169.9 (O*C*OCH$_3$), 169.7 (O*C*OCH$_3$), 169.6 (2xO*C*OCH$_3$), 167.8 (*C*O$_2$Me), 159.2 (=CH*C*O$_2$Ar), 159.0, 155.3, 151.4, 130.1, 116.1, 115.0, 114.2, 112.1, 98.8 (C-1'), 98.6 (C-1), 74.8 (C-4), 74.4 (C-5), 72.1 (C-3), 70.9 (C-2), 70.6 (C-3'), 69.1 (C-5'), 68.8 (C-4'), 61.8 (C-6'), 61.5 (C-2'), 52.9 (CO$_2$*C*H$_3$), 20.8 (OCO*C*H$_3$), 20.7 (3xOCO*C*H$_3$), 20.6 (OCO*C*H$_3$), 18.8. HRMS (ESI) for $C_{34}H_{37}N_3O_{19}Na$ $[M+Na]^+$: Calcd: 814.1919, Found: 814.1913.

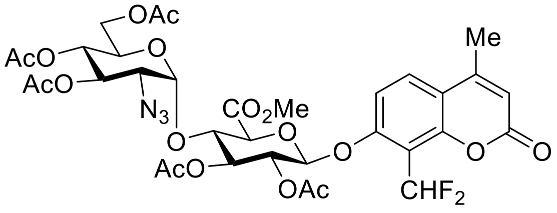


**9d**

**8'-difluoromethyl-4'-methylumbelliferyl-3,4,6-Tri-*O*-acetyl-2-azido-2-deoxy-α-D-glucopyranosyl-(1→4)-methyl 2,3-di-*O*-acetyl-β-D-glucopyranosyluronate (9d)**. To a solution of **9c** (67 mg, 0.085 mmol) in dry DCM (2 mL) was added DAST (37 µL, 0.255 mmol, 3 equiv) at room temperature under argon, the reaction mixture was stirred at room temperature for 18 h. Then the mixture was quenched with ice water. After separation, the aqueous phase was extracted with DCM, and the combined organic phases were dried over $Na_2SO_4$ and concentrated. The resulting residue was purified by column chromatography ((Hexanes: Ethyl Acetate = 1/1-1/2)) to give the desired compound as a white solid (62.1 mg, 90 %).

$^1$H NMR (800 MHz, $CDCl_3$) δ 7.62 (d, $J$ = 8.8 Hz, 1H), 7.14 (t, $J_{H,F}$ = 51.2 Hz, 1H), 7.06 (d, $J$ = 8.8 Hz, 1H), 6.17 (s, 1H), 5.33-5.32 (m, 2H, H-1, H-3'), 5.27 (dd, $J_{3,2}$ = $J_{3,4}$ = 7.6 Hz, 1H, H-3), 5.18 (d, 1H, $J_{1',2'}$ = 3.2 Hz, H-1'), 5.16 (dd, $J_{2,3}$ = $J_{2,1}$ = 7.2 Hz, H-2), 4.96 (dd, $J_{4',3'}$ = $J_{4',5'}$ = 10.0 Hz, 1H, H-4'), 4.41 (dd, $J_{4,3}$ = $J_{4,5}$ = 7.6 Hz, 1H, H-4), 4.33 (d, $J_{5,4}$ = 7.2 Hz, 1H, H-5), 4.17 (dd, $J_{6a',6b'}$ = 12.8 Hz, $J_{6a',5}$ = 2.4 Hz, 1H, H-6a'), 4.06 (d, $J_{6b',6a'}$ = 12.0 Hz, 1H, H-6b'), 3.89 (dd, $J_{5',4'}$ = 8.8 Hz, $J_{5',6a'}$ = 1.6 Hz, 1H, H-5'), 3.62 (s, 3H, $CO_2CH_3$), 3.36 (dd, $J_{2',3'}$ = 10.4, $J_{2',1'}$ = 1.6 Hz, 1H, H-2'), 2.36 (s, 3H, $CH_3$), 2.04 (s, 6H, 2x$OCOCH_3$), 2.02 (s, 6H, 2x$OCOCH_3$), 1.97 (s, 3H, $OCOCH_3$). $^{13}$C NMR (201 MHz, $CDCl_3$) δ 170.5 (O*C*OCH$_3$), 169.8 (O*C*OCH$_3$), 169.6 (O*C*OCH$_3$), 169.5 (O*C*OCH$_3$), 169.4 (O*C*OCH$_3$), 167.7 (*C*O$_2$CH$_3$), 159.0, 157.9, 153.0, 151.6, 128.2, 116.0, 114.1, 111.9, 110.04 (t, $J_{CF}$ = 239.4 Hz, *C*HF$_2$), 98.8 (C-1'), 98.6 (C-1), 74.9 (C-4),

74.3 (C-5), 72.0 (C-3), 70.6 (C-3', C-2), 69.1 (C-5'), 68.9 (C-4'), 61.8 (C-6'), 61.5 (C-2'), 52.9 ($CO_2CH_3$), 20.7 ($OCOCH_3$), 20.6 (2x$OCOCH_3$), 20.5 (2x$OCOCH_3$), 18.7. HRMS (ESI) for $C_{34}H_{37}F_2N_3O_{18}Na$ $[M+Na]^+$: Calcd: 836.1938, Found: 836.1961.

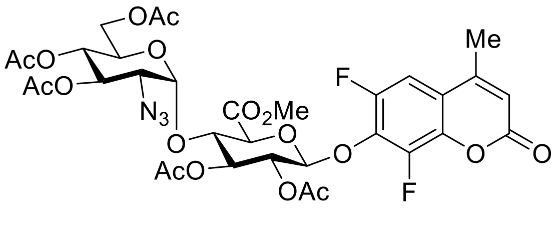


**9e**

**6',8'-difluoro-4'-methylumbelliferyl-3,4,6-Tri-*O*-acetyl-2-azido-2-deoxy-α-D-glucopyranosyl-(1→4)-methyl 2,3-di-*O*-acetyl-β-D-glucopyranosyluronate (9e)**. Compound **9e** was synthesized in 80% yield according to the glycosylation procedure employed for the synthesis of compound **9a**. $^1$H NMR (500 MHz, $CDCl_3$) δ 7.15 (d, $J_{H\text{-}F}$ = 10.5 Hz, 1H), 6.34 (s, 1H), 5.36 (m, 2H, H-3, H-3'), 5.31 (d, *J* = 7.5 Hz, 1H, H-1), 5.25 (t, *J* = 7.5 Hz, 1H, H-2), 5.22 (d, *J* = 3.0 Hz, 1H, H-1'), 5.03 (t, *J* = 9.5 Hz, 1H, H-4'), 4.39 (t, *J* = 9.0 Hz, 1H, H-4), 4.26 (dd, *J* = 12.5, 2.5 Hz, 1H, H-6a'), 4.13 (d, *J* = 9.5 Hz, 1H, H-5), 4.09 (d, *J* = 12.5 Hz, 1H, H-6b'), 3.86 – 3.81 (m, 1H, H-5'), 3.79 (s, 3H), 3.42 (dd, *J* = 10.5, 3.0 Hz, 1H, H-2'), 2.40 (s, 3H), 2.12 (s, 3H), 2.11 (s, 3H), 2.09 (s, 3H), 2.07 (s, 3H), 2.02 (s, 3H). $^{13}$C NMR (126 MHz, $CDCl_3$) δ 170.7, 169.9, 169.7, 169.6, 167.3, 158.8, 151.3 (dd, $J_{C\text{-}F}$ = 248.2, 2.5 Hz), 151.1, 143.4 (dd, $J_{C\text{-}F}$ = 257.0, 4.8 Hz), 139.7 (dd, $J_{C\text{-}F}$ = 10.2, 2.5 Hz), 135.2 (dd, $J_{C\text{-}F}$ =11.5, 16.0 Hz), 116.9 (d, $J_{C\text{-}F}$ = 8.7 Hz), 116.0, 105.9 (dd, $J_{C\text{-}F}$ =21.9, 3.7 Hz), 101.5, 98.9, 75.7, 74.4, 73.3, 71.4, 70.2, 68.7, 68.1, 61.2, 60.9, 53.2, 20.8, 20.7, 20.6, 18.9. HRMS (ESI) for $C_{33}H_{35}F_2N_3O_{18}Na^+$ $[M+Na]^+$: Calcd: 822.1776, Found: 822.1744.

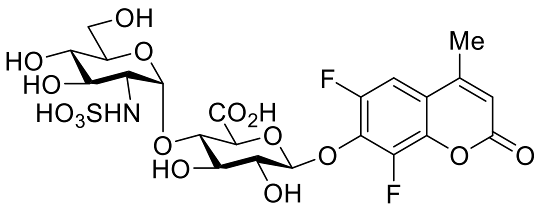

**1** (**HADP**)

**6’,8’-difluoro-4’-methylumbelliferyl-2-deoxy-2-sulfonamido-α-D-glucopyranosyl-(1→4)-β-D-glucopyranosyluronate (1, HADP).** General procedure for global deacetylation, saponification, reduction and sulfation. To a solution of disaccharide in MeOH was added NaOMe (25% w/w NaOMe in MeOH, 0.2 equiv). Then reaction process was monitored with HPLC. When all the acetyl group was removed, the reaction was quenched and neutralized by adding Dowex® resin 50WX8 followed by filtration and concentration. Then the crude compound was dissolved in methanol and 20 wt% Pd/C was added to the mixture. Then the flask was evacuated and back-filled with hydrogen balloon. After completion of hydrogenation monitored with HPLC, the reaction mixture was filtered and concentrated. The crude amine was dissolved in THF/$H_2O$ (or MeOH/$H_2O$) = 1/2, the pH of the solution was adjusted to 11 with 1M NaOH and maintained until the methyl group was removed. Finally, Py•$SO_3$ (10 equiv) was added in portions to above solution and the pH of the solution was maintained between 9-10. After completion of sulfation, the mixture was purified with HPLC to afford desired product after lyophilization. (overall yield 42%) $^1$H NMR (600 MHz, $D_2O$) δ 7.37 (d, $J$ = 11.4 Hz, 1H), 6.36 (s, 1H), 5.69 (d, $J$ = 3.0 Hz, 1H, H-1’), 5.24 (d, $J$ = 7.8 Hz, 1H, H-1), 3.98 (dd, $J_{3,2}$ = $J_{3,4}$ = 9.0 Hz, 1H, H-3), 3.92 (dd, $J_{4,3}$ = $J_{4,5}$ = 9.0 Hz, 1H, H-4), 3.88 (d, $J$ = 9.6 Hz, 1H, H-5), 3.84 – 3.79 (m, 2H, H-6a’, H-6b’), 3.76 – 3.72 (m, 2H, H-2, H-5’), 3.62 (dd, $J_{3',4'}$ = $J_{3',2'}$ = 9.6 Hz, 1H, H-3’), 3.50 (dd, $J_{4',3'}$ = $J_{4',5'}$ = 9.6 Hz, 1H, H-4’), 3.27 (dd, $J_{2',3'}$ = 10.2 Hz, $J_{2',1'}$ = 3.6 Hz, 1H, H-2’), 2.38 (s, 3H). $^{13}$C NMR (151 MHz, $D_2O$) δ 174.3, 162.1, 154.8, 151.3 (dd, $J_{C\text{-}F}$ = 3.0, 246.1 Hz), 142.8 (dd, $J_{C\text{-}F}$ = 6.0, 252.2 Hz), 138.6 (dd, $J_{C\text{-}F}$ = 1.5, 10.6 Hz), 135.0 (dd, $J_{C\text{-}F}$ = 12.1, 16.6 Hz), 116.7 (d, $J_{C\text{-}F}$ = 10.6 Hz), 114.0, 106.7 (dd, $J_{C\text{-}F}$ = 3.0, 22.6 Hz), 103.3, 97.3, 76.9, 76.0, 76.0, 73.1, 71.6, 71.2, 69.6, 60.1, 58.0, 17.9. HRMS (ESI) for $C_{22}H_{29}F_2N_2O_{16}S^+$ $[M+NH_4]^+$ : Calcd: 647.1200, Found: 647.1212.

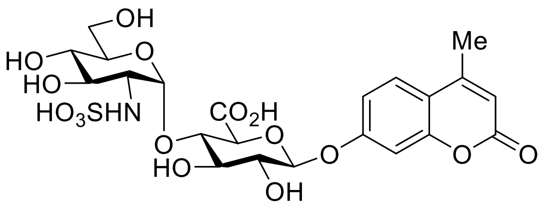

**2**

**4’-methylumbelliferyl-2-deoxy-2-sulfonamido-α-D-glucopyranosyl-(1→4)-β-D-glucopyranosyluronate (2).** Compound **2** was synthesized in 92% yield according to the general procedure employed for the synthesis of compound **1**. $^{1}$H NMR (600 MHz, $D_2O$) δ 7.67 (d, $J$ = 9.0 Hz, 1H), 7.09 (dd, $J$ = 9.0, 2.4 Hz, 1H), 7.05 (s, 1H), 6.21 (s, 1H), 5.70 (d, $J$ = 3.6 Hz, 1H, H-1’), 5.27 (d, $J$ = 7.8 Hz, 1H, H-1), 4.05 (d, $J$ = 9.6 Hz, 1H, H-5), 4.01 (dd, $J$ = $J$ = 9.0 Hz, 1H), 3.93 (dd, J = $J$ = 9.0 Hz, 1H), 3.85 – 3.79 (m, 2H), 3.78 – 3.74 (m, 1H), 3.72 (dd, $J$ = $J$ = 9.0 Hz, 1H), 3.65 (dd, $J$ = $J$ = 9.0 Hz, 1H), 3.52 (dd, $J$ = $J$ = 9.6 Hz, 1H), 3.28 (dd, $J$ = 10.2, 3.6 Hz, 1H), 2.41 (s, 3H). $^{13}$C NMR (151 MHz, $D_2O$) δ 174.5, 164.6, 159.4, 156.3, 153.8, 126.7, 115.3, 113.9, 111.2, 103.6, 99.4, 97.3, 76.8, 76.1, 75.9, 72.5, 71.6, 71.2, 69.6, 60.1, 58.0, 17.9. HRMS (ESI) for $C_{22}H_{31}N_2O_{16}S^+$ $[M+NH_4]^+$ : Calcd: 611.1389, Found: 611.1386.

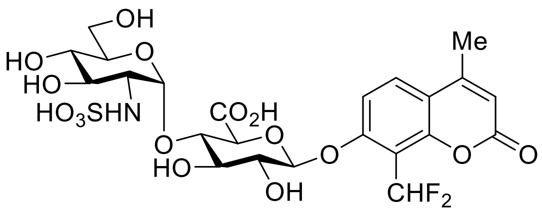

**3**

**8’-difluoromethyl-4’-methylumbelliferyl-2-deoxy-2-sulfonamido-α-D-glucopyranosyl-(1→4)-β-D-glucopyranosyluronate (3).** Compound **3** was synthesized in 60% yield according to the general procedure employed for the synthesis of compound **1.** $^{1}$H NMR (600 MHz, $D_2O$) δ 7.92 (d, $J$ = 9.0 Hz, 1H), 7.41 (t, $J$ = 53.4 Hz, 1H), 7.28 (d, $J$ = 9.0 Hz, 1H), 6.35 (s, 1H), 5.70 (d, $J_{1',2'}$ = 3.6 Hz, 1H, H-1’), 5.33 (d, $J_{1,2}$ = 7.8 Hz, 1H, H-1), 4.03 (d, $J_{5,4}$ = 9.6 Hz, 1H, H-5), 3.99 (dd, $J_{4,5}$ = $J_{4,3}$ = 9.0 Hz, 1H, H-3), 3.93 (dd, $J_{4,3}$ = $J_{4,5}$ = 9.2 Hz, 1H, H-4), 3.84-3.81 (m, 2H, H-6’), 3.77 (dd, $J_{2,3}$=$J_{2,1}$ = 8.4 Hz, 1H, H-2), 3.75-3.73 (m, 1H, H-5’), 3.64 (dd, $J_{3',4'}$ = $J_{3',5'}$ = 9.6 Hz, 1H, H-3’), 3.51 (dd, $J_{4',3'}$ = $J_{4',5'}$ = 9.6 Hz, 1H, H-4’), 3.27 (dd, $J_{2',3'}$ = 10.4 Hz, $J_{2',1'}$= 3.6 Hz, 1H, H-2’),

2.47 (s, 3H). $^{13}$C NMR (151 MHz, $D_2O$) δ 174.4, 157.6, 156.0, 151.8, 129.7, 115.8, 113.4, 112.1, 111.9, 110.8 (t, $J_{CF}$ = 235.6 Hz, *C*HF$_2$),109.9, 99.9, 97.3, 76.8, 76.0, 75.9, 72.4, 71.6, 71.2, 69.6, 60.1, 58.0, 18.1. HRMS (ESI) for $C_{23}H_{31}F_2N_2O_{16}S^+$ $[M+NH_4]^+$ : Calcd: 661.1357, Found: 661.1378.

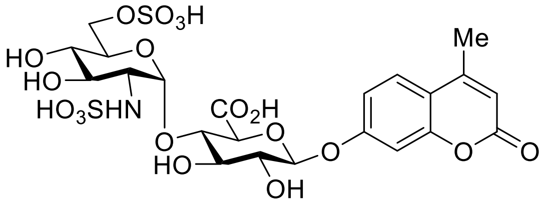

**4**

**4'-methylumbelliferyl-6-O-sulfonato-2-deoxy-2-sulfonamido-α-D-glucopyranosyl-(1→4)-β-D-glucopyranosyluronate (4).** Compound **4** was synthesized in 32% yield according to the general procedure employed for the synthesis of compound **1.** $^1$H NMR (600 MHz, $D_2O$) δ 7.79 (d, $J$ = 8.4 Hz, 1H), 7.17-7.15 (m, 2H), 6.31 (s, 1H), 5.72 (d, $J$ = 3.6 Hz, 1H, H-1'), 5.29 (d, $J$ = 7.8 Hz, 1H, H-1), 4.37 (d, $J$ = 10.8 Hz, 1H, H-6a'), 4.19 (d, $J$ = 10.8 Hz, 1H, H-6b'), 4.05 (d, $J$ = 9.6 Hz, 1H, H-5), 4.01 (dd, $J_{3,2}$ = $J_{3,4}$ = 9.0 Hz, 1H, H-3), 3.94-3.89 (m, 2H, H-4, H-5'), 3.72 (d, $J$ = 7.8 Hz, 1H, H-2), 3.65(dd, $J_{3',4'}$ = $J_{3',2'}$ = 9.0 Hz, 1H, H-3'), 3.61 (dd, $J_{4',3'}$ = $J_{4',5'}$ = 9.0 Hz, 1H, H-4'), 3.31 (dd, $J_{2',3'}$ = 10.2 Hz, $J_{2',1'}$ = 3.6 Hz, 1H, H-2'), 2.49 (s, 3H). $^{13}$C NMR (151 MHz, $D_2O$) δ 174.47, 164.77, 159.47, 156.37, 154.03, 126.76, 115.54, 113.78, 111.36, 103.81, 99.51, 97.27, 76.63, 76.00, 75.91, 72.41, 71.07, 69.78, 68.95, 66.28, 57.87, 17.97. HRMS (ESI) for $C_{22}H_{26}NO_{19}S_2^-$ $[M-H]^-$: Calcd: 672.0546, Found: 672.0527.

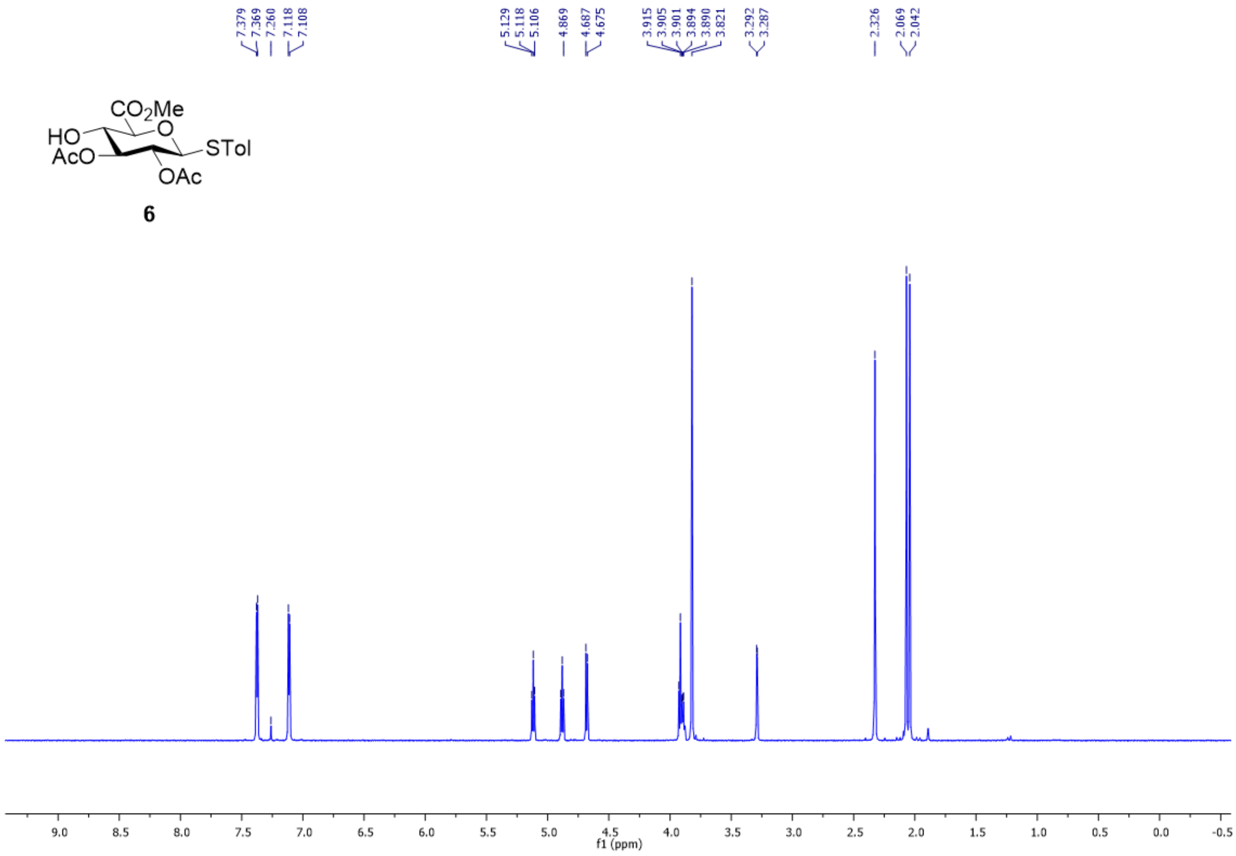


Fig. S1. $^{1}$H NMR spectrum of **6** in $CDCl_3$.

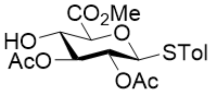


**6**

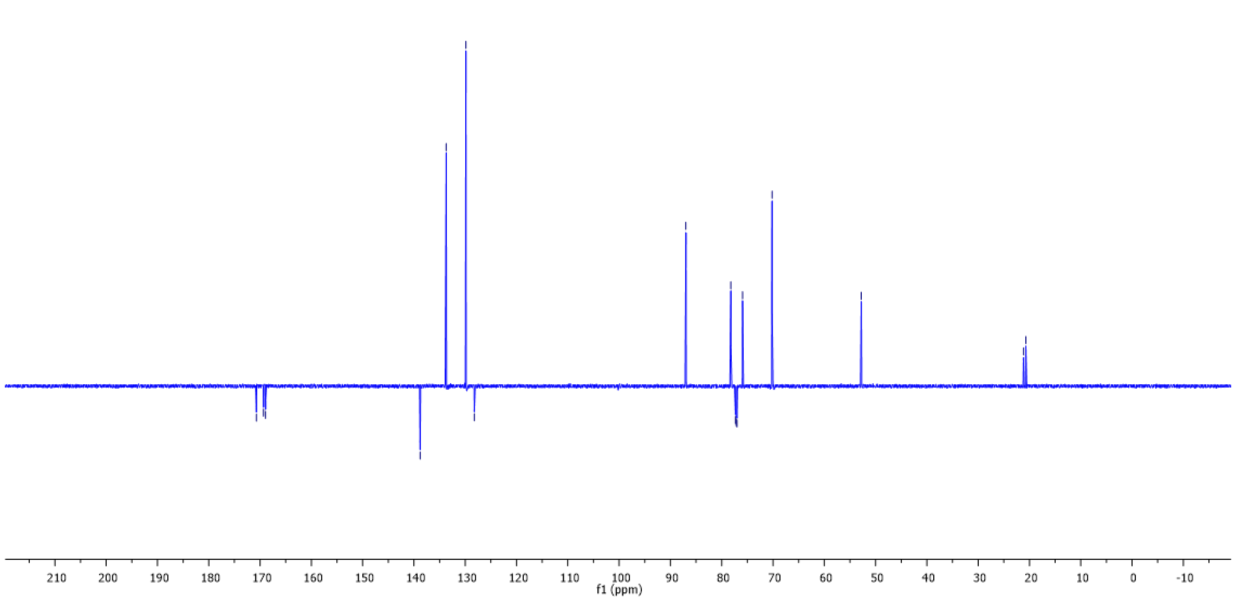


Fig. S2. $^{13}C$ NMR spectrum of **6** in $CDCl_3$.

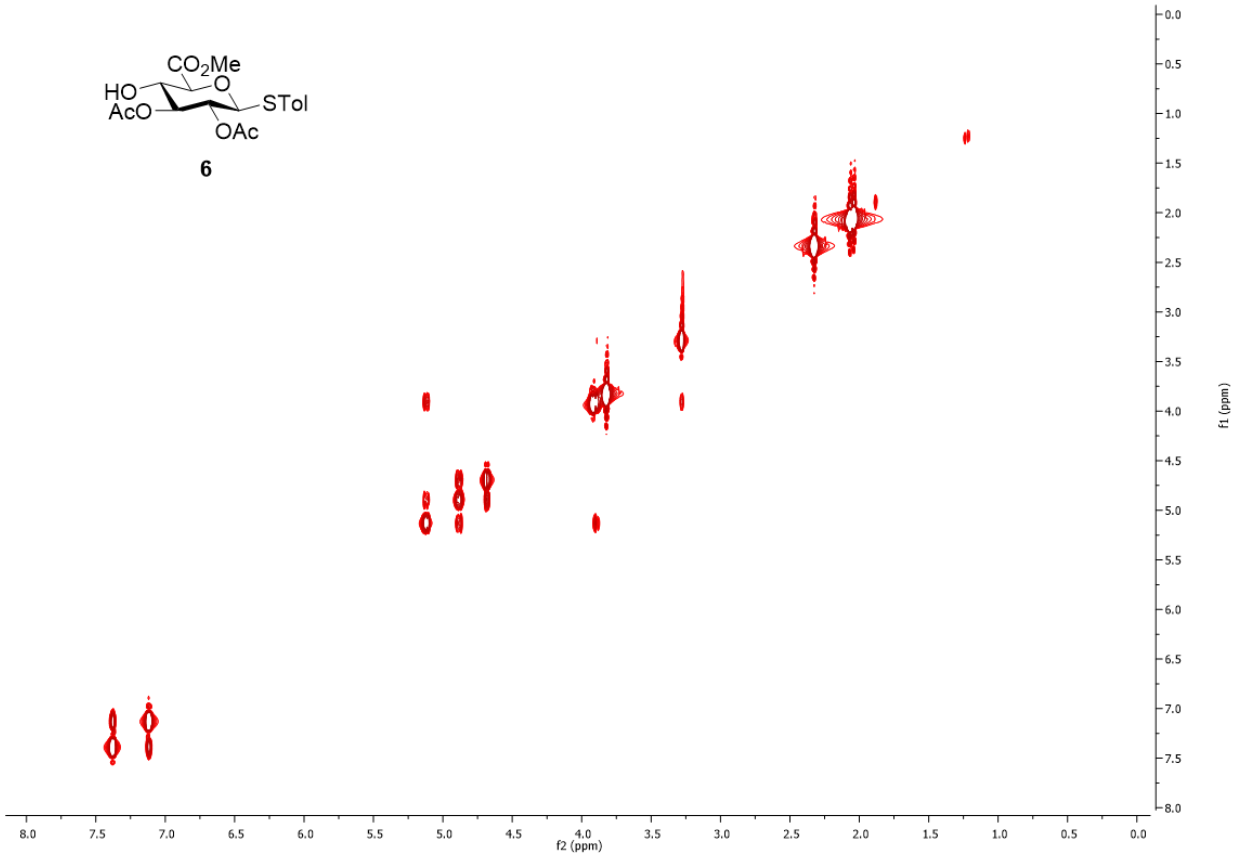


Fig. S3. $^{1}$H-$^{1}$H COSY spectrum of **6** in CDCl3.

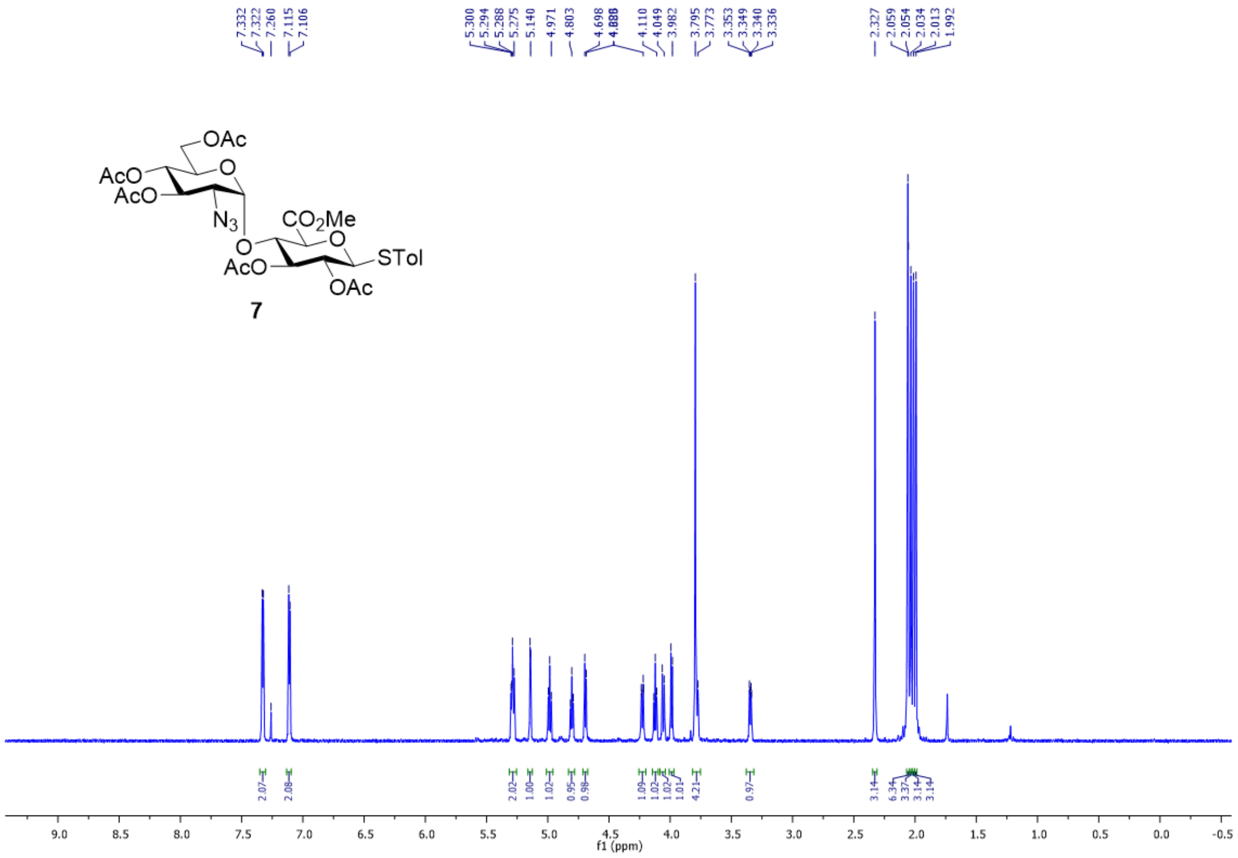

Fig. S4. [1]H NMR spectrum of **7** in $CDCl_3$.

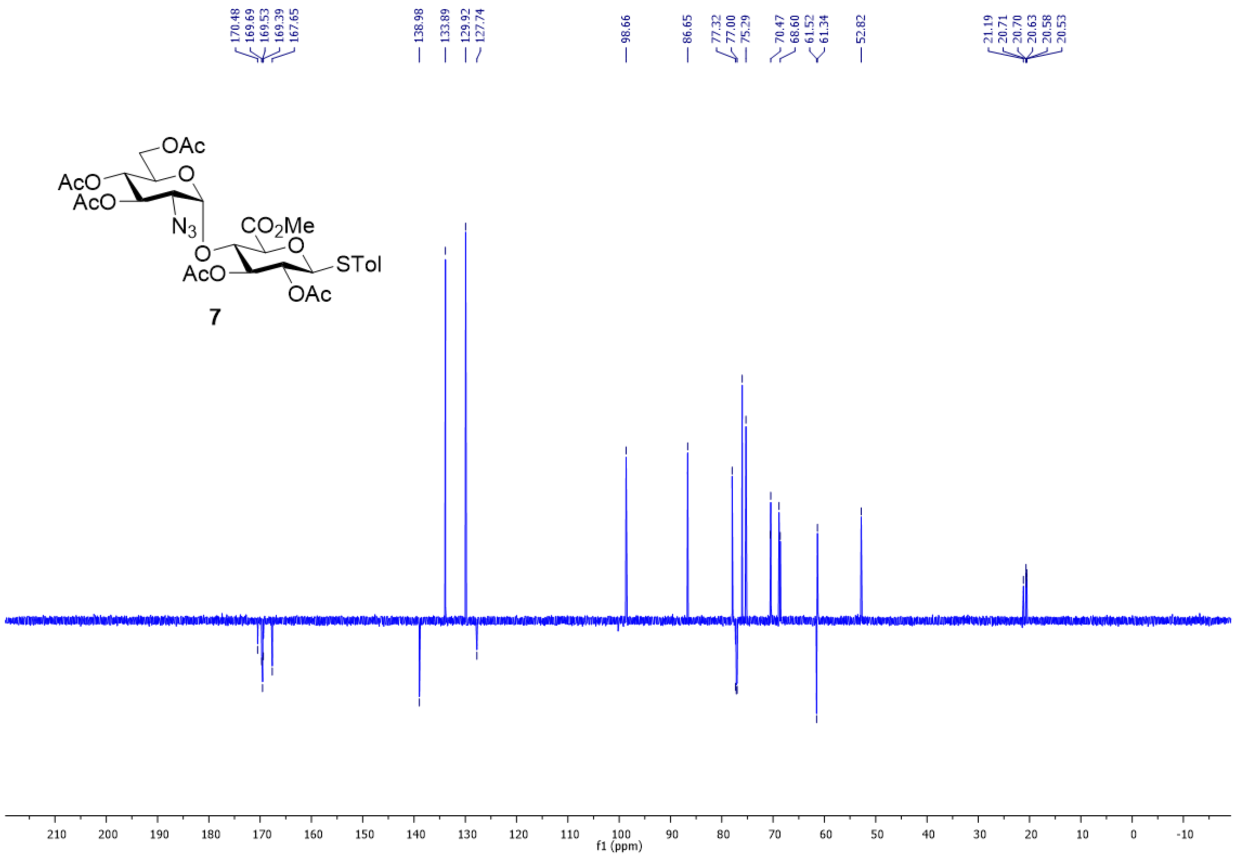


Fig. S5. $^{13}C$ NMR spectrum of **7** in $CDCl_3$.

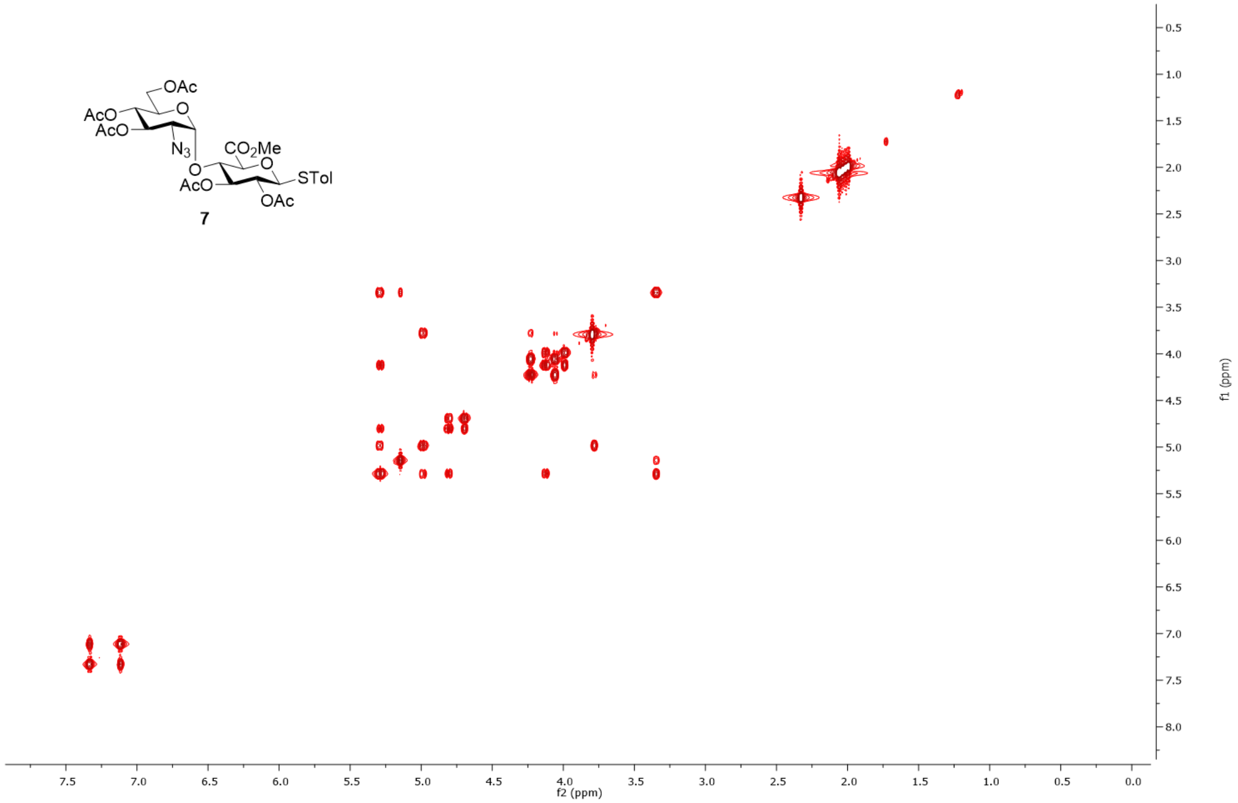


Fig. S6. $^{1}$H-$^{1}$H COSY spectrum of **7** in $CDCl_3$.

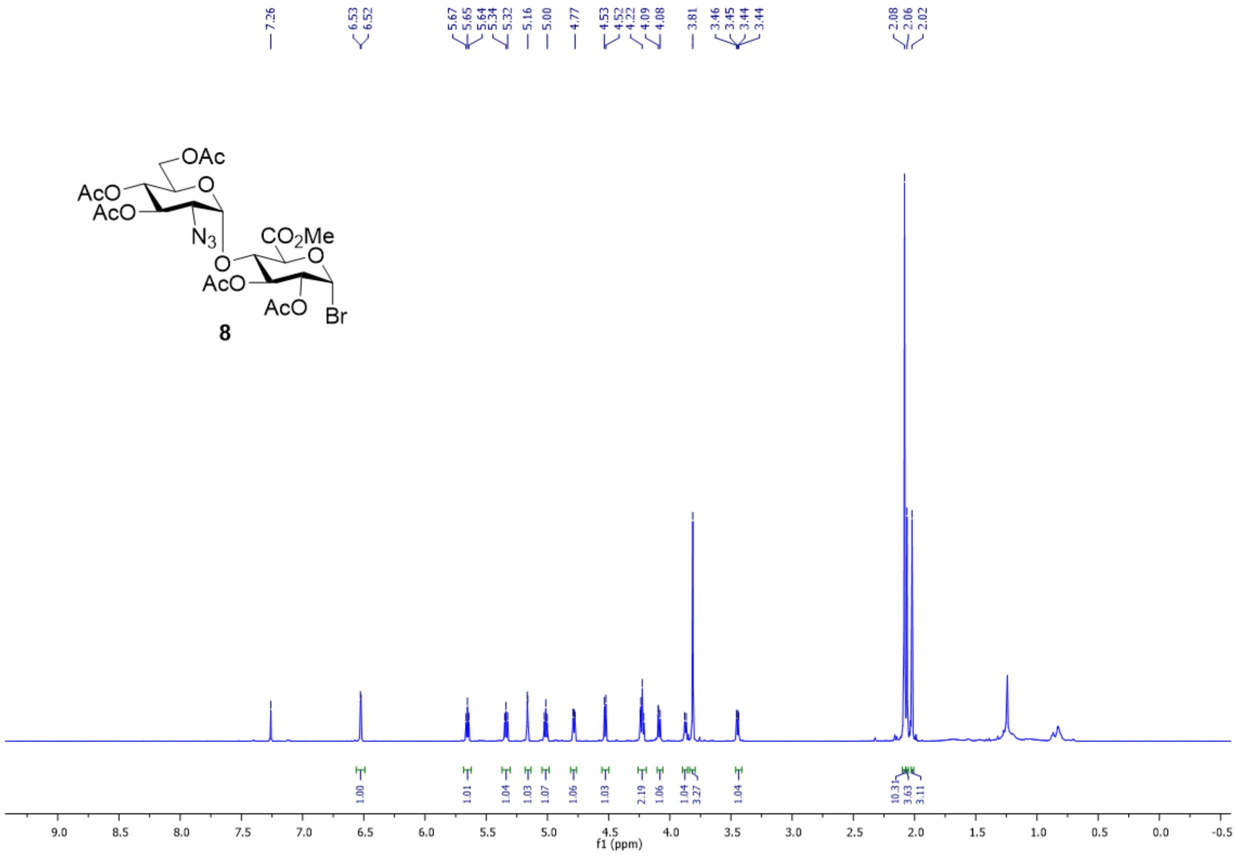

Fig. S7. $^1$H NMR spectrum of **8** in $CDCl_3$.

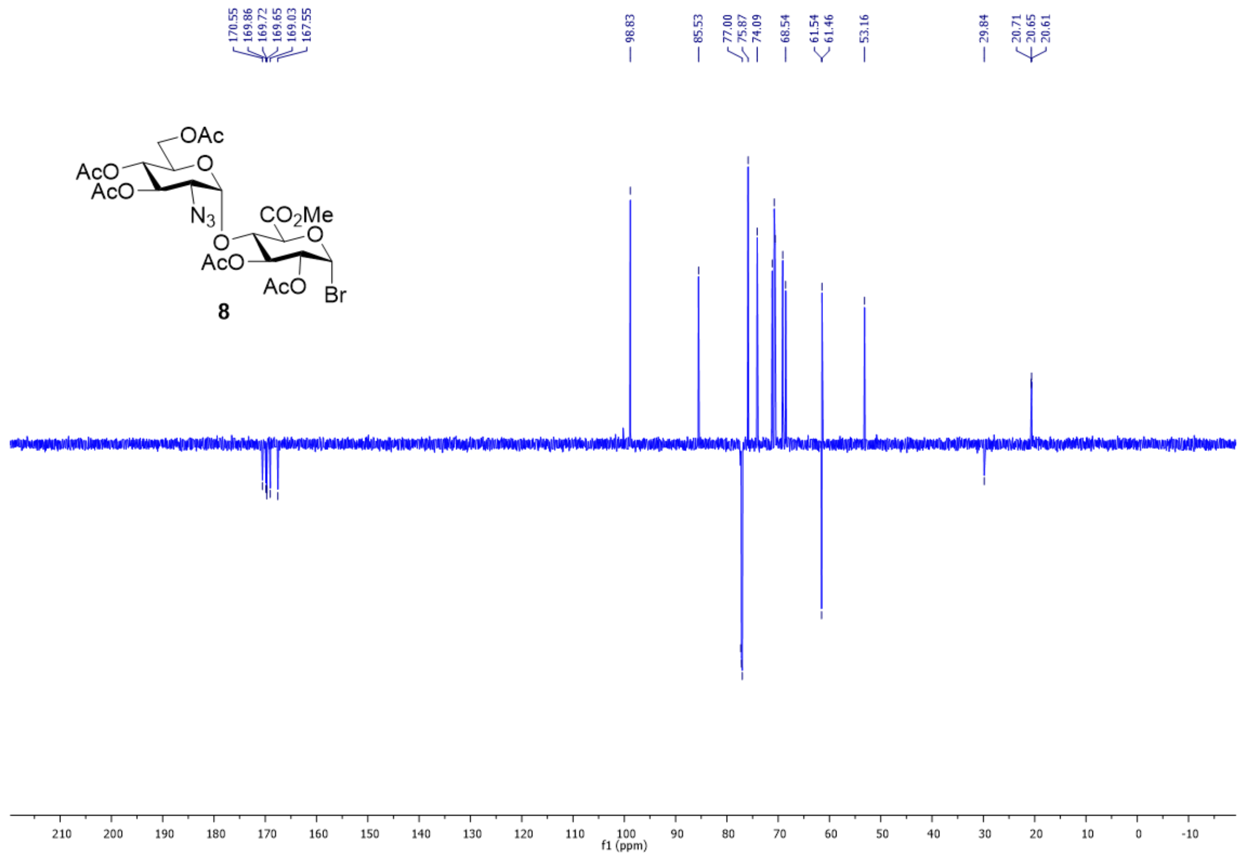


Fig. S8. $^{13}C$ NMR spectrum of **8** in $CDCl_3$.

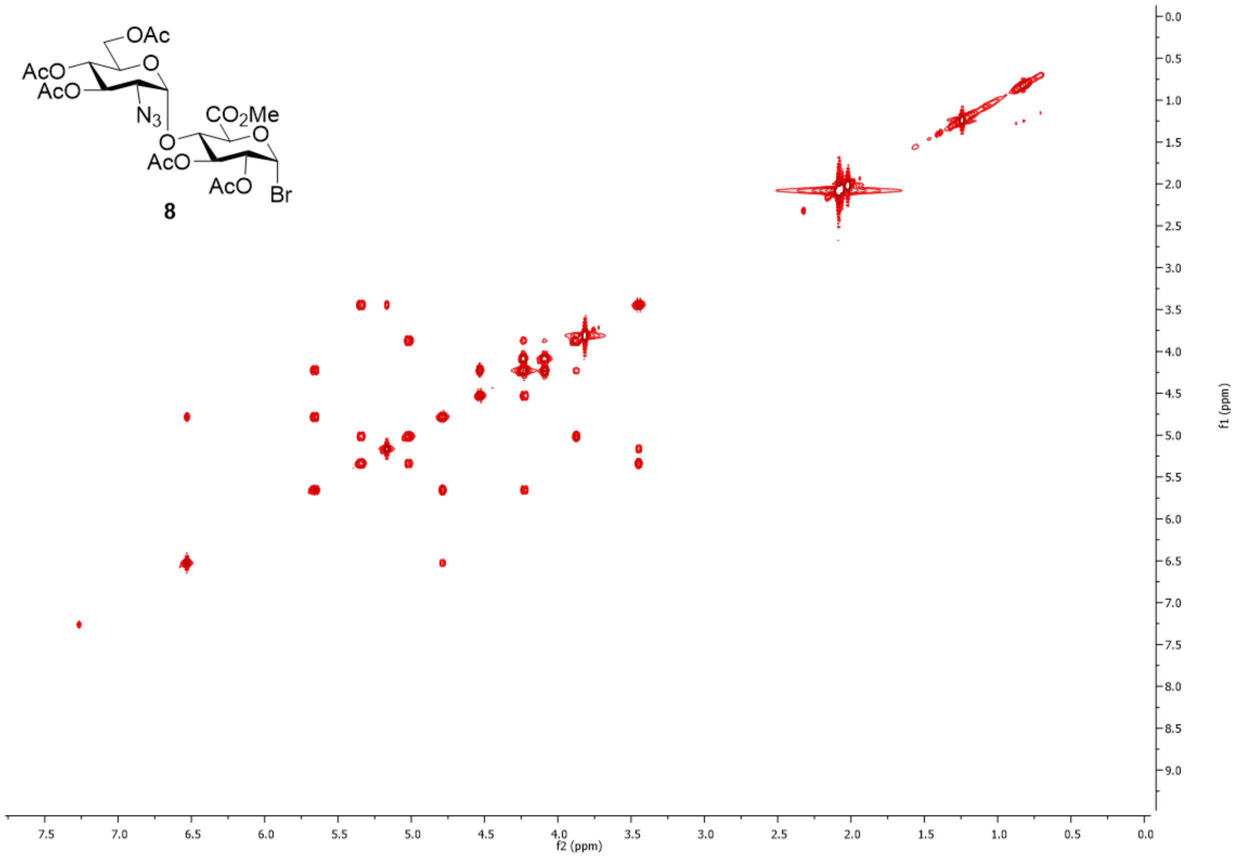


Fig. S9. $^1$H-$^1$H COSY spectrum of **8** in $CDCl_3$.

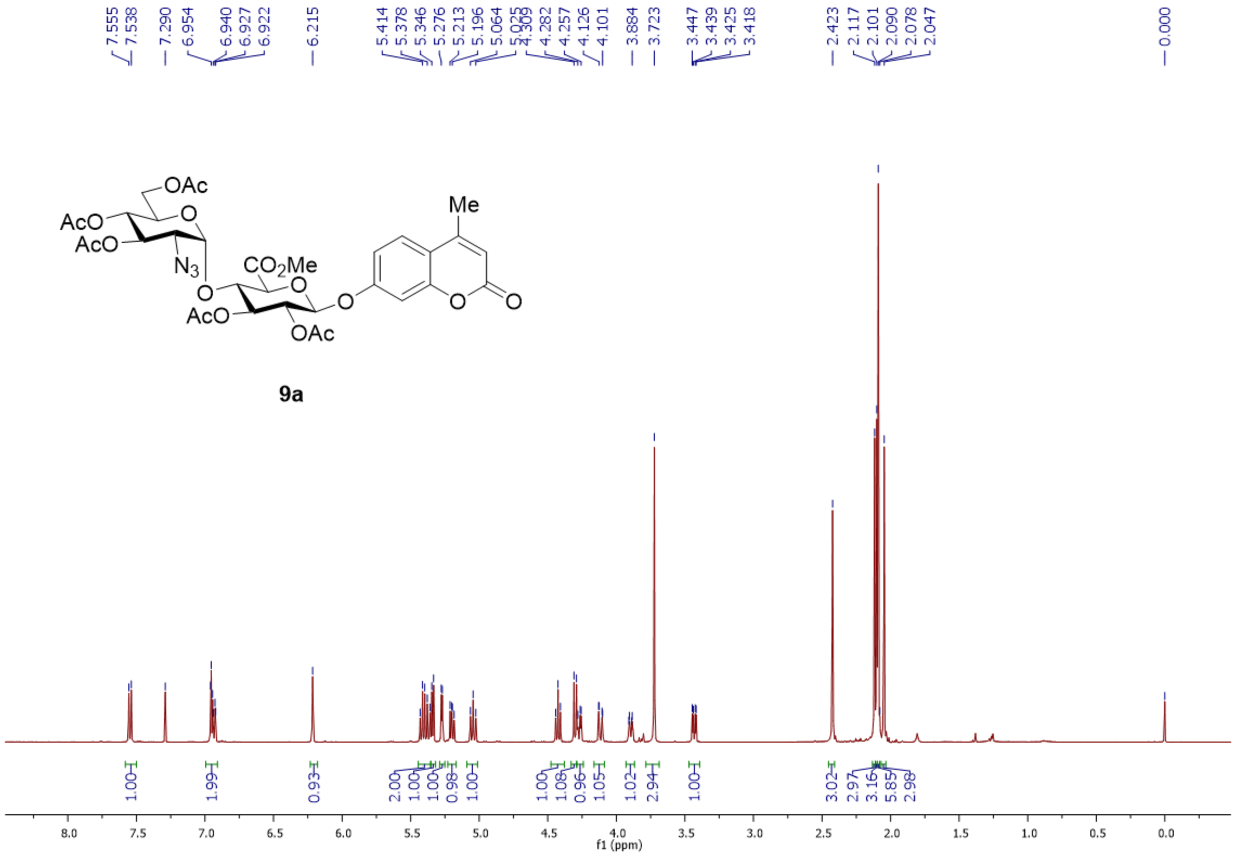

Fig. S10. $^{1}$H NMR spectrum of **9a** in $CDCl_3$.

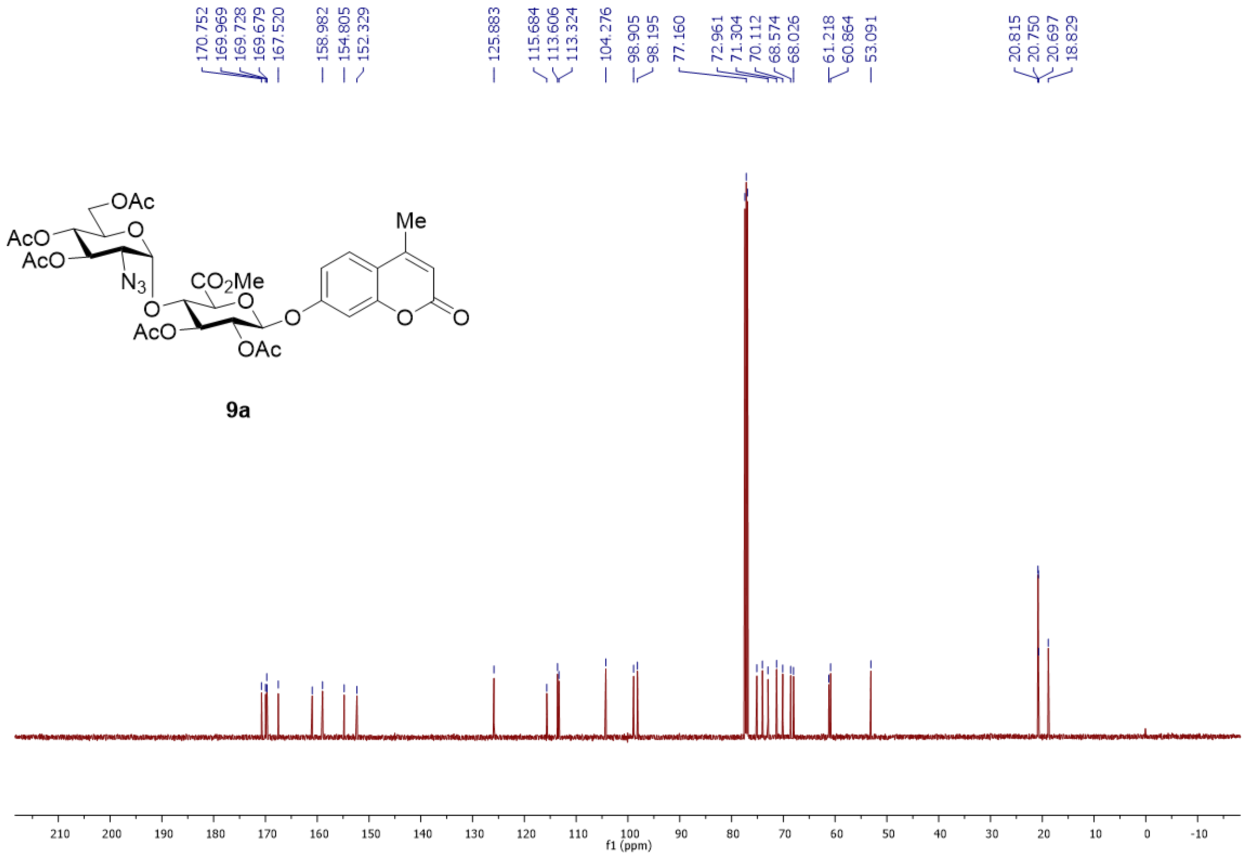


Fig. S11. $^{13}$C NMR spectrum of **9a** in $CDCl_3$.

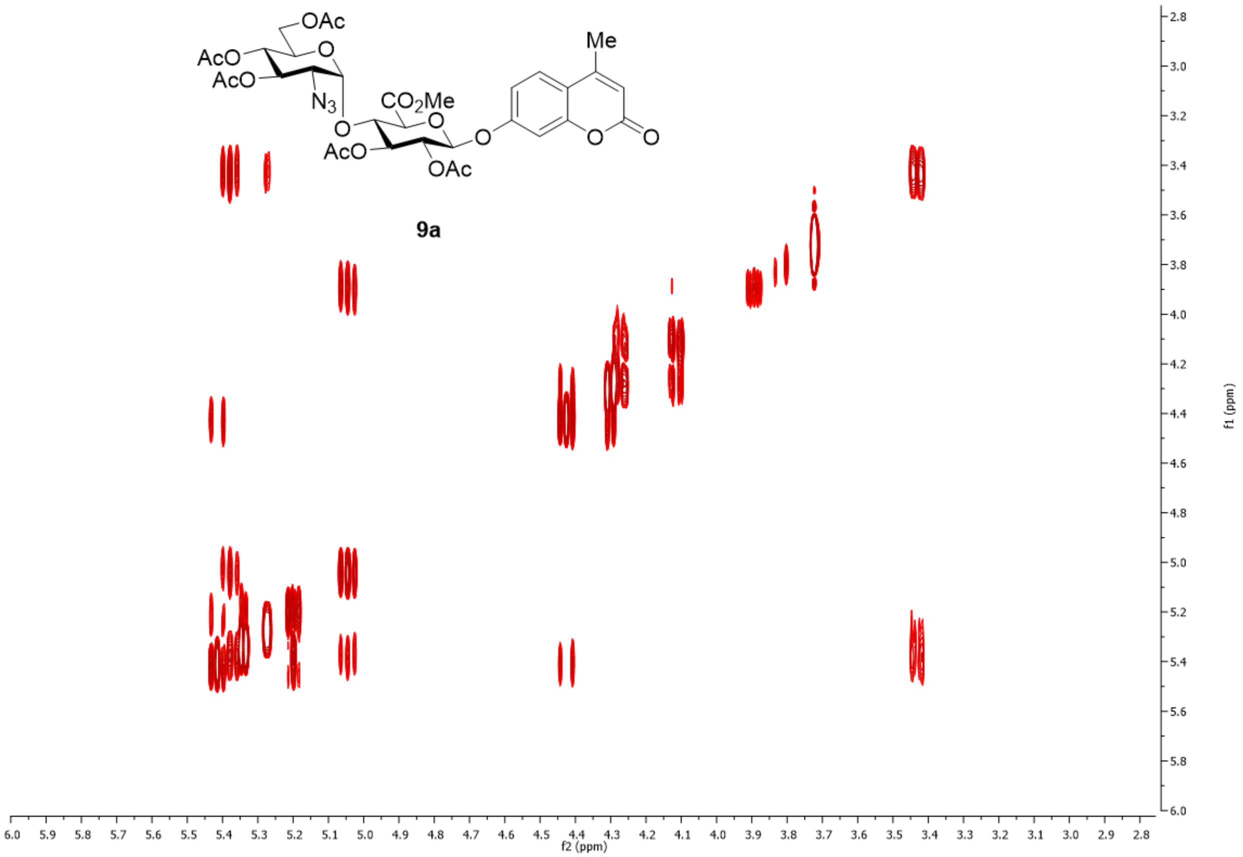


Fig. S12. $^{1}H$-$^{1}H$ COSY spectrum of **9a** in $CDCl_3$.

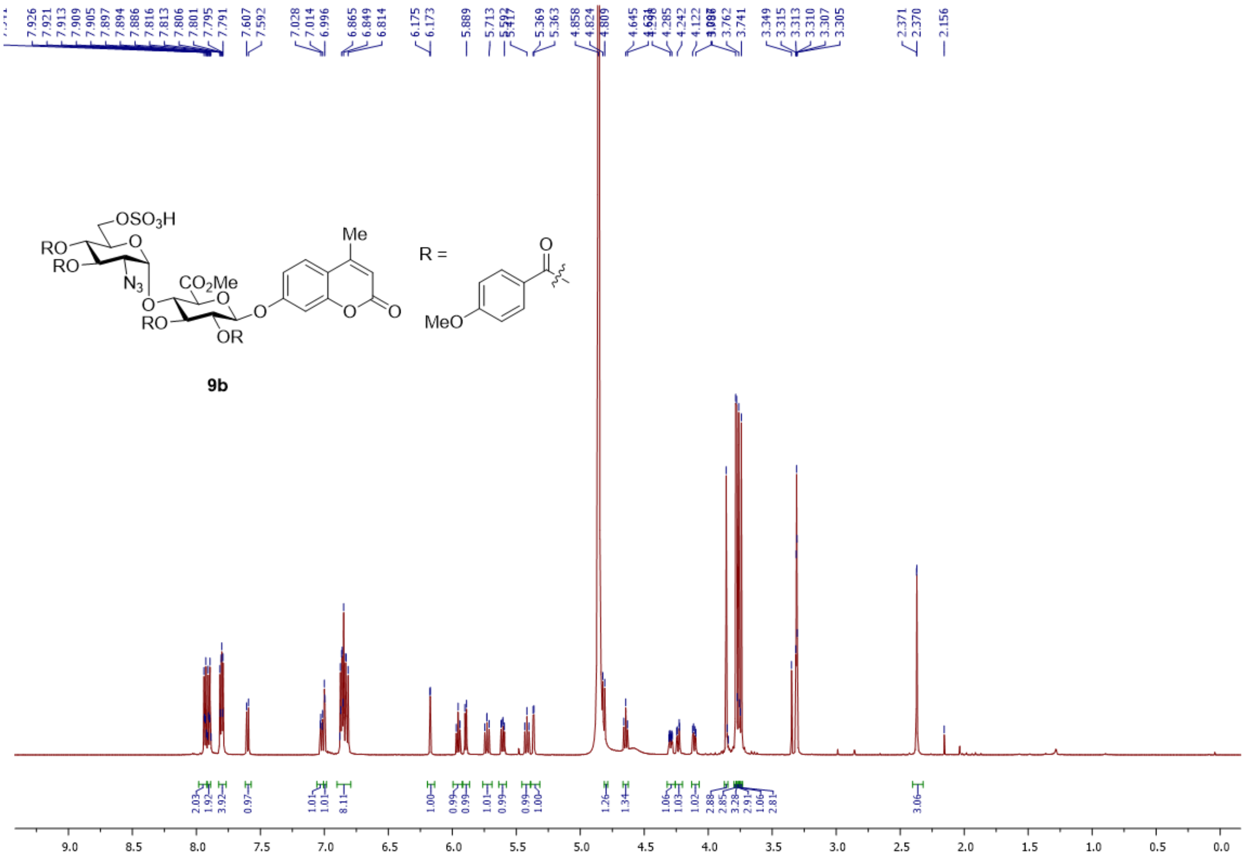

Fig. S13. $^{1}H$ NMR spectrum of **9b** in $CDCl_3$.

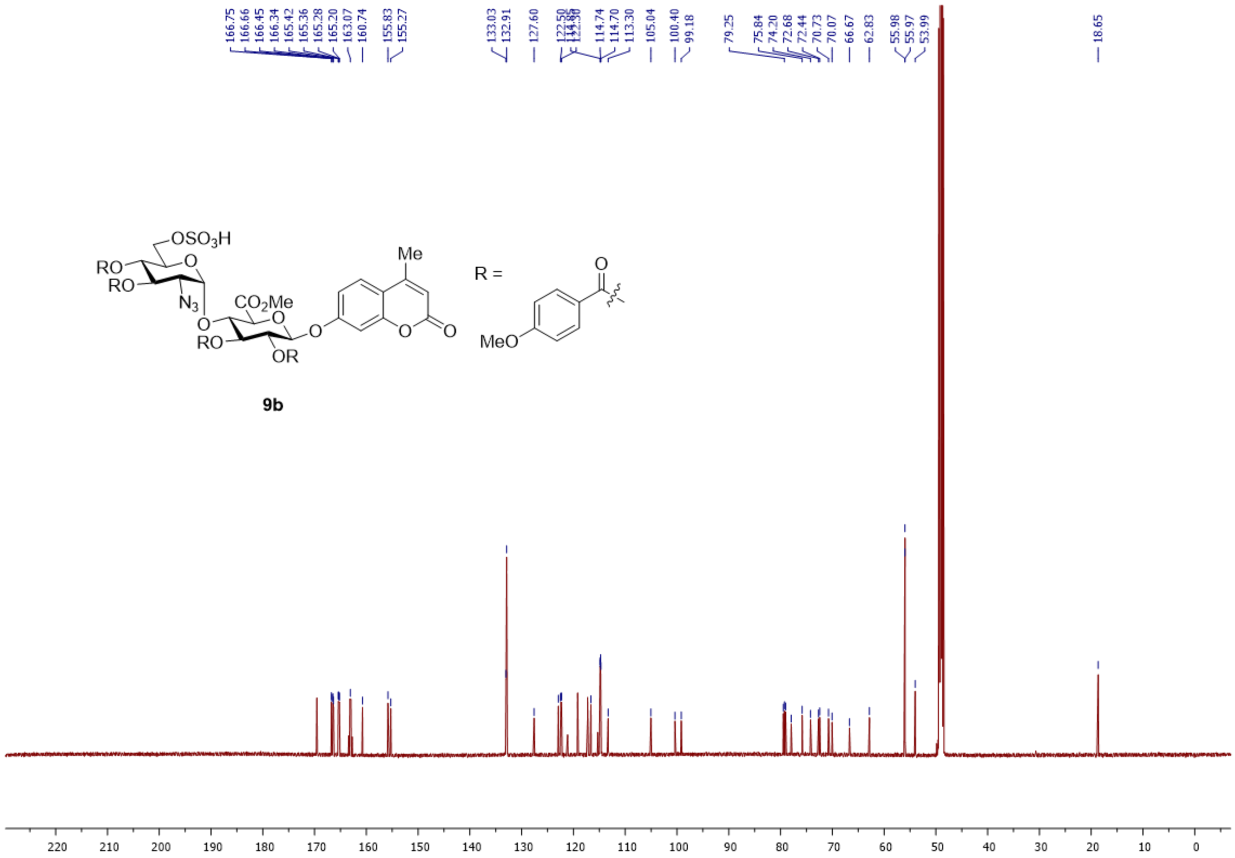

Fig. S14. $^{13}$C NMR spectrum of **9b** in $CDCl_3$.

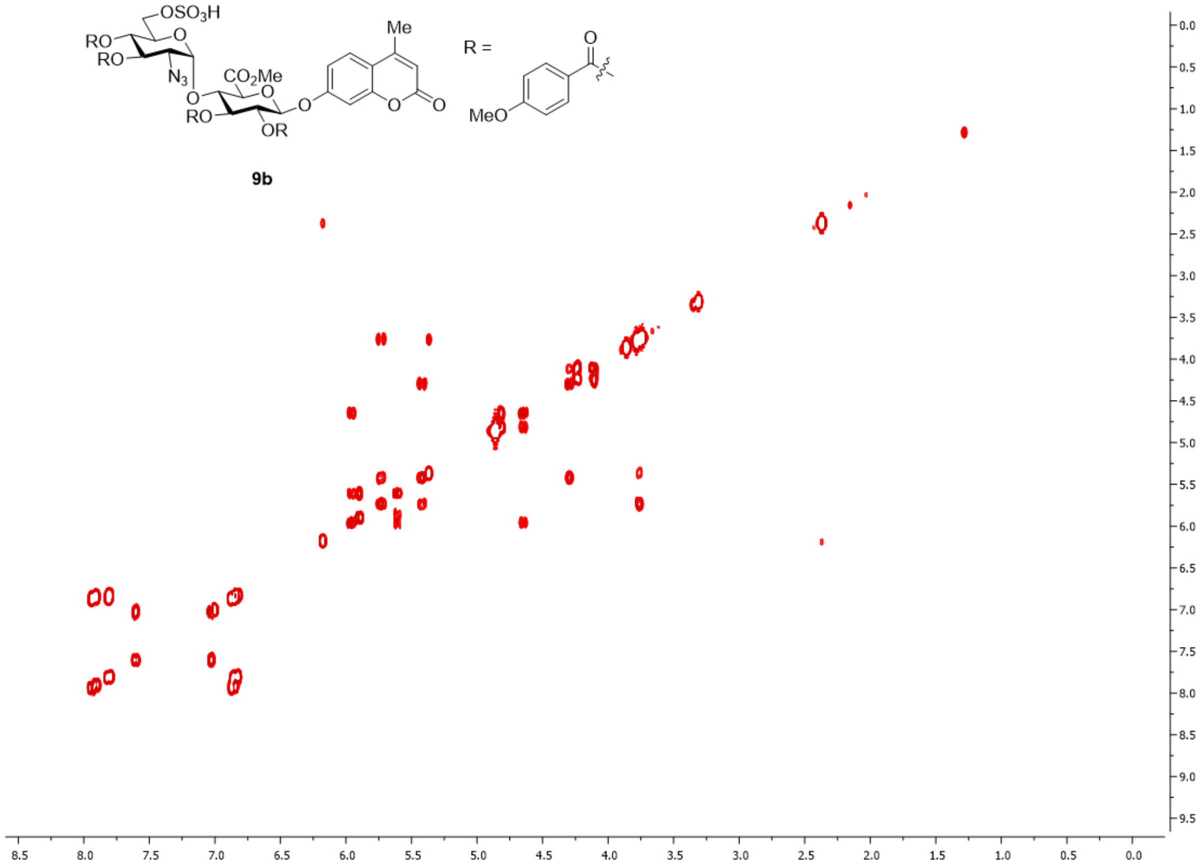


Fig. S15. $^{1}H$-$^{1}H$ COSY spectrum of **9b** in $CDCl_3$.

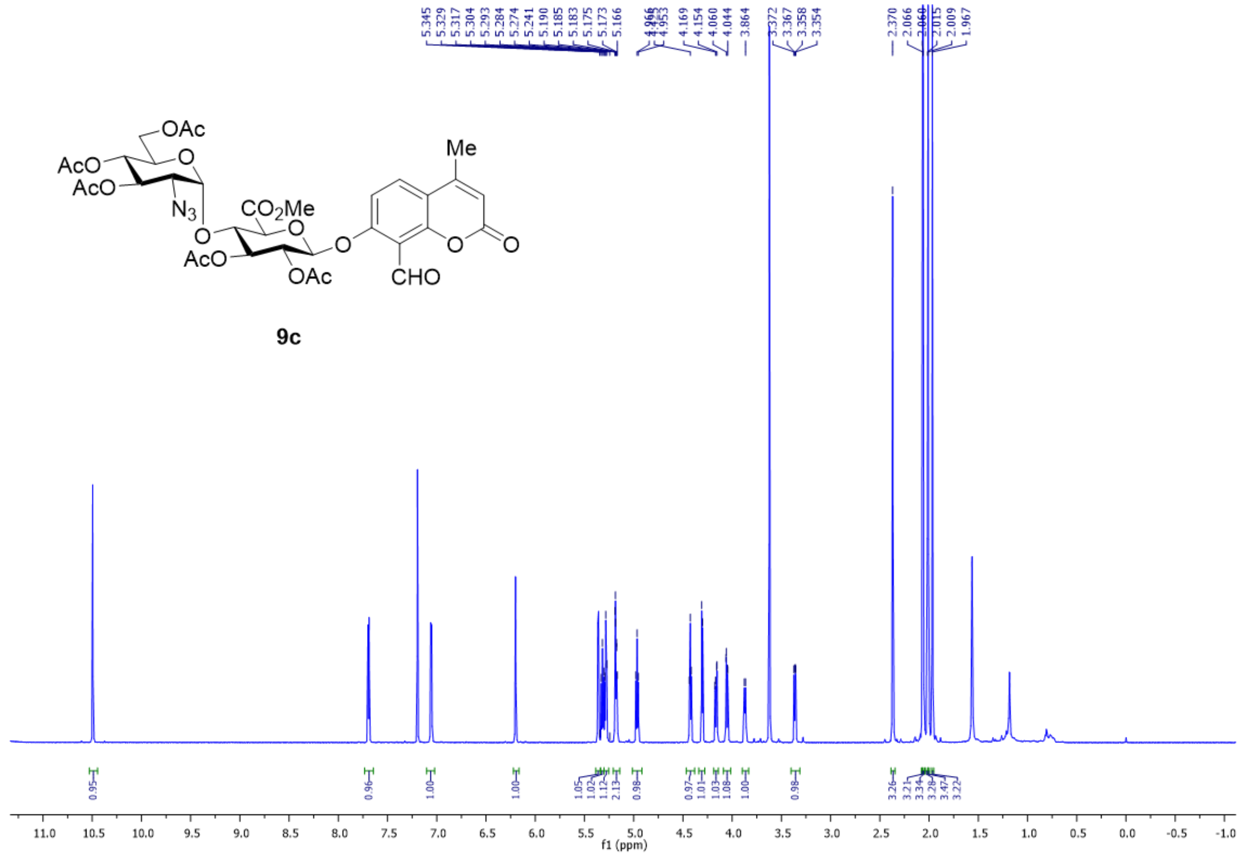

Fig. S16. $^{1}$H NMR spectrum of **9c** in $CDCl_3$.

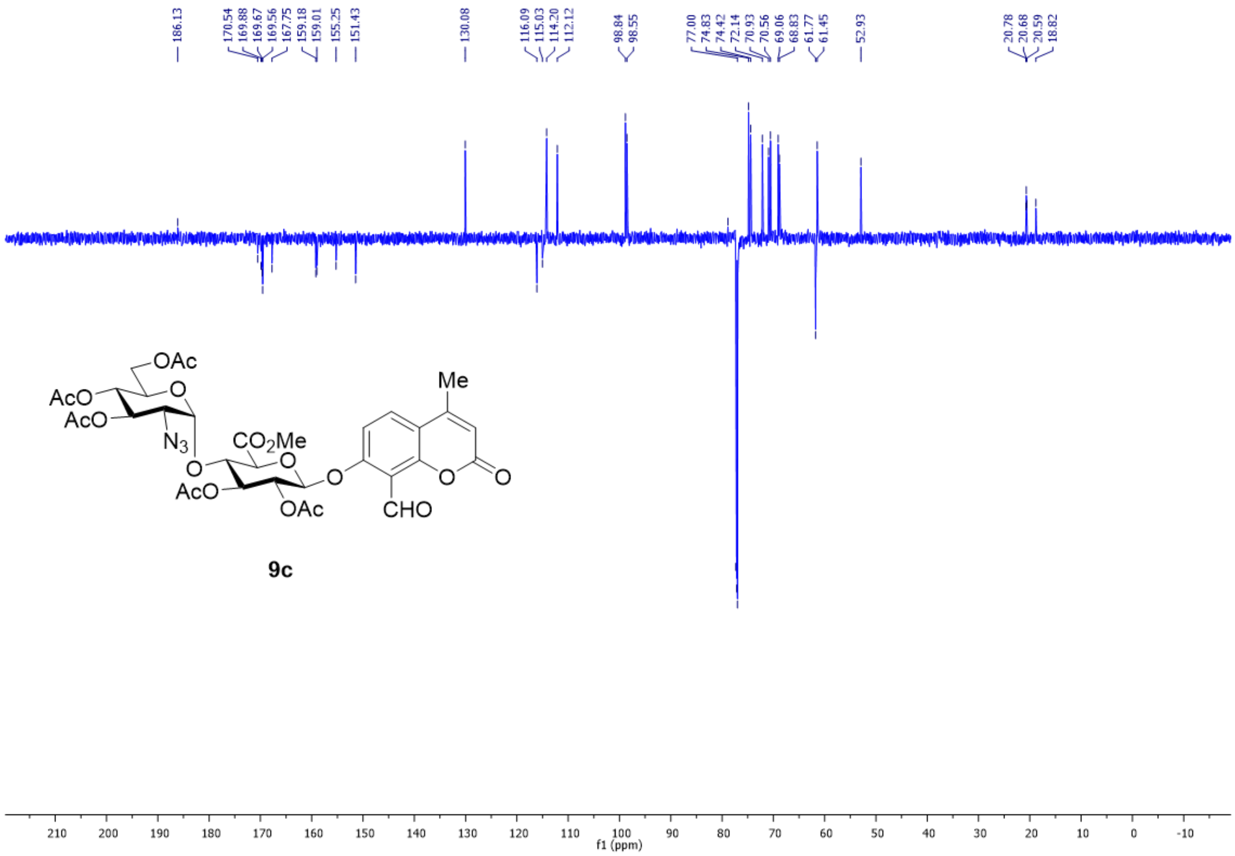

Fig. S17. $^{13}$C NMR spectrum of **9c** in $CDCl_3$.

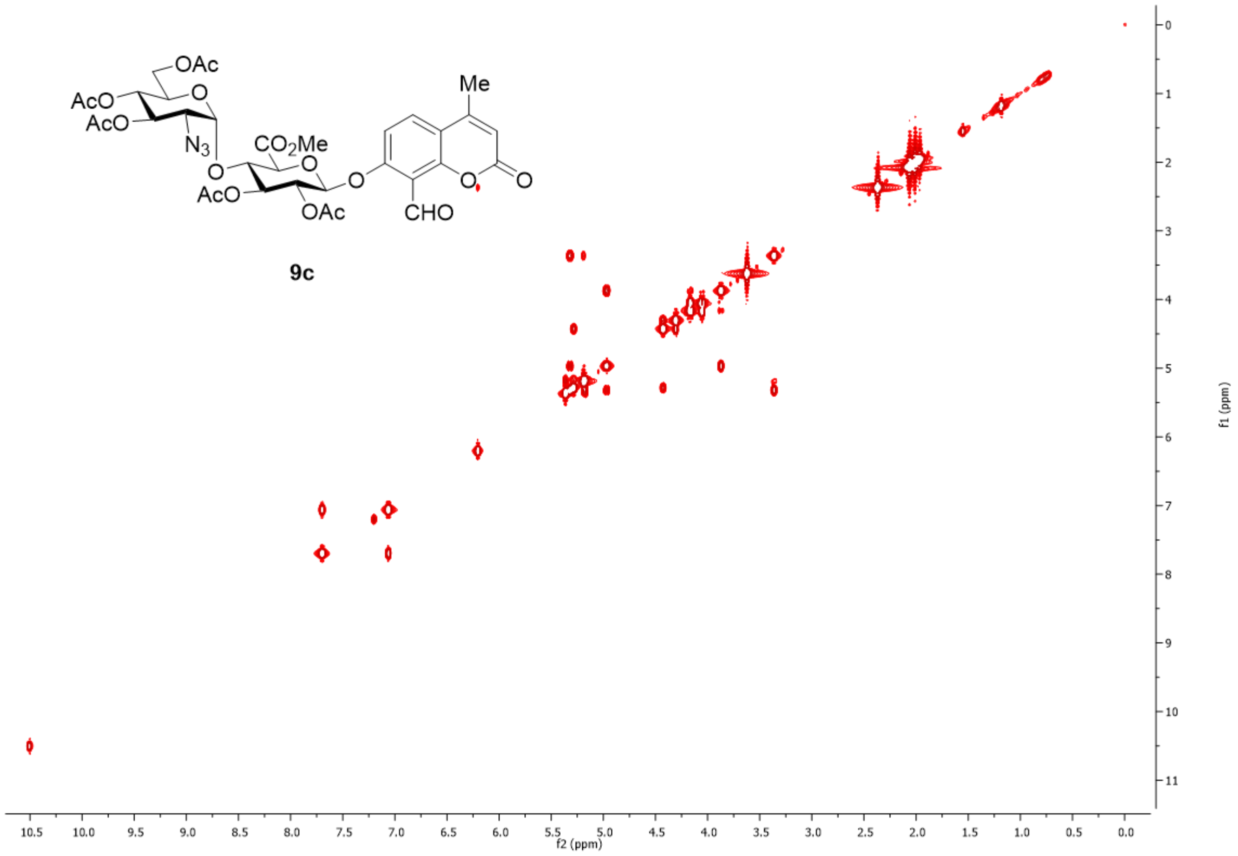


Fig. S18. $^{1}H$-$^{1}H$ COSY spectrum of **9c** in $CDCl_3$.

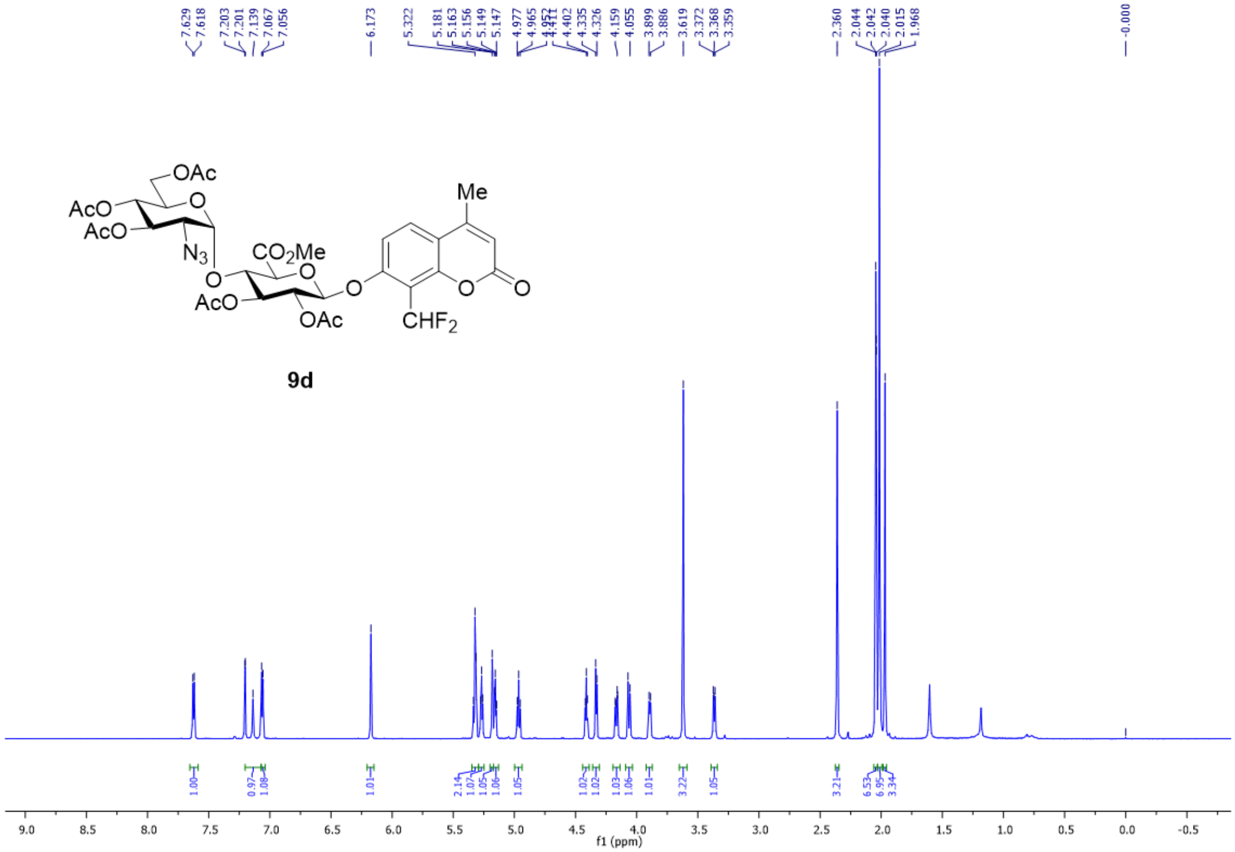

Fig. S19. $^{1}H$ NMR spectrum of **9d** in $CDCl_3$.

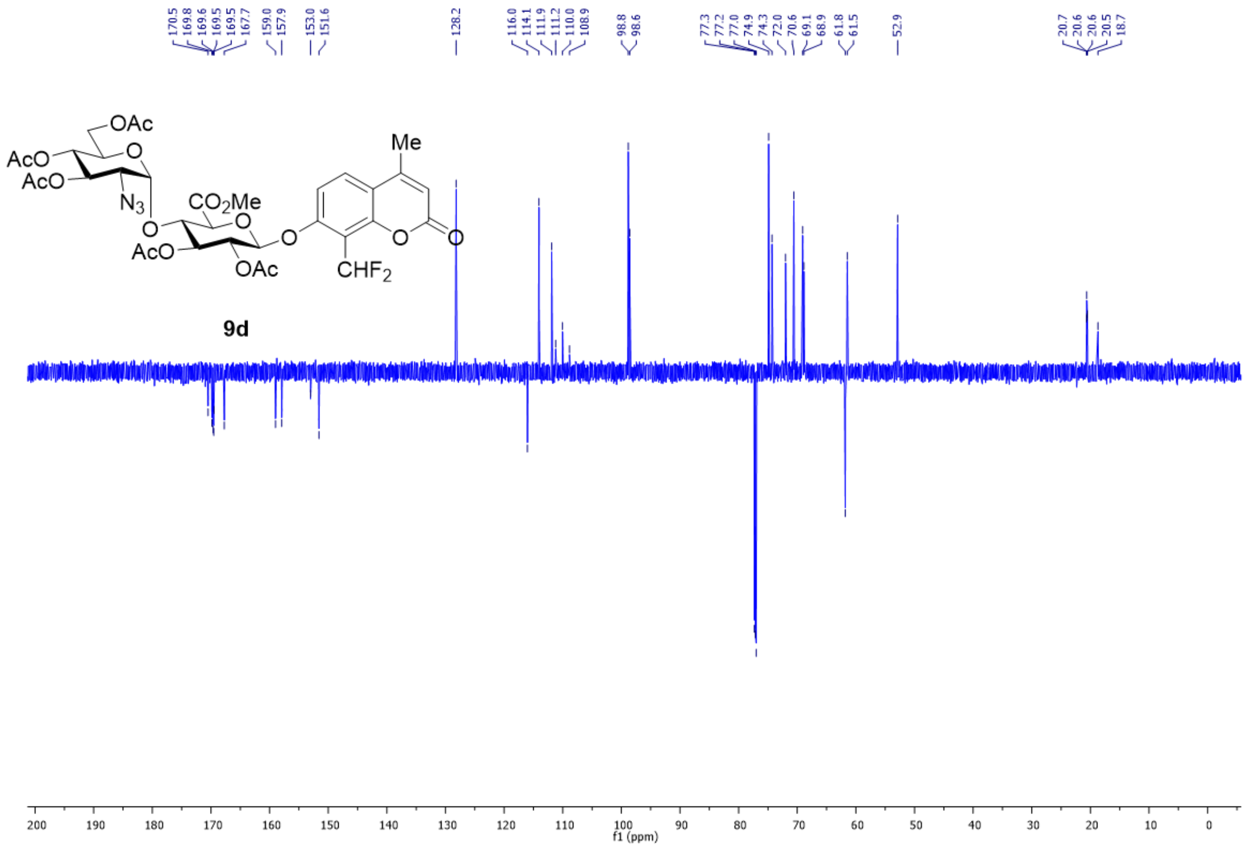

Fig. S20. $^{13}$C NMR spectrum of **9d** in $CDCl_3$.

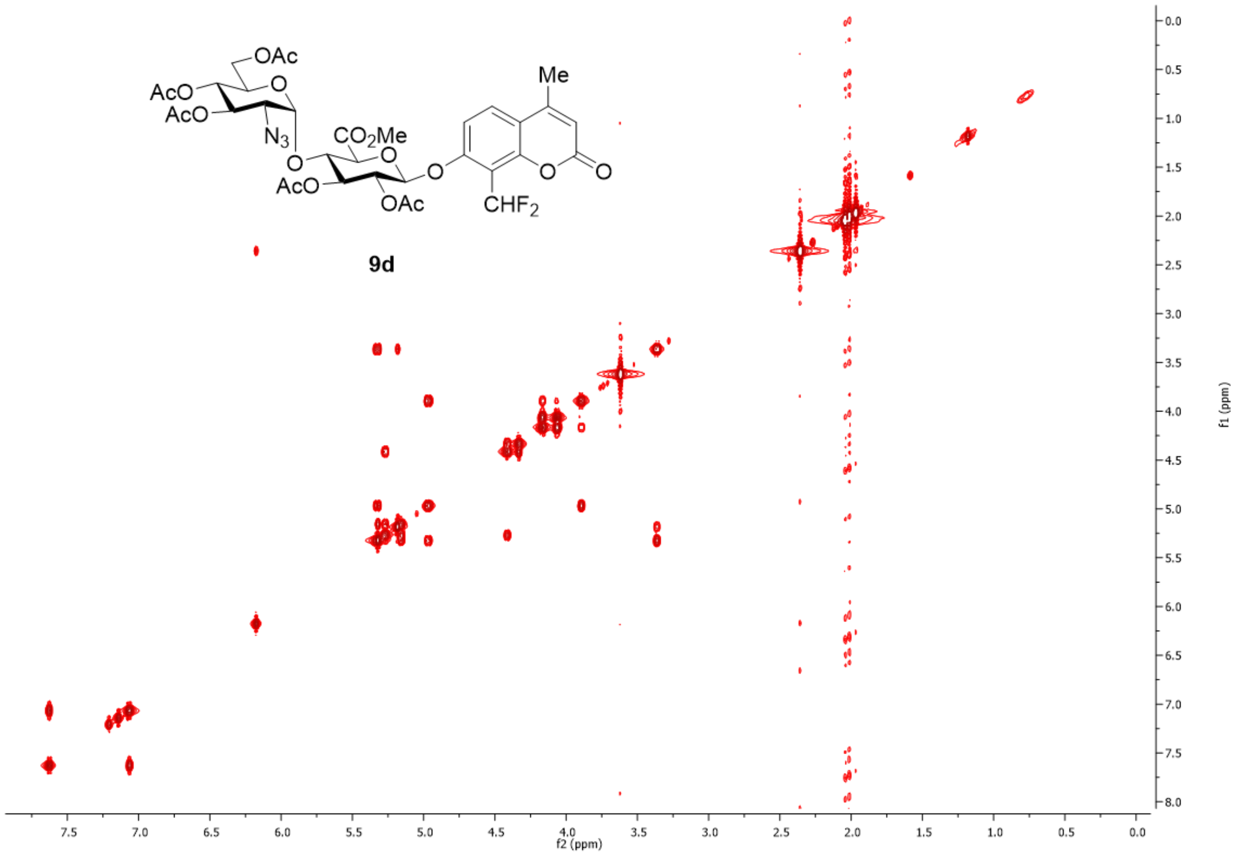


Fig. S21. $^1$H-$^1$H COSY spectrum of **9d** in $CDCl_3$.

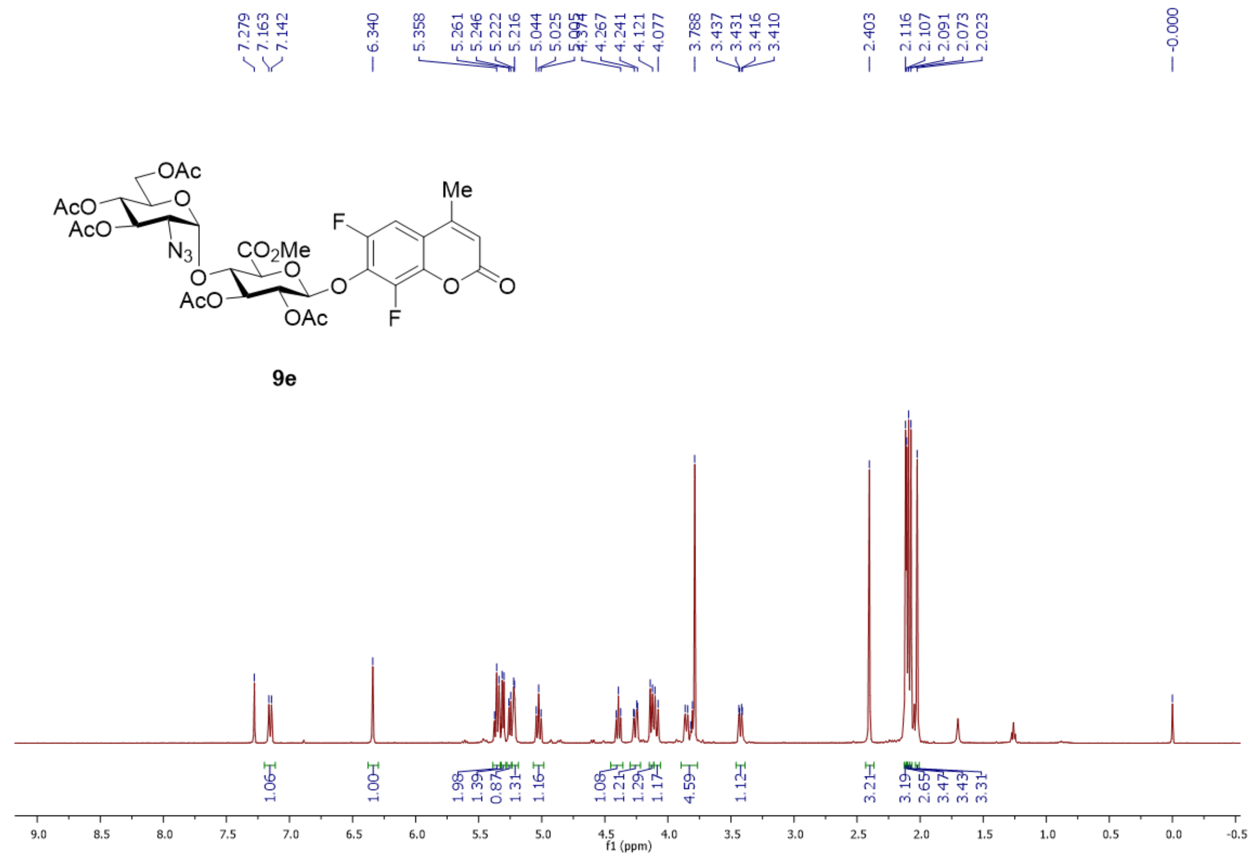

Fig. S22. $^{1}H$ NMR spectrum of **9e** in $CDCl_3$.

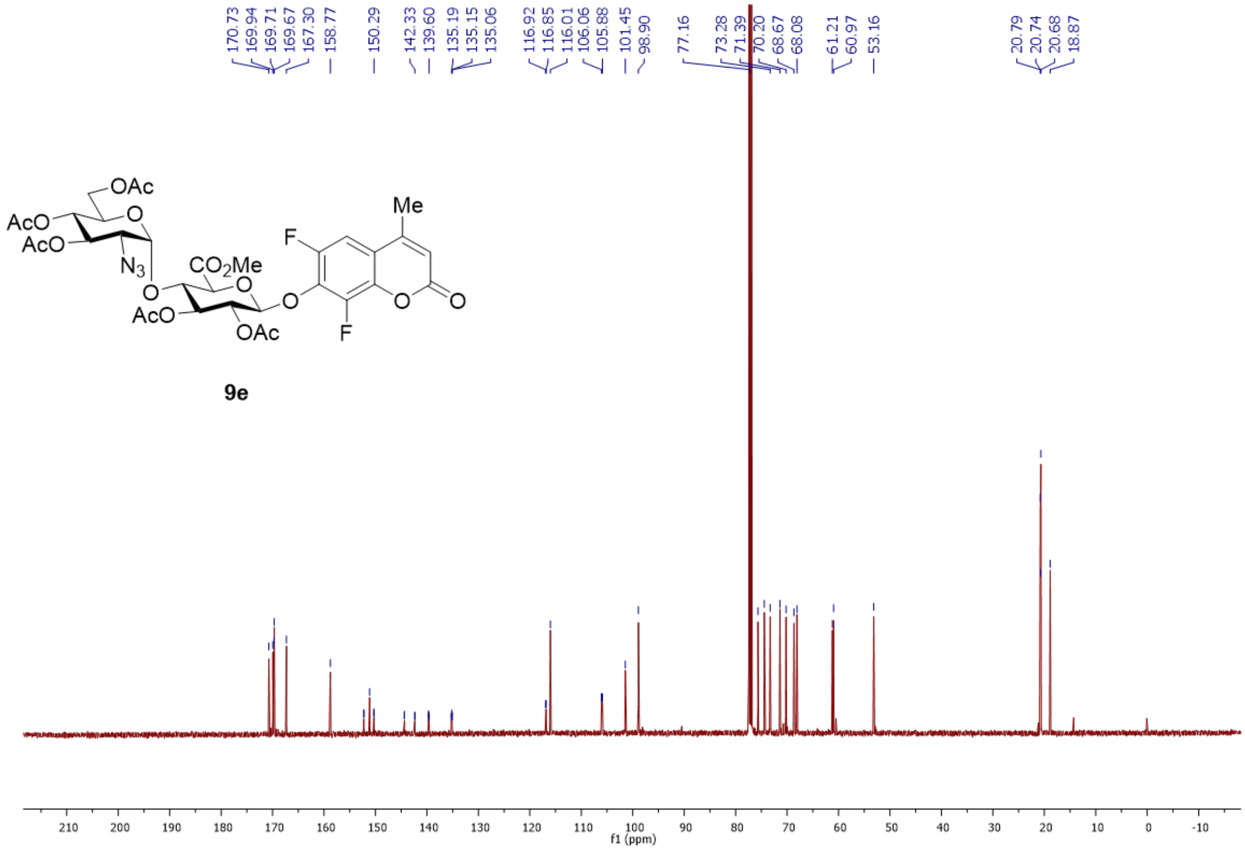


Fig. S23. $^{13}$C NMR spectrum of **9e** in $CDCl_3$.

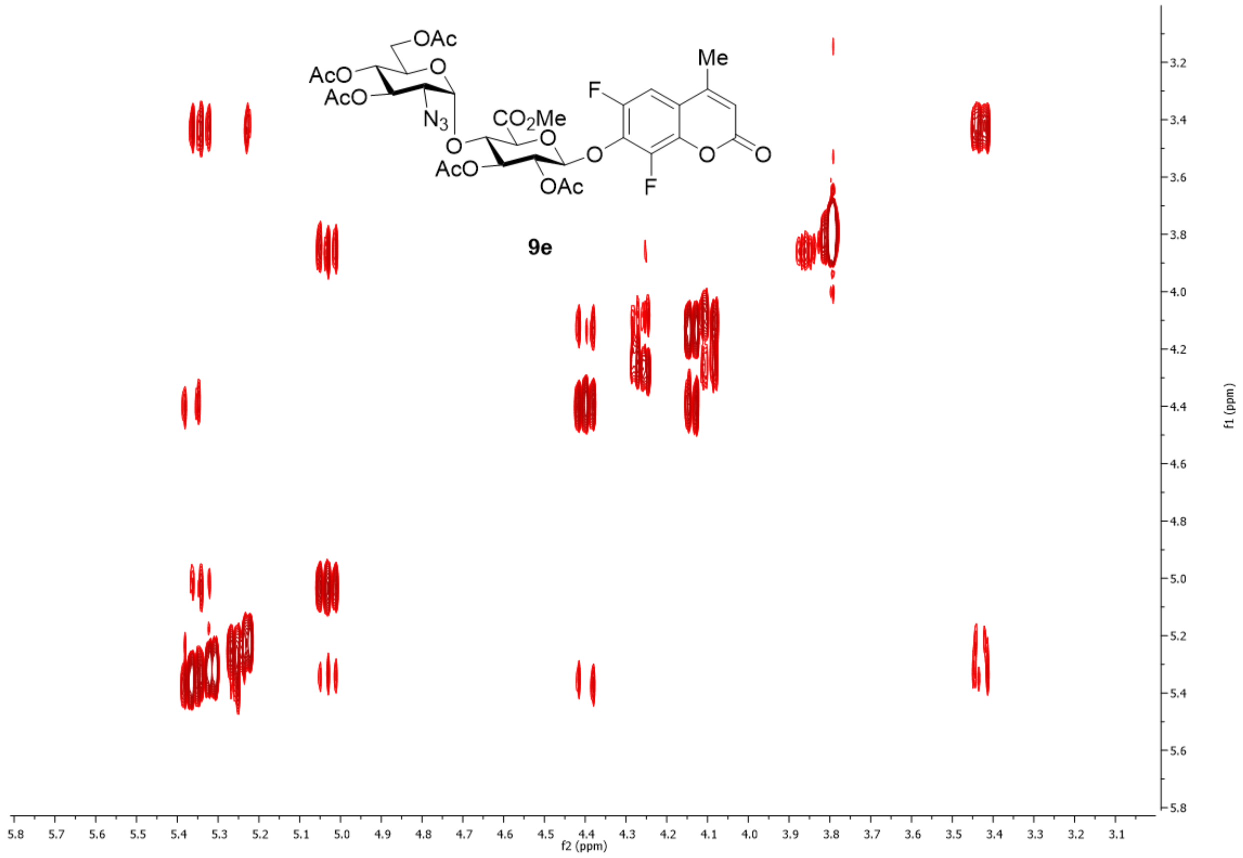


Fig. S24. $^{1}$H-$^{1}$H COSY spectrum of **9e** in $CDCl_3$.

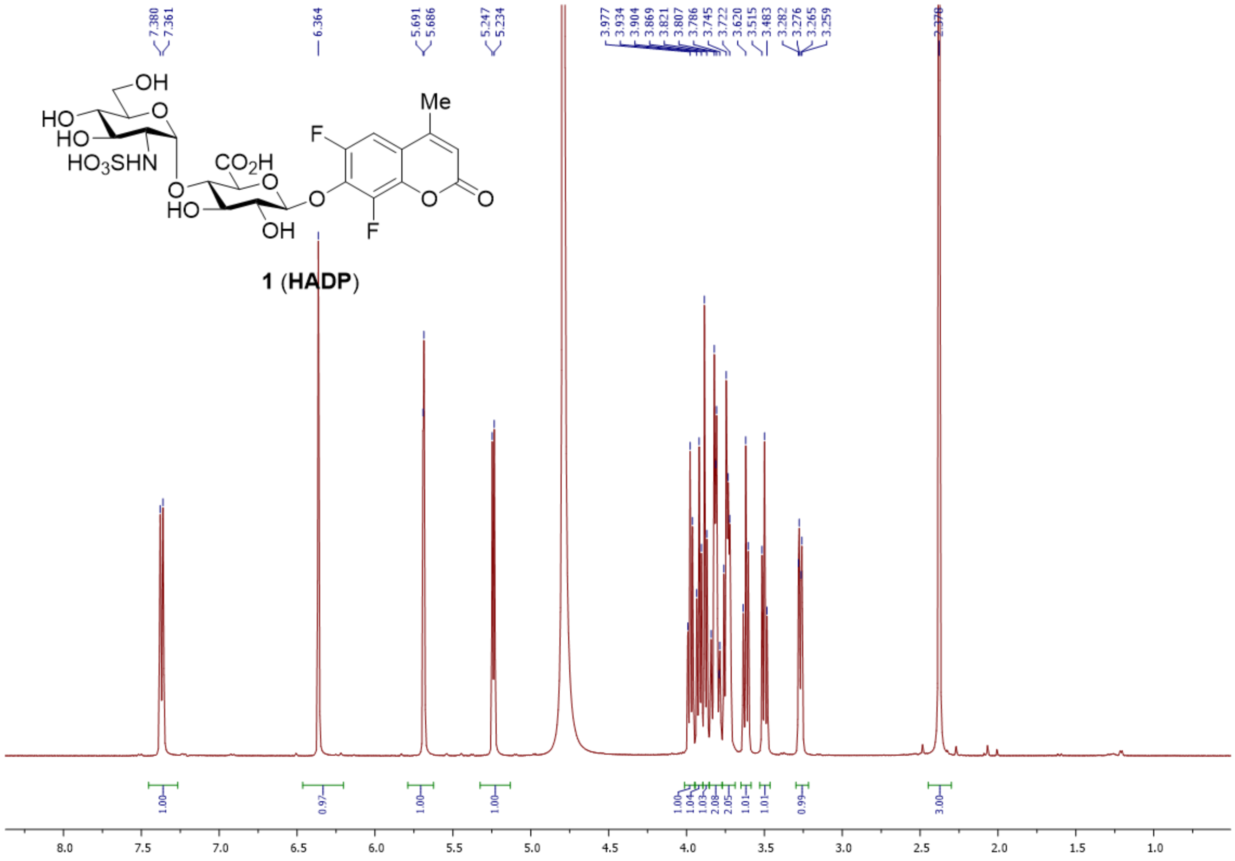

Fig. S25. $^{1}$H NMR spectrum of **1** in $D_2O$.

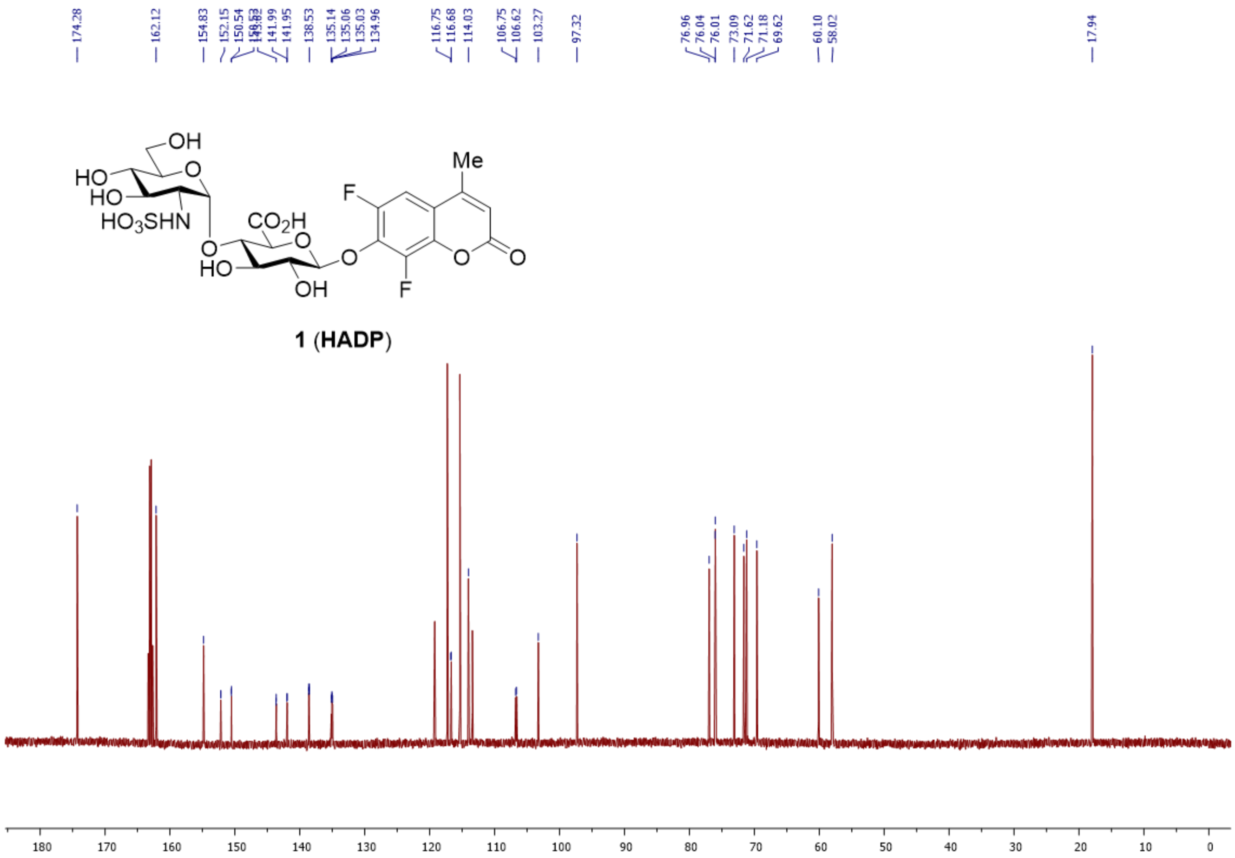


Fig. S26. $^{13}C$ NMR spectrum of **1** in $D_2O$.

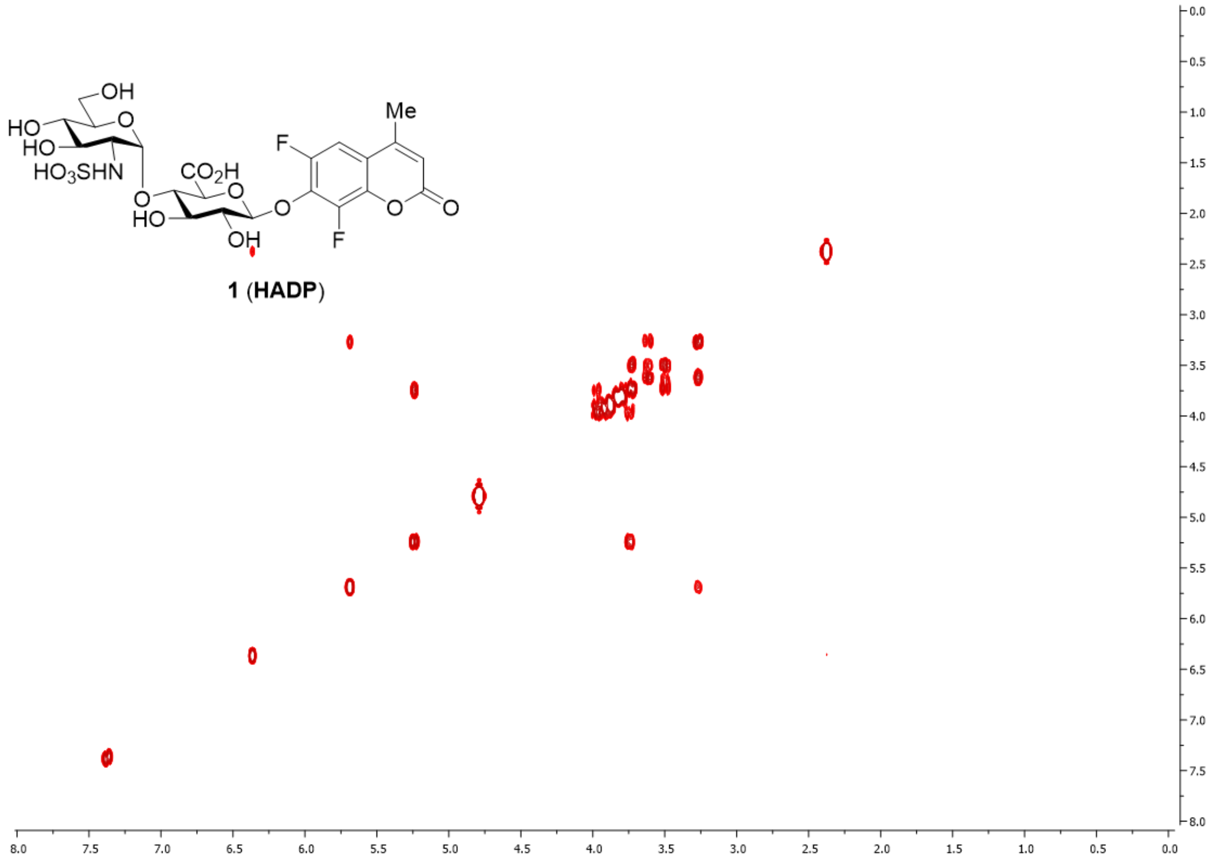


Fig. S27. $^{1}H$-$^{1}H$ COSY spectrum of **1** in $D_2O$.

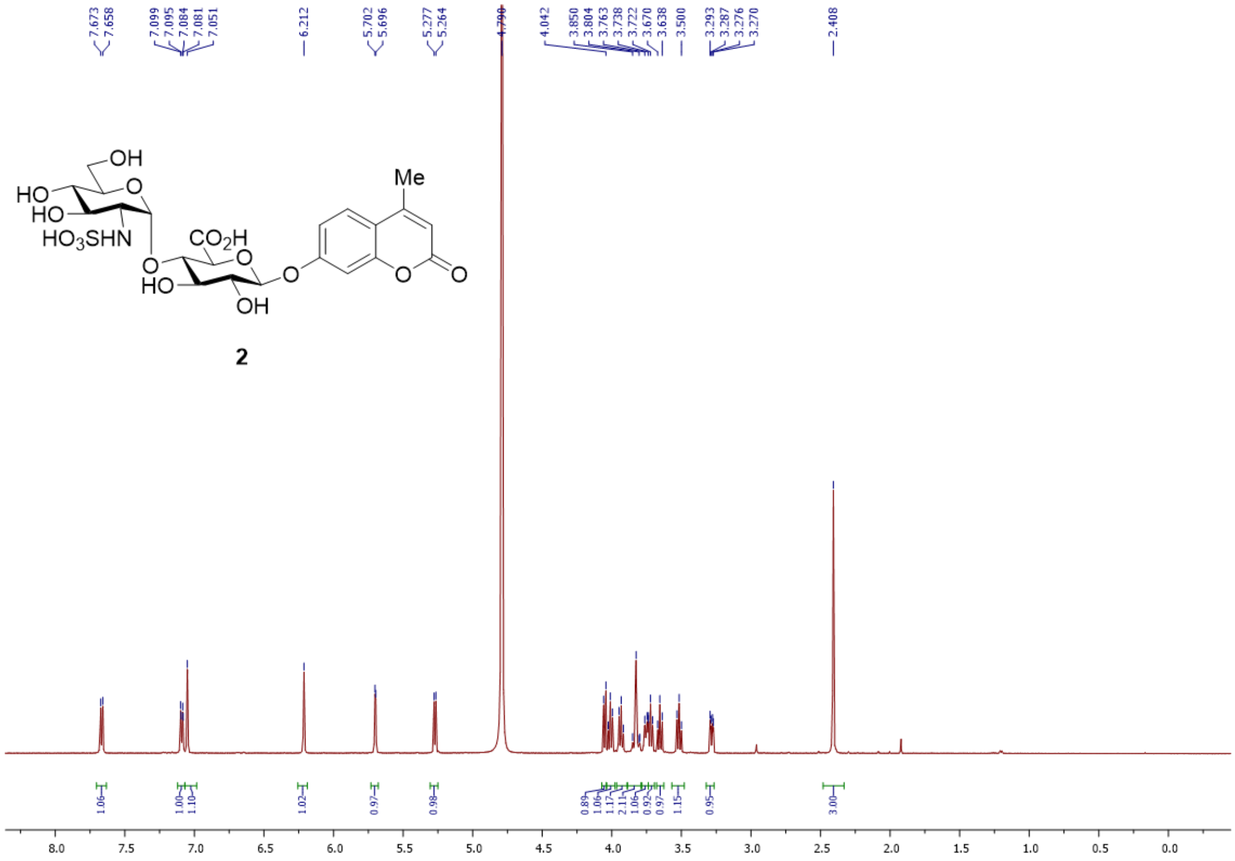

Fig. S28. $^{1}H$ NMR spectrum of **2** in $D_2O$.

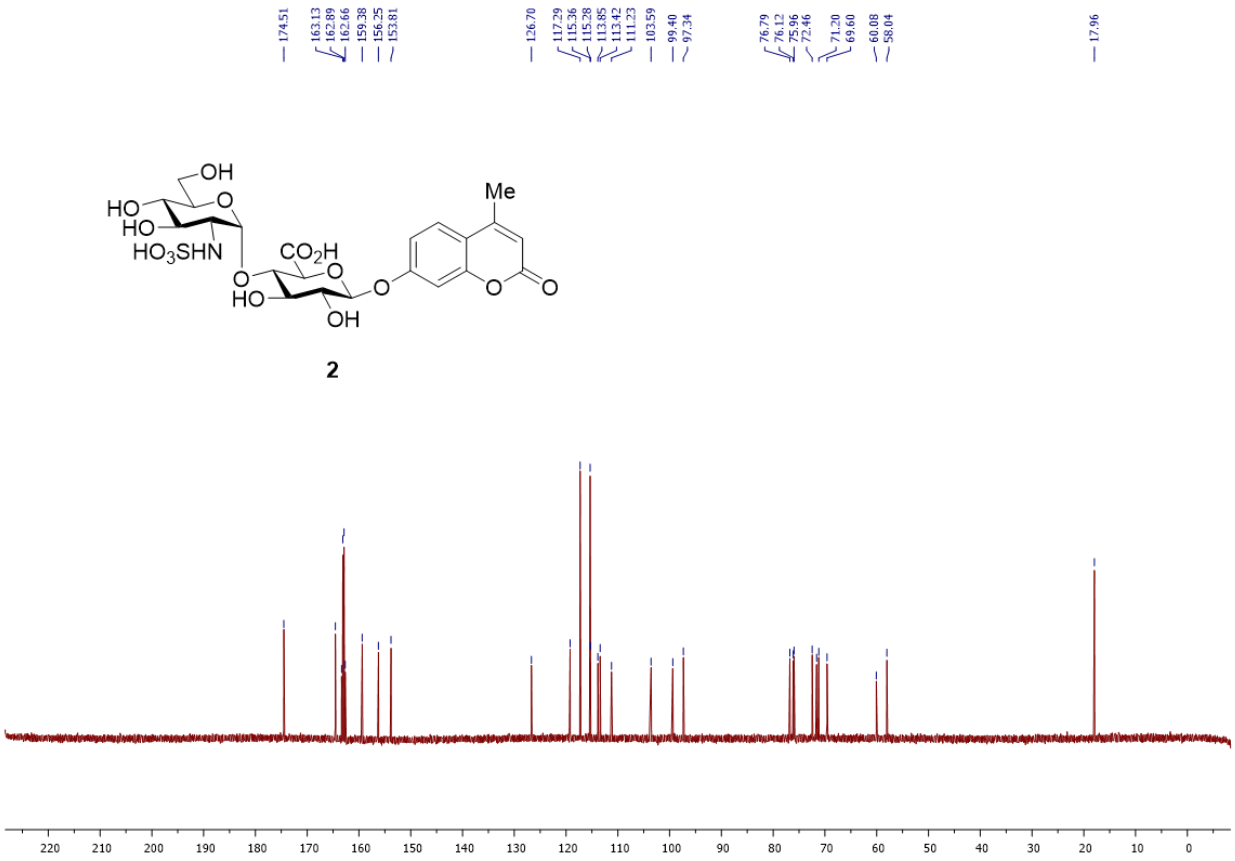

Fig. S29. $^{13}C$ NMR spectrum of **2** in $D_2O$.

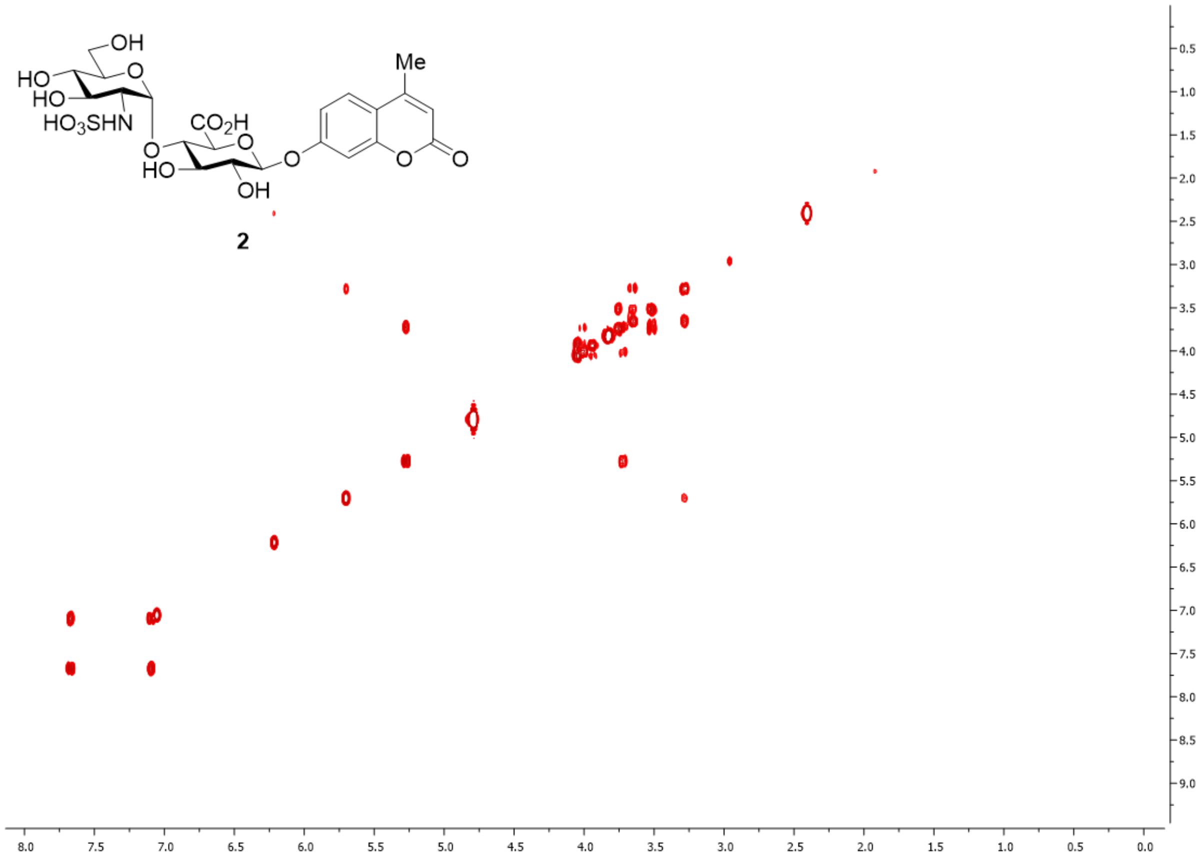


Fig. S30. $^{1}H$-$^{1}H$ COSY spectrum of **2** in $D_2O$.

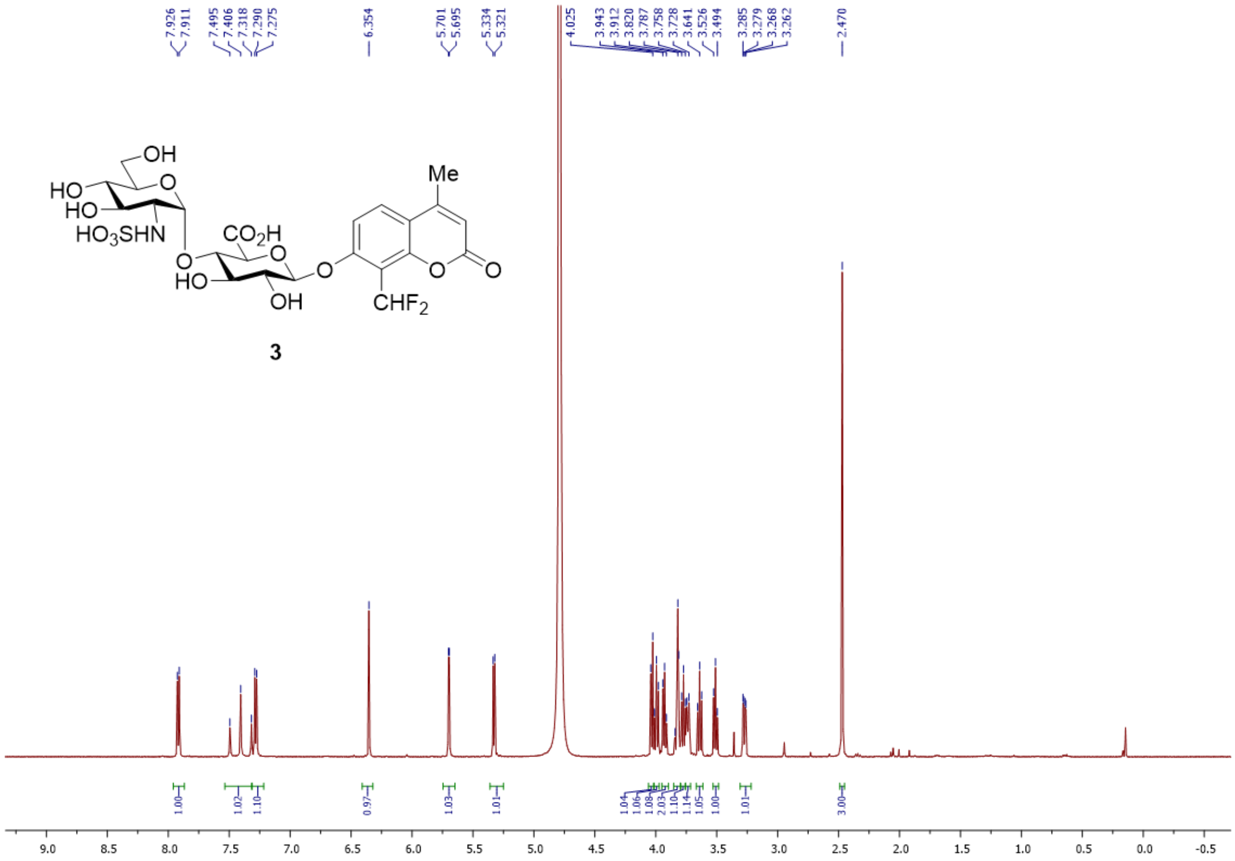

Fig. S31. $^{1}$H NMR spectrum of **3** in $D_2O$.

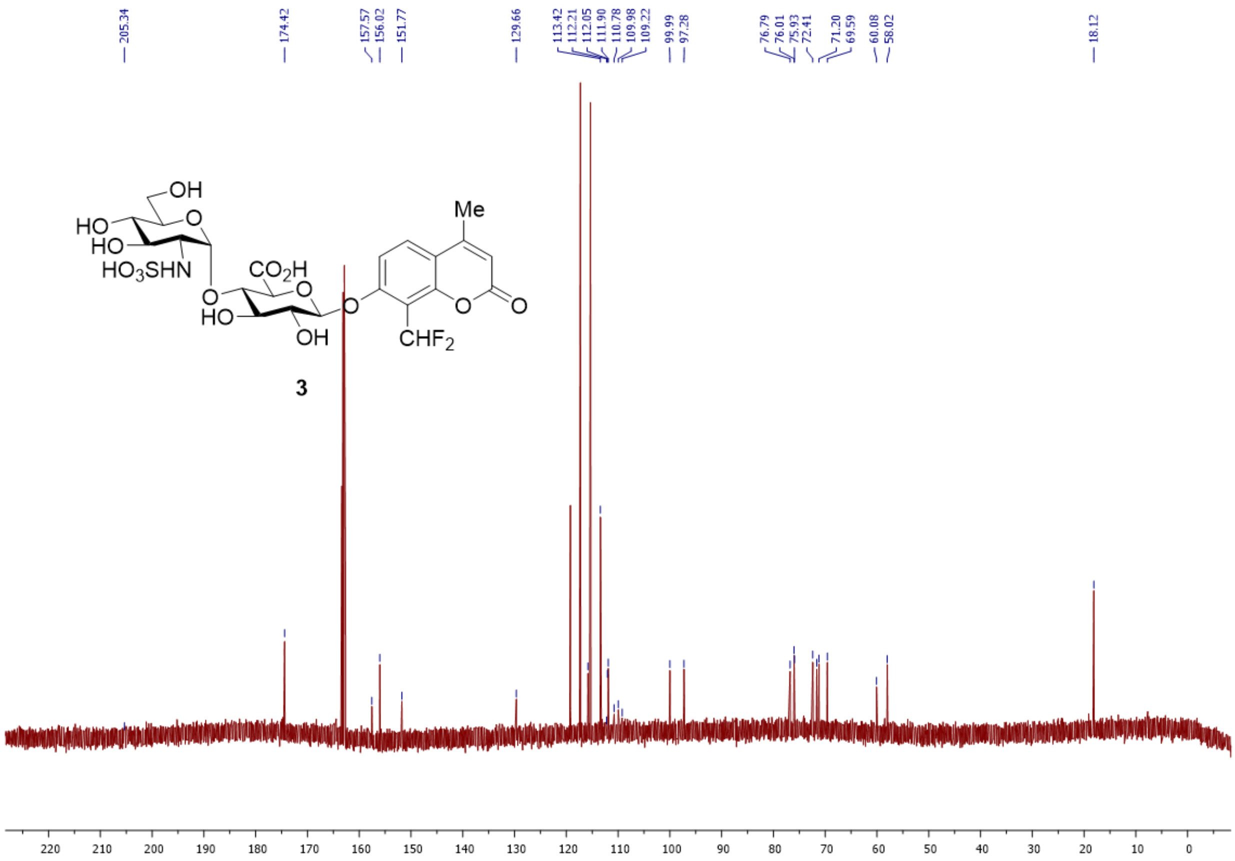

Fig. S32. $^{13}C$ NMR spectrum of **3** in $D_2O$.

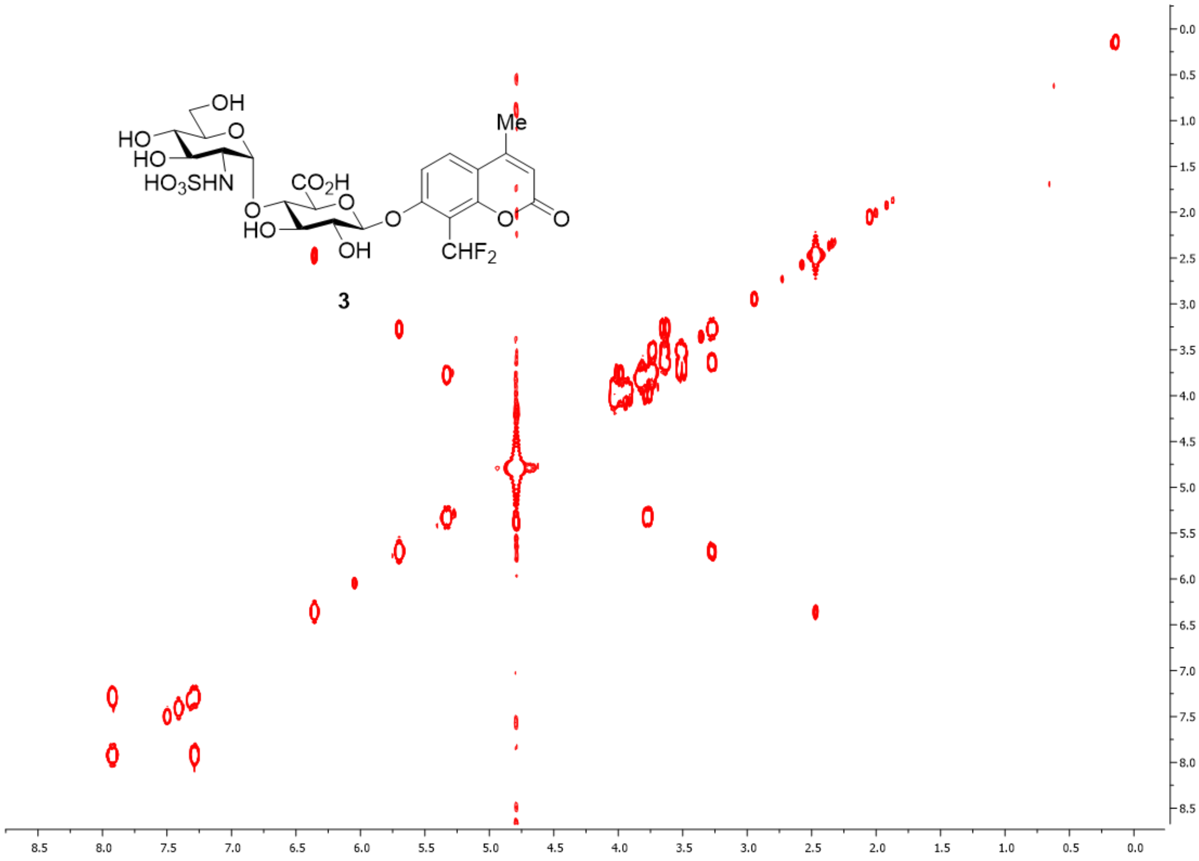


Fig. S33. [1]H-[1]H COSY spectrum of **3** in $D_2O$.

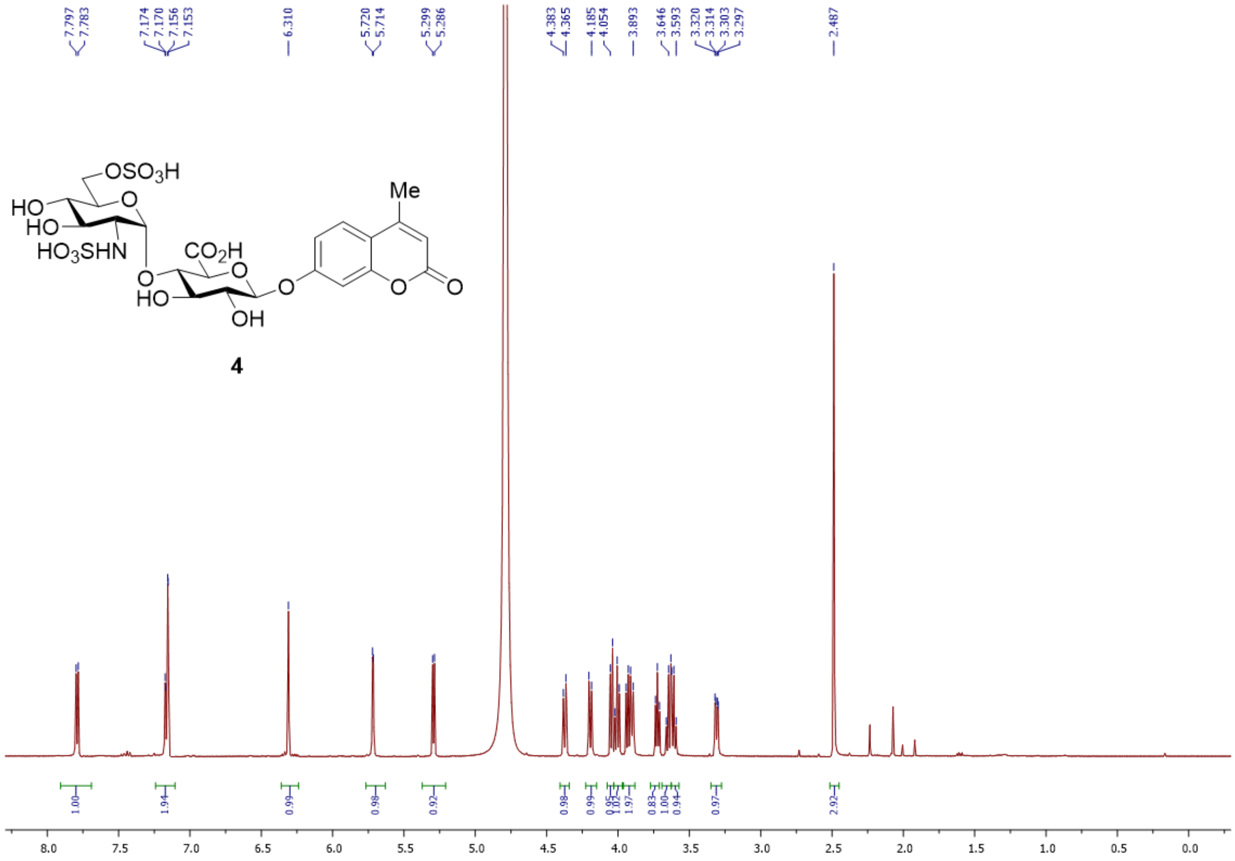

Fig. S34. $^1H$ NMR spectrum of **4** in $D_2O$.

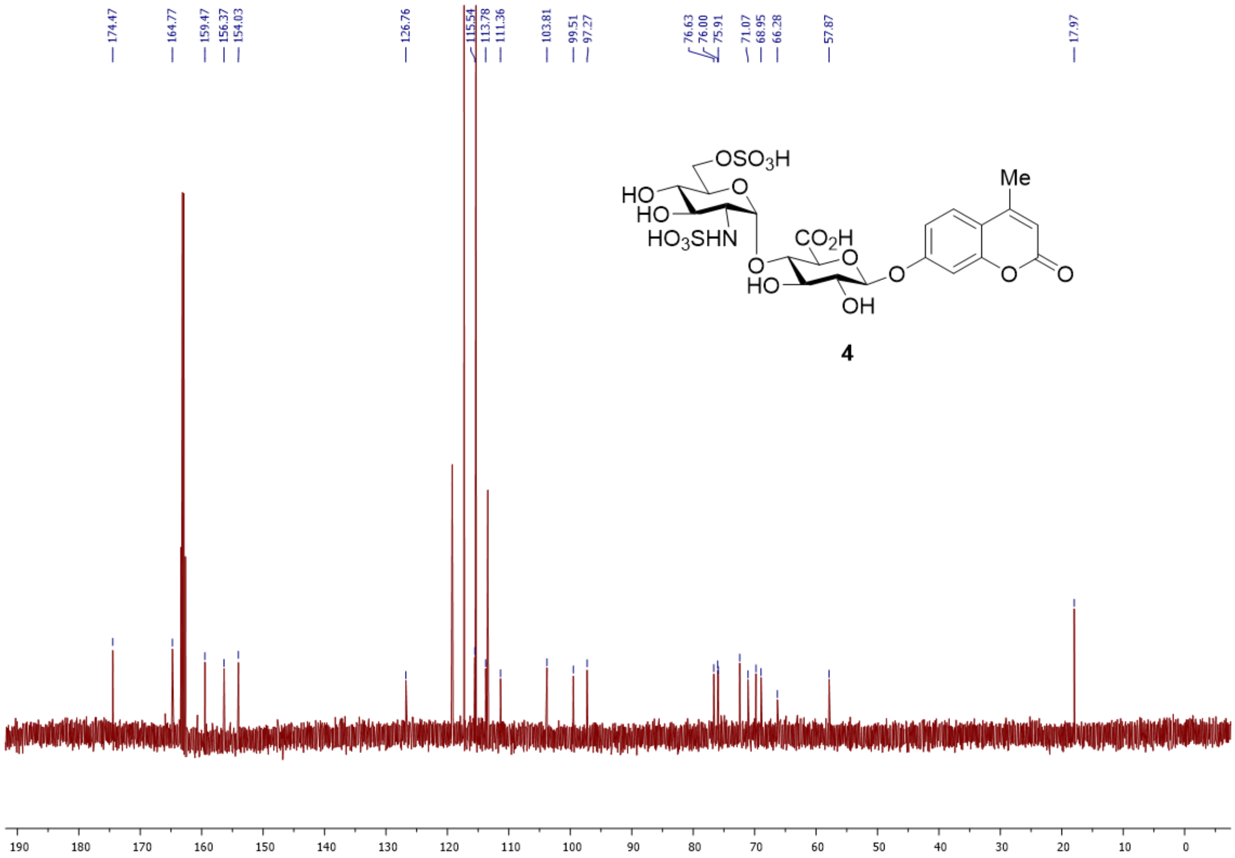


Fig. S35. [13]C NMR spectrum of **4** in $D_2O$.

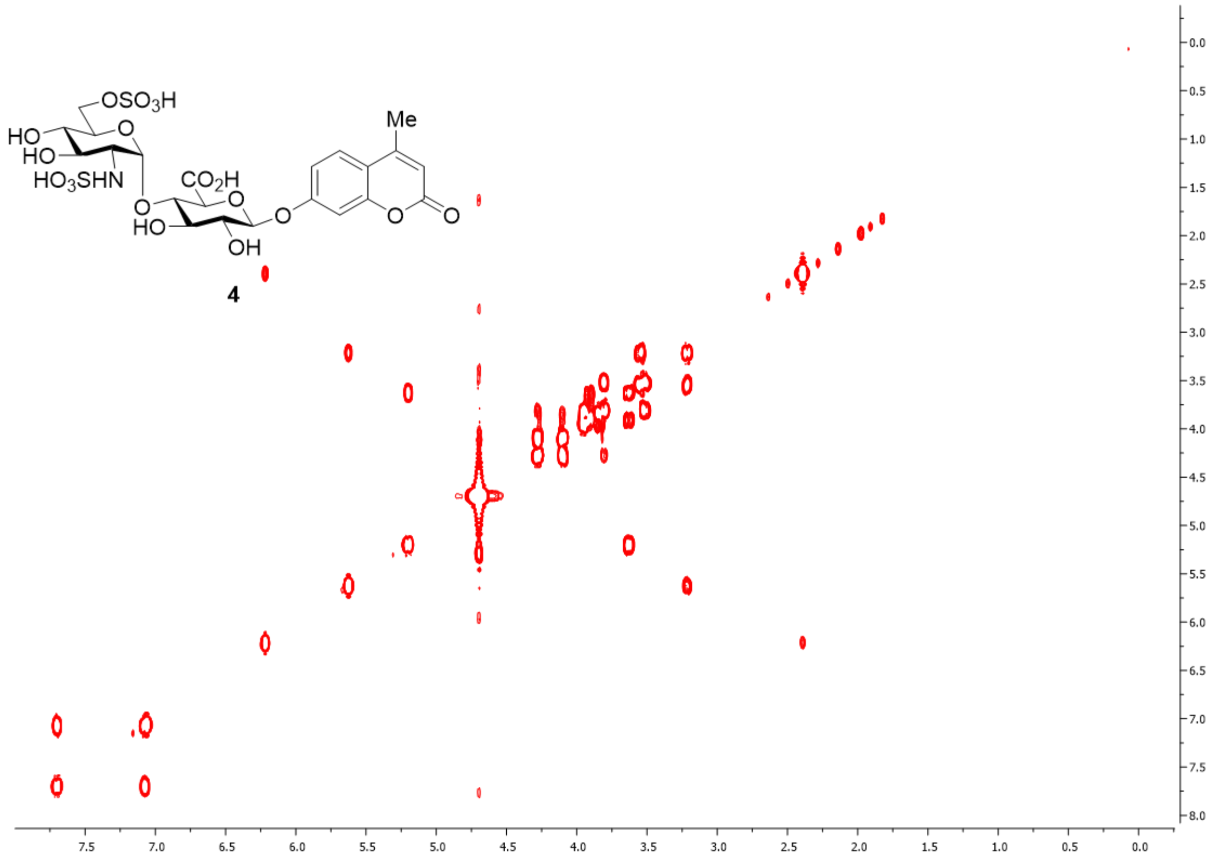

Fig. S36. $^{1}H$-$^{1}H$ COSY spectrum of **4** in $D_2O$.
